# Study design features increase replicability in cross-sectional and longitudinal brain-wide association studies

**DOI:** 10.1101/2023.05.29.542742

**Authors:** Kaidi Kang, Jakob Seidlitz, Richard A.I. Bethlehem, Jiangmei Xiong, Megan T. Jones, Kahini Mehta, Arielle S. Keller, Ran Tao, Anita Randolph, Bart Larsen, Brenden Tervo-Clemmens, Eric Feczko, Oscar Miranda Dominguez, Steve Nelson, Lifespan Brain Chart Consortium, 3R-BRAIN, AIBL, Alzheimer’s Disease Neuroimaging Initiative, Alzheimer’s Disease Repository Without Borders Investigators, CALM Team, CCNP, COBRE, cVEDA, Harvard Aging Brain Study, IMAGEN, POND, The PREVENT-AD Research Group, Jonathan Schildcrout, Damien Fair, Theodore D. Satterthwaite, Aaron Alexander-Bloch, Simon Vandekar

## Abstract

Brain-wide association studies (BWAS) are a fundamental tool in discovering brain-behavior associations. Several recent studies showed that thousands of study participants are required for good replicability of BWAS because the standardized effect sizes (ESs) are much smaller than the reported standardized ESs in smaller studies. Here, we perform analyses and meta-analyses of a robust effect size index using 63 longitudinal and cross-sectional magnetic resonance imaging studies from the Lifespan Brain Chart Consortium (77,695 total scans) to demonstrate that optimizing study design is critical for increasing standardized ESs and replicability in BWAS. A meta-analysis of brain volume associations with age indicates that BWAS with larger variability in covariate have larger reported standardized ES. In addition, the longitudinal studies we examined reported systematically larger standardized ES than cross-sectional studies. Analyzing age effects on global and regional brain measures from the United Kingdom Biobank and the Alzheimer’s Disease Neuroimaging Initiative, we show that modifying longitudinal study design through sampling schemes improves the standardized ESs and replicability. Sampling schemes that improve standardized ESs and replicability include increasing between-subject age variability in the sample and adding a single additional longitudinal measurement per subject. To ensure that our results are generalizable, we further evaluate these longitudinal sampling schemes on cognitive, psychopathology, and demographic associations with structural and functional brain outcome measures in the Adolescent Brain and Cognitive Development dataset. We demonstrate that commonly used longitudinal models can, counterintuitively, reduce standardized ESs and replicability. The benefit of conducting longitudinal studies depends on the strengths of the between-versus within-subject associations of the brain and non-brain measures. Explicitly modeling between-versus within-subject effects avoids averaging the effects and allows optimizing the standardized ESs for each separately. Together, these results provide guidance for study designs that improve the replicability of BWAS.

---

Brain-wide association studies (BWAS) use noninvasive magnetic resonance imaging (MRI) to identify associations between inter-individual differences in behavior, cognition, biological or clinical measurements and brain structure or function^1,2^. A fundamental goal of BWAS is to identify true underlying biological associations that improve our understanding of how brain organization and function are linked to brain health across the lifespan.

Recent studies have raised concerns about the replicability of BWAS^1–3^. Statistical replicability is typically defined as the probability of obtaining consistent results from hypothesis tests across different studies. Like statistical power, replicability is a function of both the standardized effect sizes (ESs) and the sample size^4–6^. Low replicability in BWAS has been attributed to a combination of small sample sizes, small standardized ESs, and bad research practices (such as *p*-hacking and publication bias)^1,2,7–11^. The most obvious solution to increasing the replicability in BWAS is to increase study sample sizes. Several recent studies showed that thousands of study participants are required to obtain replicable findings in BWAS^1,2^. However, massive sample sizes are often infeasible in practice.

Standardized ESs (such as Pearson’s correlation and Cohen’s *d*) are statistical values that not only depend on the underlying biological association in the population, but also on the study design. Two studies of the same biological effect with different study designs will have different standardized ESs. For example, contrasting brain function of depressed versus non-depressed groups will have a different Cohen’s *d* ES if the study design measures more extreme depressed states contemporaneously with measures of brain function, as opposed to less extreme depressed states, even if the underlying biological effect is the same. While researchers cannot increase the magnitude of the underlying biological association, its standardized ES – and thus its replicability – can be increased by critical features of study design.

In this paper, we focus on identifying modifiable study design features that can be used to improve the replicability of BWAS by increasing standardized ESs. Increasing standardized ESs through study design prior to data collection stands in sharp contrast to bad research practices that can artificially inflate reported ESs, such as *p*-hacking and publication bias.

Surprisingly, there has been very little research regarding how modifications to the study design might improve BWAS replicability. Specifically, we focus on two major design features that directly influence standardized ESs: variation in sampling scheme and longitudinal designs^1,12–14^. Notably, these design features can be implemented without inflating the sample estimate of the underlying biological effect when using correctly specified models^15^. By increasing the replicability of BWAS through study design, we can more efficiently utilize the National Institutes of Health’s $1.8 billion average annual investment in neuroimaging research in the past decade^16^.

Here, we conduct a comprehensive investigation of cross-sectional and longitudinal BWAS designs by capitalizing on multiple large-scale data resources. Specifically, we begin by analyzing and meta-analyzing 63 neuroimaging datasets including 77,695 scans from 60,900 cognitively normal (CN) participants from the Lifespan Brain Chart Consortium^17^ (LBCC). We leverage longitudinal data from the UK Biobank (UKB; up to 29,031 scans), the Alzheimer’s Disease Neuroimaging Initiative (ADNI; 2,232 scans), and the Adolescent Brain Cognitive Development study (ABCD; up to 17,210 scans). Within these datasets, we consider many of the most commonly measured brain phenotypes, including multiple measures of brain structure and function. To ensure that our results are broadly generalizable, we evaluate associations with diverse covariates of interest, including age, sex, cognition, and psychopathology. To facilitate comparison between BWAS designs, we introduce a new version of the robust effect size index (RESI) that allows us to demonstrate how longitudinal study design directly impacts standardized ESs. We show that covariate variability has a marked impact on the standardized ES and the replicability of BWAS. Additionally, we find that longitudinal designs are not always beneficial and can in fact reduce standardized ESs in specific cases where careful modeling is required. Critically, the impact of longitudinal data depends directly on the difference in between- and within-subjects effects. Together, our results emphasize that careful study design can markedly improve the replicability of BWAS. To this end, we provide recommendations that allow investigators to optimally design their studies prospectively.

## Meta-analyses show standardized ESs depend on study population and design

To fit each study-level analysis, we regress each of the global brain measures (total gray matter volume (GMV), total subcortical gray matter volume (sGMV), and total white matter volume (WMV), and, mean cortical thickness (CT)) and regional brain measures (regional GMV and CT, based on Desikan-Killiany parcellation^18^) on sex and age in each of the 63 neuroimaging datasets from the LBCC. Age is modeled using nonlinear spline function in linear regression models for the cross-sectional datasets and generalized estimating equations (GEEs) for the longitudinal datasets (Methods). Site effects are removed before the regressions using ComBat^19,20^ (Methods). Analyses for total GMV, total sGMV and total WMV use all 63 neuroimaging datasets (16 longitudinal; Tab. S1). Analyses of regional brain volumes and CT used 43 neuroimaging datasets (13 longitudinal; Methods; Tab. S2).

Throughout the present study, we use the robust effect size index (RESI)^21–23^ as a measure of standardized ES. The RESI is a recently developed index encompassing many types of test statistics and is equal to ½ Cohen’s *d* under some assumptions (Methods; Supplementary Information)^21^. To investigate the effects of study design features on the RESI, we perform meta-analyses for the four global brain measures and two regional brain measures in the LBCC to model the association of study-level design features with standardized ESs.

Study design features are quantified as the sample mean, standard deviation (SD), and skewness of the age covariate as nonlinear terms, and a binary variable indicating the design type (cross-sectional or longitudinal). After obtaining the estimates of the standardized ESs of age and sex in each analysis of the global and regional brain measures, we conduct meta-analyses of the estimated standardized ESs using weighted linear regression models with study design features as covariates (Methods).

For total GMV, the partial regression plots of the effect of each study design feature demonstrate a strong cubic-shape relationship between the standardized ES for total GMV-age association and study population mean age. This cubic shape indicates that the strength of the age effect varies with respect to the age of the population being studied. The largest age effect on total GMV in the human lifespan occurs during early and late adulthood (Tab. S3; Fig. 1a). There is also a strong positive linear effect of the study population SD of age and the standardized ES for total GMV-age association. For each unit increase in the SD of age (in years), expected standardized ES increases by about 0.1 (Fig. 1a). This aligns with the well-known relationship between correlation strength and covariate SD indicated by statistical principles^24^. Plots for total sGMV, total WMV, and mean CT show U-shaped changes of the age effect with respect to study population mean age (Fig. 1b-d). A similar but sometimes weaker relationship is shown between expected standardized ES and study population SD and skewness of the age covariate (Fig. 1b-d; Tab. S4-S6). Finally, the meta-analyses also show a moderate effect of study design on the standardized ES of age on each of the global brain measures (Fig. 1a-d; Tab. S3-S6). The average standardized ES for total GMV-age associations in longitudinal studies (RESI=0.39) is substantially larger than in cross-sectional studies (RESI=0.08) after controlling for the study design variables, corresponding to a >380% increase in the standardized ES for longitudinal studies. This value quantifies the systematic differences in the standardized ESs between the cross-sectional and longitudinal studies among the 63 neuroimaging studies. Notably, longitudinal study design does not improve the standardized ES for biological sex, because sex does not vary within subjects in these studies (Tab. S7-S10; Fig. S2).

**Fig. 1.**
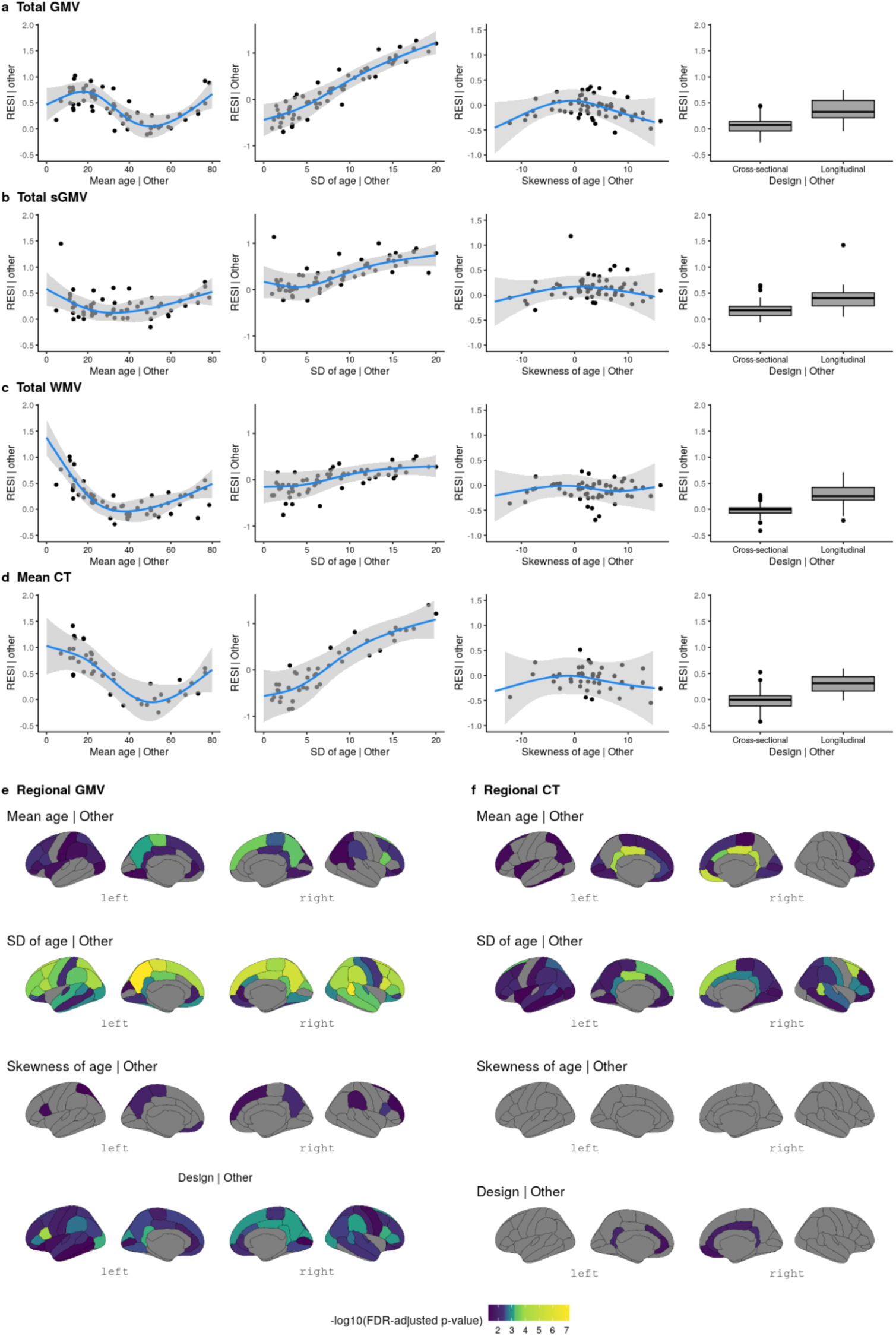
Meta-analyses reveal study design features that are associated with larger standardized effect sizes (ESs) of age on different brain measures. (a-d) Partial regression plots of the meta-analyses of standardized ESs (RESI) for the association between age and global brain measures (a) total gray matter volume (GMV), (b) total subcortical gray matter volume (sGMV), (c) total white matter volume (WMV), and (d) mean cortical thickness (CT) (“| Other” means after fixing the other features at constant levels: design = cross-sectional, mean age = 45 years, sample age SD = 7 and/or skewness of age = 0 (symmetric)) show that standardized ESs vary with respect to the mean and SD of age in each study. The blue curves are the expected standardized ESs for age from the locally estimated scatterplot smoothing (LOESS) curves. The gray areas are the 95% confidence bands from the LOESS curves. (e-f) The effects of study design features on the standardized ESs for the associations between age and regional brain measures (regional GMV and CT). Regions with Benjamini-Hochberg adjusted *p*-values<0.05 are shown in color.

For regional GMV and CT, similar effects of study design features also occur across regions (Fig. 1e-f; 34 regions per hemisphere). In most of the regions, the standardized ESs of age on regional GMV and CT are strongly associated with the study population SD of age.

Longitudinal study designs generally tend to have a positive effect on the standardized ESs for regional GMV-age associations and a positive, but weaker effect on the standardized ESs for regional CT-age associations (see Extended Display Item). To improve the comparability of standardized ESs between cross-sectional and longitudinal studies, we propose a new ES index: the cross-sectional RESI for longitudinal datasets (Supplementary Information: Section 1). The cross-sectional RESI for longitudinal datasets represents the RESI in the same study population, if the longitudinal study had been conducted cross-sectionally. This newly developed ES index allows us to quantify the benefit of using a longitudinal study design in a single dataset (Supplementary Information: Section 1.3).

The meta-analysis results demonstrate that standardized ESs are dependent on study design features, such as mean age, SD of the age of the sample population, and cross-sectional/longitudinal design. Moreover, the results suggest that modifying study design features, such as increasing variability and conducting longitudinal studies, can increase the standardized ESs in BWAS.

## Improved sampling schemes can increase standardized ESs and replicability

To investigate the impact of modifying the variability of the age covariate on increasing standardized ESs and replicability, we implement three sampling schemes that produce different sample SD of the age covariate. We treat the large-scale cross-sectional United Kingdom Biobank (UKB) data as the population and draw samples whose age distributions follow a pre-specified shape (bell-shaped, uniform, and U-shaped; Fig. S3; Methods). In UKB, the U-shaped sampling scheme on age increases the standardized ES for the total GMV-age association by 60% compared to bell-shaped and by 27% compared to uniform (Fig. 2a), with an associated increase in replicability (Fig. 2b). To achieve 80% replicability for detecting the total GMV-age association (Methods), <100 subjects are sufficient if using the U-shaped sampling scheme, whereas about 200 subjects are needed if the bell-shaped sampling scheme is used (Fig. 2b). A similar pattern can be seen for the regional outcomes of GMV and CT (Fig. 2c-f). The U-shaped sampling scheme typically provides the largest standardized ESs of age and the highest replicability, followed by the uniform and bell-shaped schemes. The U-shaped sampling scheme shows greater region-specific improvement in the standardized ESs for regional GMV-age and regional CT-age associations and replicability compared to the bell-shaped scheme (Fig. S4).

**Fig. 2.**
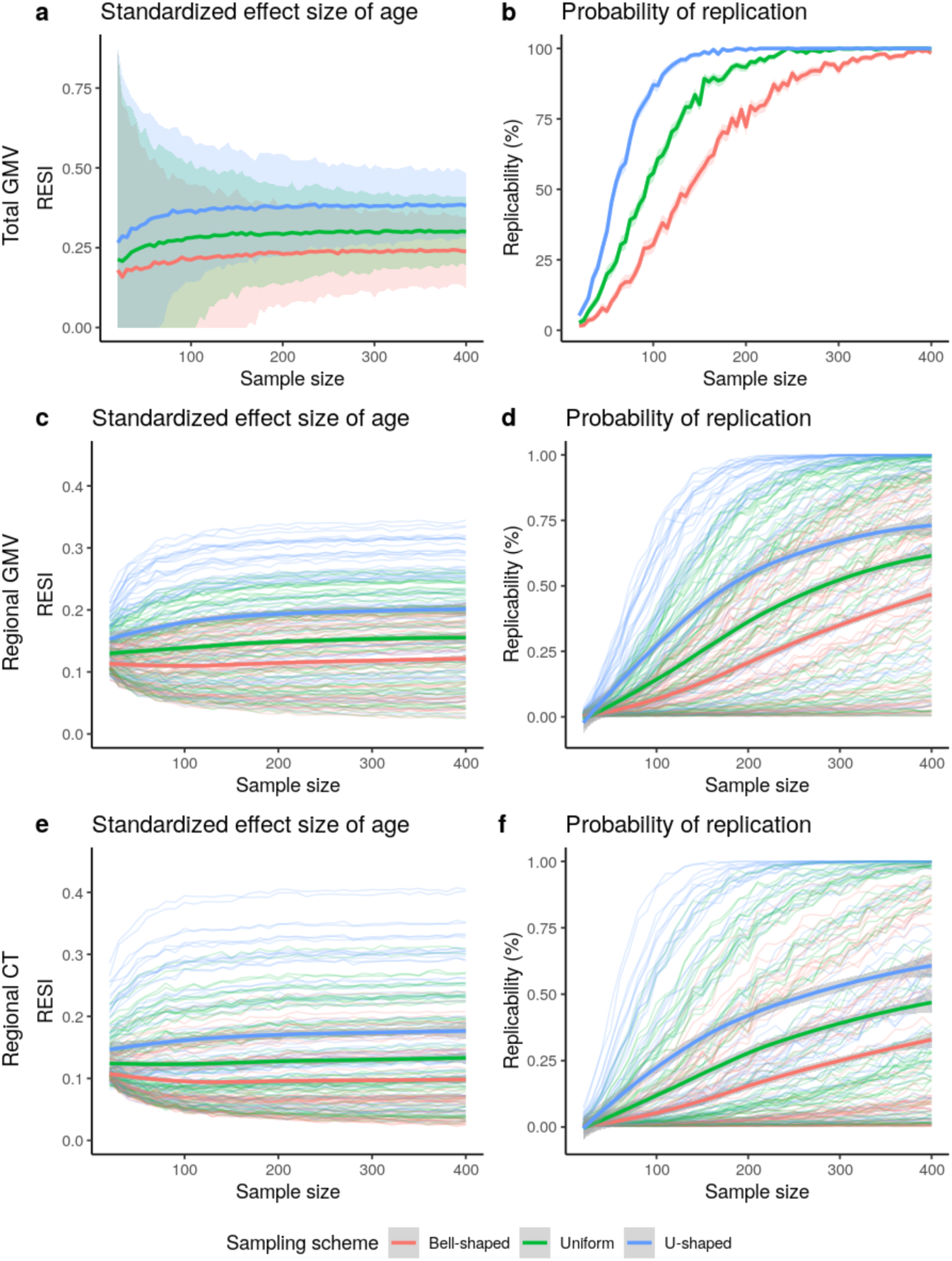
Increased standardized effect sizes (ESs) and replicability for age associations with different brain measures: (a-b) total gray matter volume (GMV), (c-d) regional GMV, and (e-f) regional cortical thickness (CT), under three sampling schemes in the UKB study (*N* = 29,031 for total GMV and *N* = 29,030 for regional GMV and CT). The sampling schemes target different age distributions to increase the variability of age: Bell-shaped<Uniform<U-shaped (Fig. S3). Using the resampling schemes that increase age variability increases (**a**) the standardized ESs and (**b**) replicability (at significance level of 0.05) for total GMV-age association. The same result holds for (**c-d**) regional GMV and (**e-f**) regional CT. The curves represent the average standardized ES or estimated replicability at a given sample size and sampling scheme. The shaded areas represent the corresponding 95% confidence bands. (**c-f**) The bold curves are the average standardized ES or replicability across all regions with significant uncorrected effects using the full UKB data.

To investigate the effect of increasing the variability of the age covariate longitudinally, we implement sampling schemes to adjust the between- and within-subject variability of age in the bootstrap samples from the longitudinal ADNI dataset. In the bootstrap samples, each subject has two measurements (baseline and a follow-up). To imitate the true operation of a study, we select each subject’s two measurements based on baseline age and the follow-up age by targeting specific distributions for the baseline age and the age change at the follow-up time point (Fig. S5; Methods). Increasing between-subject and within-subject variability of age increases the average observed standardized ESs, with corresponding increases in replicability (Fig. 3). U-shaped between-subject sampling scheme on age increases the standardized ES for total GMV-age association by 23.6% compared to bell-shaped and by 12.1% compared to uniform, when using uniform within-subject sampling scheme (Fig. 3a).

**Fig. 3.**
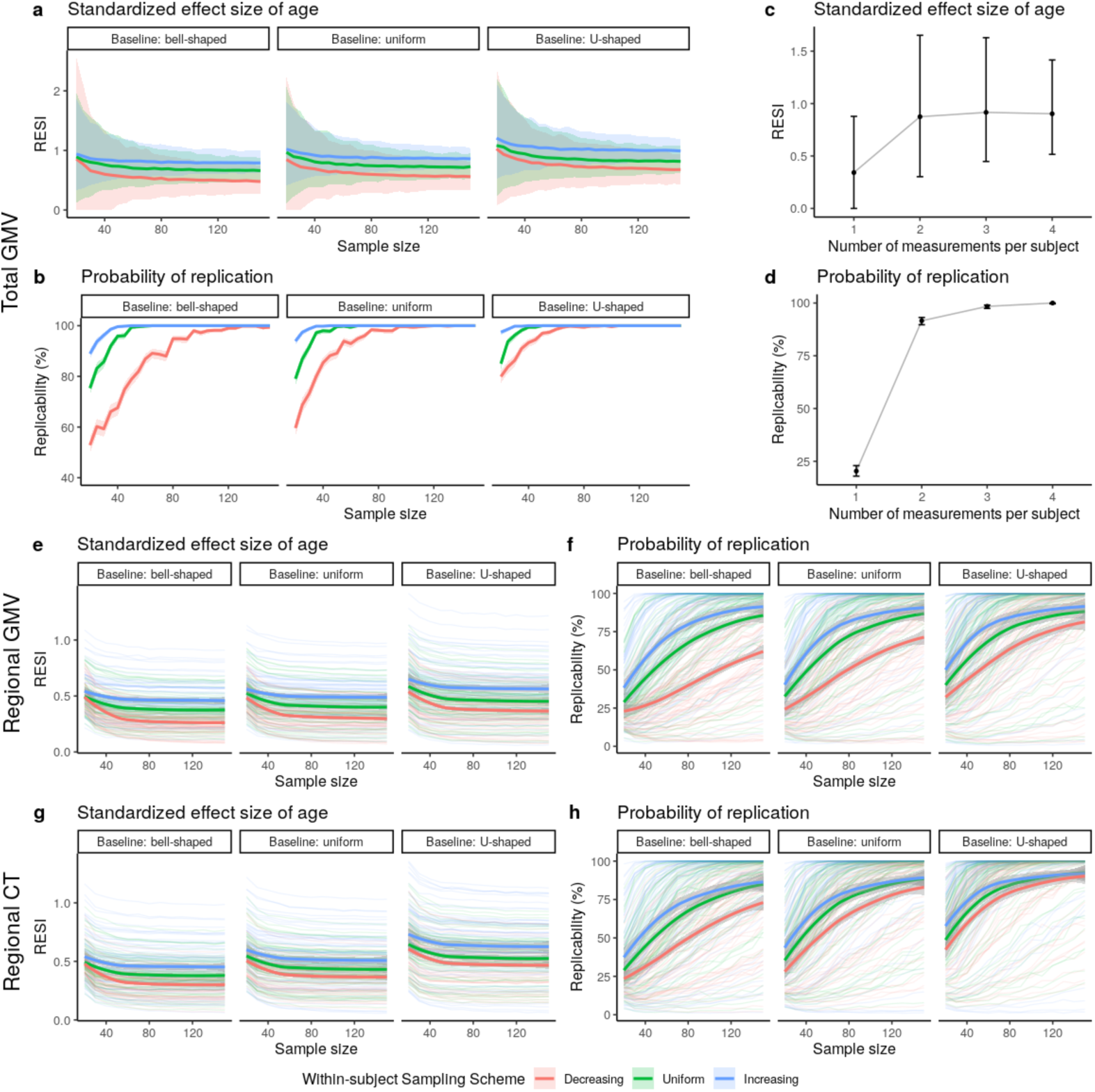
Increased standardized effect sizes (ESs) and replicability for age associations with structural brain measures under different longitudinal sampling schemes in the ADNI data. Three different sampling schemes (Fig. S5) are implemented in bootstrap analyses to modify the between- and within-subject variability of age, respectively. (**a-b**) Implementing the sampling schemes results in higher (between- and/or within-subject) variability and increases the (**a**) standardized ES and (**b**) replicability (at significance level of 0.05) for the total gray matter volume (GMV)-age association. The curves represent the average standardized ES or estimated replicability and the gray areas are the 95% confidence bands across the 1,000 bootstraps. (**c-d**) Increasing the number of measurements from one to two per subject provides the most benefit on (**c**) standardized ES and (**d**) replicability for the total GMV-age association when using uniform between- and within-subject sampling schemes and *N* = 30. The points represent the mean standardized ESs or estimated replicability and the whiskers are the 95% confidence intervals. (**e-h**) Increased standardized ESs and replicability for the associations of age with regional (**e-f**) GMV and (**g-h**) cortical thickness (CT), across all brain regions under different sampling schemes. The bold curves are the average standardized ES or estimated replicability across all regions with significant uncorrected effects using the full ADNI data. The gray areas are the corresponding 95% confidence bands. Increasing the between- and within-subject variability of age by implementing different sampling schemes can increase the standardized ESs of age, and the associated replicability, on regional GMV and regional CT.

Additionally, we investigate the effect of the number of measurements per subject on the standardized ES and replicability in longitudinal data using the ADNI dataset. Adding a single additional measurement after the baseline increases the standardized ES for total GMV-age association by 156% and replicability by 350%. The benefit of additional measurements is minimal (Fig. 3c-d). Finally, we also evaluate the effects of the longitudinal sampling schemes on regional GMV and CT in ADNI (Fig. 3e-h). When sampling two measurements per subject, the between- and within-subject sampling schemes producing larger age variability increase the standardized ES and replicability across all regions.

Together, these results suggest having larger spacing in between- and within-subject age measurements increases standardized ES and replicability. Most of the benefit of the number of within-subject measurements is due to the first additional measurement after baseline.

## Preferred sampling schemes and longitudinal designs depend on the between- and within-subject associations of the brain and non-brain measures

To explore if the proposed sampling schemes are effective on a variety of non-brain covariates and their associations with structural and functional brain measures, we implement the sampling schemes on all subjects (with and without neuropsychiatric symptoms) with cross-sectional and longitudinal measurements from the ABCD dataset. The non-brain covariates include the NIH toolbox^25^, Child Behavior Checklist (CBCL), body mass index (BMI), birth weight, and handedness (Tab. S11-S12; Methods). Functional connectivity (FC) is used as a functional brain measure and is computed for all pairs of regions in the Gordon atlas^26^ (Methods). We use the bell- and U-shaped target sampling distributions to control the between- and within-subject variability of each non-brain covariate (Methods). For each non-brain covariate, we show the results for the four combinations of between- and within-subject sampling schemes. Overall, there is a consistent benefit to increasing between-subject variability of the covariate (Fig. S6), which leads to >1.8 factor reduction in sample size needed to scan for 80% replicability and >1.4 factor increase in the standardized ES for over 50% of associations. Moreover, 72% of covariate-outcome associations had increased standardized ESs by increasing the between-subject variability of the covariates (Fig. S7).

Importantly, increasing *within*-subject variability decreases the standardized ESs for many structural associations (Fig. 4a-f; Fig. S6a-f), suggesting that conducting longitudinal analyses can result in decreased replicability compared to cross-sectional analyses. For the FC outcomes, there is a slight positive effect of increasing within-subject variability (Fig. 4g; Fig. S6g). To evaluate the lower replicability of the structural associations with increasing within-subject variability, we compare cross-sectional standardized ESs of the non-brain covariates on each brain measure using the baseline measurements to the standardized ESs estimated using the full longitudinal data (Fig. 5a-d). Consistent with the reduction in standardized ES by increasing within-subject variability, for most structural associations (GMV and CT), conducting cross-sectional analyses using the baseline measurements results in larger standardized ESs (and higher replicability) than conducting analyses using the full longitudinal data. This finding is consistent when fitting a cross-sectional model using the 2-year follow-up measurement (Fig. S8). Identical results are found using linear mixed models (LMMs) with individual-specific random intercepts, which are commonly used in BWAS (Fig. S9). Together, these results suggest that the benefit of conducting longitudinal studies and increasing within-subject variability is highly dependent on the brain-behavior association, and counterintuitively, longitudinal designs can reduce the standardized ESs and replicability (unless the models are properly specified - see below).

**Fig. 4.**
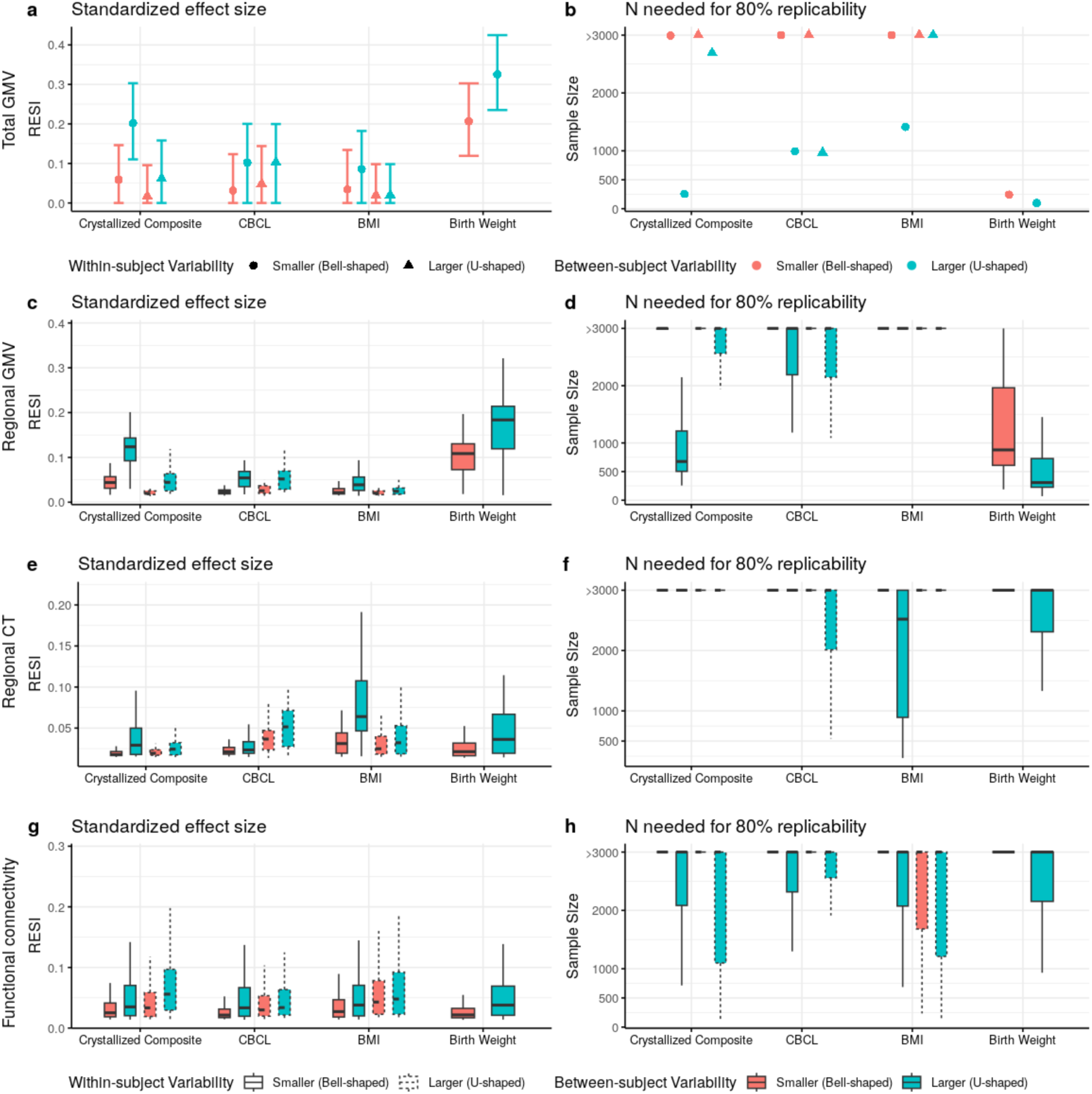
Heterogeneous improvement of standardized effect sizes (ESs) for select cognitive, mental health, and demographic associations with structural and functional brain measures in the ABCD study with bootstrapped samples of *N* = 500. (**a**) U-shaped between-subject sampling scheme (blue) that increases between-subject variability of the non-brain covariate produces larger standardized ESs and (**b**) reduces the number of participants scanned to obtain 80% replicability in total gray matter volume (GMV). The points and triangles are the average standardized ESs across bootstraps and the whiskers are the 95% confidence intervals. Increasing within-subject sampling (triangles) can reduce standardized ESs. A similar pattern holds in (**c-d**) regional GMV and (**e-f**) regional cortical thickness (CT); boxplots show the distributions of the standardized ESs across regions (or region pairs for functional connectivity (FC)). In contrast, (**g**) regional pairwise FC standardized ESs improve by increasing between-(blue) and within-subject variability (dashed borders) with a corresponding reduction in the (**h**) number of participants scanned for 80% replicability. See Fig. S6 for the results for all non-brain covariates examined.

**Fig. 5.**
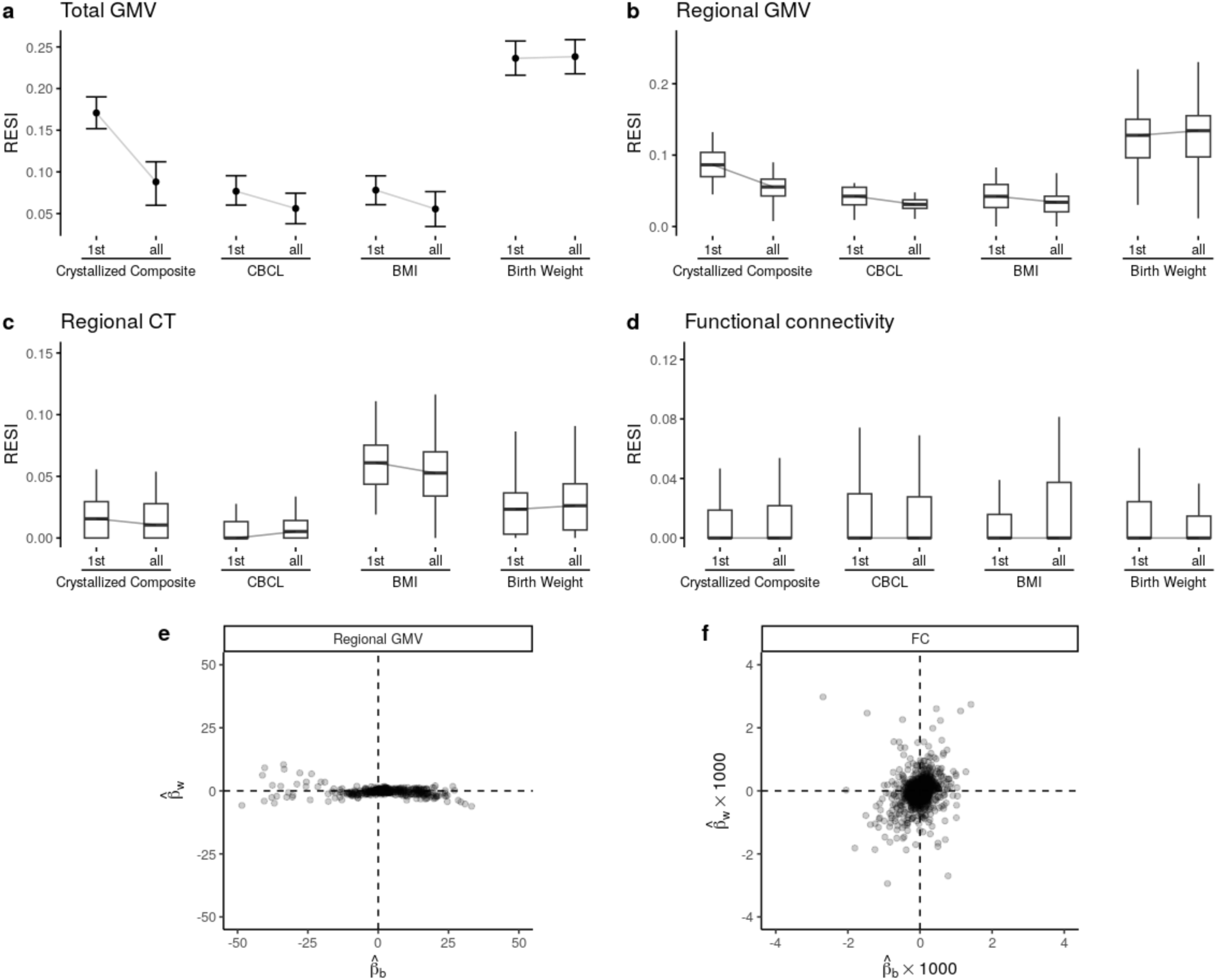
Longitudinal study designs can reduce standardized effect sizes (ESs) and replicability due to differences in between-versus within-subject associations of brain and behavior measures. Boxplots show the distributions of the standardized ESs across regions. (**a-c**) Cross-sectional analyses (using only the baseline measures; indicated by “1st”s on the x-axes) can have larger standardized ESs than the same longitudinal analyses (using the full longitudinal data; indicated by “all”s on the x-axes) for structural brain measures in ABCD. (**d**) The functional connectivity (FC) measures do not show such a reduction of standardized ESs in longitudinal modeling. See Fig. S8 for the results for all non-brain covariates examined. (**e**) Most regional gray matter volume (GMV) associations (Fig. 5c) have larger between-subject parameter estimates (*β*_*b*_, x-axis) than within-subject parameter estimates (*β_w_*, y-axis; see Supplementary Information: Eqn (13)), whereas (**f**) FC associations (Fig. 5g) show less heterogeneous relationships between the two parameters.

To investigate why increasing within-subject variability or using longitudinal designs is not beneficial for some associations, we examined an assumption common to widely used GEEs and LMMs in BWAS. Widely used models assume that there is consistent association strength between the brain measure and non-brain covariate across between- and within-subject changes in the non-brain covariate. However, the between- and within-subject association strengths can differ because non-brain measures can be more variable than structural brain measures for a variety of reasons. For example, crystallized composite scores may vary within a subject longitudinally because of time-of-day effects, lack of sleep, or natural noise in the measurement. In contrast, GMV is more precise and it is not vulnerable to other sources of variability that might accompany the crystallized composite score. This combination leads to low within-subject association between these variables (Tab. S13). FC measures are more similar to crystallized composite scores in that they are subject to higher within-subject variability and natural noise, so they have a higher potential for stronger within-subject associations with crystallized composite scores (i.e., they are more likely to vary together based on time of day, lack of sleep, etc). To demonstrate this, we fit models that estimate distinct effects for between- and within-subject associations in the ABCD (Methods) and find that there are large between-subject parameter estimates and small within-subject parameter estimates in total and regional GMV (Supplementary Information: Section 3.2; Tab. S13; Fig. 5e), whereas the FC associations are distributed more evenly across between- and within-subject parameters (Fig. 5f). If the between- and within-subject associations are different, these widely used longitudinal models average the two associations (Supplementary Information: Section 3 Eqn. (14)). Fitting these associations separately avoids averaging the larger effect with the smaller effect and can inform our understanding of brain-behavior associations (Supplementary Information: Section 3.2). This approach ameliorates the reduction in standardized ESs caused by longitudinal designs for structural brain measures in the ABCD (Fig. S10-S11; Supplementary Information: Section 3). This longitudinal model has a similar between-subject standardized ES to the cross-sectional model (Methods: *Estimation of the between- and within-subject effects*; Fig. S11). In short, longitudinal designs can be detrimental to replicability when the between- and within-subject effects differ and the model is incorrectly specified.

## Discussion

BWAS are a fundamental tool to discover brain-behavior associations that may inform our understanding of brain health across the lifespan. With increasing evidence of small standardized ESs and low replicability in BWAS, optimizing study design to increase standardized ESs and replicability is a critical prerequisite for progress^1,27^. We show the influence of study design features on standardized ESs and replicability using diverse datasets, neuroimaging features, and covariates of interest. Our results indicate that increasing between-subject variability of the covariate systematically increases standardized ESs and replicability. Furthermore, we demonstrate that the benefit of performing a longitudinal study strongly depends on the nature of the association and the model used to fit the data. We use statistical principles to show that typical longitudinal models used in BWAS average between- and within-subject effects. When we model between- and within-subject effects separately, we can increase the standardized ESs for the between- and within-subject effects separately using the modified sampling schemes and retain similar or slightly higher replicability compared to the cross-sectional model (Fig. S10-S11). While striving for large sample sizes remains important when designing a study, our findings emphasize the importance of considering other design features to improve standardized ESs and replicability of BWAS.

### Sampling strategies can increase replicability

Our results demonstrate that standardized ES and replicability can be increased by enriched sampling of subjects with small and large values of the covariate of interest. This is well-known in linear models where the standardized ES is explicitly a function of the SD of the covariate^24^. We show that designing a study to have larger covariate SD increases standardized ESs by a median factor of 1.4, even when there is nonlinearity in the association, such as with age and GMV (Fig. S1). When the association is very non-monotonic – as in the case of a U-shape relationship between covariate and outcome – sampling the tails more heavily could decrease replicability, and diminish our ability to detect nonlinearities in the center of the study population. In such a case, sampling to obtain a uniform distribution of the covariate balances power across the range of the covariate and can increase replicability relative to random sampling when the covariate has a normal distribution in the population. Increasing between-subject variability is beneficial in more than 72% of the association pairs we studied, despite the presence of such nonlinearities (Fig. S7).

Because standardized ESs are dependent on study design, careful design choices can simultaneously increase standardized ESs and study replicability. Two-phase, extreme group, and outcome-dependent sampling designs can inform which subjects should be selected for imaging from a larger sample in order to increase the efficiency and standardized ESs of brain-behavior associations^28–34^. For example, given the high degree of accessibility of cognitive and behavioral testing (e.g., to be performed virtually or electronically), individuals scoring at the extremes on a testing scale/battery (“phase I”) could be prioritized for subsequent brain scanning (“phase II”). When there are multiple covariates of interest, multivariate two-phase designs can be used to increase standardized ESs and replicability^35^. Multivariate designs are also needed to stratify sampling to avoid confounding by other socio-demographic variables.

Together, the use of optimal designs can increase both standardized ESs and replicability relative to a design that uses random sampling^32^. If desired, weighted regression (such as inverse probability weighting) can be combined with optimized designs to estimate a standardized ES that is consistent with the standardized ES if the study had been conducted in the full population^35–37^. Choosing highly reliable psychometric measurements or interventions (e.g., medications or neuromodulation within a clinical trial)^38–40^ may also be effective for increasing replicability. The decision to pursue an optimized design will also depend on other practical factors, such as the cost and complexity of acquiring other (non-imaging) measures of interest and the specific translational goals of the research.

### Longitudinal design considerations

In the meta-analysis, longitudinal studies of the total GMV-age associations have, on average, >380% larger standardized ESs than cross-sectional studies. However, we see in subsequent analyses that the benefit of conducting a longitudinal design is highly dependent on both the between- and the within-subject effects. When the between- and the within-subject effect parameters are equal and the within-subject brain measure error is low, longitudinal studies offer larger standardized ESs than cross-sectional studies (Supplementary Information: Section 3.1)^41^. This combination of equal between- and within-subject effects and low within-subject measurement error is the reason there is a benefit of longitudinal design in ADNI for the total GMV-age association (Fig. S12). While the standardized ES in these analyses did not account for cost per measurement, comparing efficiency per measurement supports the approach of collecting two measurements per subject in this scenario (Supplementary Information: Section 3.1).

Longitudinal models offer the unique ability to separately estimate between- and within-subject effects. When the between- and within-subject effects differ but we still fit them with a single effect, we mistakenly assume they are equal and the interpretation of that coefficient becomes complicated. The effect becomes a weighted average of the between- and within-subject effects whose weights are determined by the study design features (Supplementary Information: Section 3). The apparent lack of benefit of longitudinal designs in the ABCD on the study of GMV associations is because within-subject changes in the non-brain measures are not associated with within-subject changes in GMV (Fig. 5e; Tab. S13). The smaller standardized ESs we find in longitudinal analyses are due to the contribution from the smaller within-subject effect to the weighted average of the between- and within-subject effects (Supplementary Information: Section 3 Eqn. (14)).

We provide recommendations for longitudinal design and analysis strategies (Fig. S13-S14). Briefly, fitting the between- and within-subject effects separately prevents averaging the two effects (Supplementary Information: Section 3.2). These two effects are often not directly comparable to the effect obtained from a cross-sectional model because they have different interpretations (Supplementary Information: Section 3.2)^42–44^. Using sampling strategies to increase between- and within-subject variability of the covariate will respectively increase the standardized ESs for between- and within-subject associations. Thus, longitudinal designs can be helpful and optimal even when the between- and within-subject effects differ, if modeled correctly.

## Conclusions

While it is difficult to provide universal recommendations for study design and analysis, the present study provides general guidelines for designing and analyzing BWAS for optimal standardized ESs and replicability based on both empirical and theoretical results (Fig. S13-S14). Although the decision for a particular design or analysis strategy may depend on unknown features of the brain and non-brain measures and their association, these characteristics can be evaluated in pilot data or the analysis dataset (Fig. S12; Supplementary Information: Section 3.2). One general principle that increases standardized ESs for most associations is to increase the covariate SD (through, e.g., two-phase, extreme group, and outcome-dependent sampling), which is practically applicable to a wide range of BWAS contexts. Moreover, longitudinal designs can provide larger standardized ES and higher replicability, but the benefit is dependent on the form of the association between the covariate and outcome. Longitudinal BWAS enable us to study between- and within-subject effects, and they should be used when the two effects are hypothesized to be different. Together, our findings highlight the importance of rigorous evaluation of study designs and statistical analysis methods that can improve the replicability of BWAS.

## Supporting information

Extended Display Item

## Methods

### LBCC dataset and processing

The original LBCC dataset includes 123,984 MRI scans from 101,457 human participants across more than 100 studies (which include multiple publicly available datasets^45–55^) and is described in our prior work^17^ (see Supplemental Information and Tab. S1 from prior study). We filtered to the subset of cognitively normal (CN) participants whose data were processed using FreeSurfer version 6.1. Studies were curated for the analysis by excluding duplicated observations and studies with fewer than 4 unique age points, sample size less than 20, and/or only participants of one sex. If there were fewer than three participants having longitudinal observations, only the baseline observations were included and the study was considered cross-sectional. If a subject had changing demographic information during the longitudinal follow-up (e.g., changing biological sex), only the most recent observation was included. We updated the LBCC dataset with the ABCD release 5 resulting in a final dataset that includes 77,695 MRI scans from 60,900 CN participants with available total GMV, sGMV, and GMV measures across 63 studies (Tab. S1). In this dataset, 74,148 MRI scans from 57,538 participants across 43 studies have complete-case regional brain measures (regional GMV, regional surface area, and regional cortical thickness (CT), based on Desikan-Killiany parcellation^18^) (Tab. S2). The global brain measure mean CT was derived using the regional brain measures (see below).

#### Structural brain measures

Details of data processing are described in our prior work^17^. Briefly, total GMV, sGMV, and WMV were estimated from T1-weighted and T2-weighted (when available) MRIs using the “aseg” output from Freesurfer 6.0.1. All three cerebrum tissue volumes were extracted from the aseg.stats files output by the recon-all process: ’Total cortical gray matter volume’ for GMV; ’Total cerebral white matter volume’ for WMV; and ‘Subcortical gray matter volume’ for sGMV (inclusive of thalamus, caudate nucleus, putamen, pallidum, hippocampus, amygdala, and nucleus accumbens area; https://freesurfer.net/fswiki/SubcorticalSegmentation). Regional GMV and CT across 68 regions (34 per hemisphere, based on Desikan-Killiany parcellation^18^) were obtained from the aparc.stats files output by the recon-all process. Mean CT across the whole brain is the weighted average of the regional CT weighted by the corresponding regional surface areas.

### Preprocessing specific to ABCD

#### Functional connectivity measures

Longitudinal functional connectivity (FC) measures were obtained from the ABCD-BIDS community collection, which houses a community-shared and continually updated ABCD neuroimaging dataset available under Brain Imaging Data Structure (BIDS) standards. The data used in these analyses were processed using the abcd-hcp-pipeline version 0.1.3, an updated version of The Human Connectome Project MRI pipeline^56^. Briefly, resting state fMRI time series were demeaned and detrended, and a GLM was used to regress out mean WM, CSF, and global signal as well as motion variables and then band-pass filtered. High motion frames (filtered-FD > 0.2mm) were censored during the demeaning and detrending. After preprocessing the time series were parcellated using the 352 regions of the Gordon atlas (including 19 subcortical structures) pairwise Pearson correlations were computed among the regions. FC measures were estimated from resting state fMRI time series using a minimum of 5 minutes of data. After Fisher’s z-transformation, the connectivities were averaged across the 24 canonical functional networks^26^, forming 276 inter-network connectivities and 24 intra-network connectivities.

#### Cognitive and other covariates

The ABCD dataset is a large-scale repository aiming to track the brain and psychological development of over 10,000 children ages 9 to 16 by measuring hundreds of variables including demographic, physical, cognitive, and mental health variables^57^. We used release 5 of ABCD study to examine the effect of the sampling schemes on other types of covariates including cognition (fully corrected *T*-scores of the individual subscales and total composite scores of NIH Toolbox^25^), mental health (total problem CBCL syndrome scale), and other common demographic variables (body mass index (BMI), birth weight, and handedness). For each of the covariates, we evaluated the effect of the sampling schemes on their associations with the global and regional structural brain measures and FC after controlling for nonlinear age and sex (and, for FC outcomes only, mean frame displacement (FD)).

For the analyses of structural brain measures, there were 3 non-brain covariates with fewer than 5% non-missing follow-ups at both 2-year and 4-year follow-ups (i.e., Dimensional Change Card Sort Test, Cognition Fluid Composite, and Cognition Total Composite Score; Tab. S11), and only their baseline cognitive measurements were included in the analyses. For the remaining 11 variables (i.e., Picture Vocabulary Test, Flanker Inhibitory Control and Attention Test, List Sorting Working Memory Test, Pattern Comparison Processing Speed Test, Picture Sequence Memory Test, Oral Reading Recognition Test, Crystallized Composite, CBCL, birth weight, BMI and handedness), all of the available baseline, 2-year and 4-year follow-up observations were used. For the analyses of the FC, only the baseline observations for List Sorting Working Memory Test were used due to missingness (Tab. S12).

The records with BMI lying outside the lower and upper 1% quantiles (i.e., BMI < 13.5 or BMI > 36.9) were considered misinput and replaced with missing values. The variable handedness was imputed using the last observation carried forward.

### Statistical analysis

#### Removal of site effects

For multi-site or multi-study neuroimaging studies, it is necessary to control for potential heterogeneity between sites to obtain unconfounded and generalizable results. Before estimating the main effects of age and sex on the global and regional brain measures (total GMV, total WMV, total sGMV, mean CT, regional GMV and regional CT), we applied ComBat^19^ and LongComBat^20^ in cross-sectional datasets and longitudinal datasets, respectively, to remove the potential site effects. The ComBat algorithm involves several steps including data standardization, site effect estimation, empirical Bayesian adjustment, removing estimated site effects, and data rescaling.

In the analysis of cross-sectional datasets, the models for ComBat were specified as a linear regression model illustrated below using total GMV:

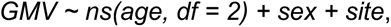

where *ns* denotes natural cubic splines on two degrees of freedom, which means that there were two boundary knots and one interval knot placed at the median of the covariate age. Splines were used to accommodate nonlinearity in the age effect. For the longitudinal datasets, the model for LongComBat used a linear mixed effects model with subject-specific random intercepts:

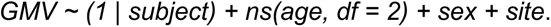

When estimating the effects of other non-brain covariates in the ABCD dataset, ComBat was used to control the site effects respectively for each of the cross-sectional covariates. The ComBat models were specified as illustrated below using GMV:

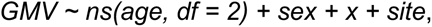

where *x* denotes the non-brain covariate. LongComBat was used for each of the longitudinal covariates with a linear mixed effects model with subject-specific random intercepts only:

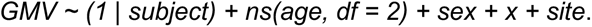

When estimating the effects of other covariates on the FC in the ABCD data, we additionally controlled for the mean frame displacement (FD) of the frames remaining after scrubbing. The longComBat models were specified as:

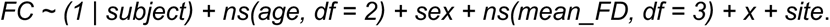

The Combat and LongComBat were implemented using the neuroCombat^58^ and longCombat^59^ R packages. Site effects were removed before all subsequent analyses including the bootstrap analyses described below.

#### Robust effect size index (RESI) for association strength

The RESI is a recently developed standardized effect size (ES) index that has consistent interpretation across many model types, encompassing all types of test statistics in most regression models^21,22^. Briefly, the RESI is a standardized ES parameter describing the deviation of the true parameter value(s) *β* from the reference value(s) *β*_0_ from the statistical null hypothesis *H*_0_: *β* = *β*_0_,

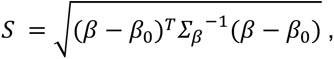

where *S* denotes the parameter RESI, *β* and *β*_0_ can be vectors, *Σ*_#_ is the covariance matrix for √*N β̂* (where *β̂* is the estimator for *β*) (Supplementary Information: Section 1).

In previous work, we defined a consistent estimator for RESI^21^,

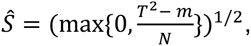

where *T*^2^ is the chi-square test statistics *T*\* = *N*(*β* – *β*_0_)*^T^Σ_β_*^-1^(*β* – *β*_0_) for testing the null hypothesis *H*_0_: *β* = *β*_0_, *m* is the number of parameters being tested (i.e., the length of *β*) and *N* is the number of subjects.

As RESI is generally applicable across different models and data types, it is also applicable to the situation where Cohen’s *d* was defined. In this scenario, the RESI is equal to ½ Cohen’s *d*^21^, so Cohen’s suggested thresholds for ES can be adopted for RESI: small (RESI=0.1); medium (RESI=0.25); and large (RESI=0.4). Because RESI is robust, when the assumptions of Cohen’s *d* are not satisfied, such as when the variances between the groups are not equal, RESI is still a consistent estimator, but Cohen’s *d* is not. The confidence intervals (CIs) for RESI in our analyses were constructed using 1,000 non-parametric bootstraps^22^.

The systematic difference in the standardized ESs between cross-sectional and longitudinal studies puts extra challenges on the comparison and aggregation of standardized ES estimates across studies with different designs. To improve the comparability of standardized ESs between cross-sectional and longitudinal studies, we proposed a new ES index: the cross-sectional RESI (i.e., CS-RESI) for longitudinal datasets. The CS-RESI for longitudinal datasets represents the RESI in the same study population if the longitudinal study had been conducted cross-sectionally. Detailed definition, point estimator, and CI construction procedure for CS-RESI can be found in Supplementary Information: Section 1. Comprehensive statistical simulation studies were also performed to demonstrate the valid performance of the proposed estimator and CI for CS-RESI (Supplementary Information: Section 1.2). With CS-RESI, we can quantify the benefit of using a longitudinal study design in a single dataset (Supplementary Information: Section 1.3).

#### Study-level models

After removing the site effects using ComBat or LongComBat in the multi-site data, we estimated the effects of age and sex on each of the global/regional brain measures using generalized estimating equations (GEEs) and linear regression models in the longitudinal datasets and cross-sectional datasets, respectively. The mean model was specified as below after ComBat/LongComBat:

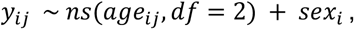

where *yij* was taken to be a global brain measure (i.e., total GMV, WMV, sGMV or mean CT) or regional brain measure (i.e., regional GMV or CT) at the *j*-th visit from the subject *i*, and *j* = 1 for cross-sectional datasets. The age effect was estimated with natural cubic splines with two degrees of freedom, which means that there were two boundary knots and one interval knot placed at the median of the covariate age. For the GEEs, we used an exchangeable correlation structure as the working structure and identity linkage function. The model assumes the mean was correctly specified, but made no assumption about the error distribution. The GEEs were fitted with the “geepack” package^60^ in R. We used the RESI as a standardized ES measure.

RESIs and confidence intervals (CIs) were computed using the “RESI” R package (version 1.2.0)^23^.

#### Meta-analysis of the age and sex effects

The weighted linear regression model for the meta-analysis of age effects across the studies was specified as:

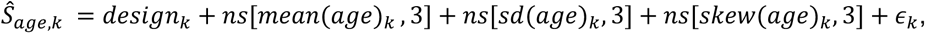

where *Ŝ*_age,k_ denotes the estimated RESI for study *k,* and the weights were the inverse of the standard error of each RESI estimate. The sample mean, standard deviation (SD), and skewness of the age were included as nonlinear terms estimated using natural splines with 3 degrees of freedom (i.e., two boundary knots plus two interval knots at the 33rd and 66^th^ percentiles of the covariates), and a binary variable indicating the design type (cross-sectional/longitudinal) was also included.

The weighted linear regression model for the meta-analysis of sex effects across the studies was specified as

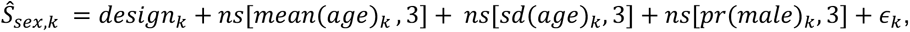

where *Ŝ_sex,k_* denotes the estimated RESI of sex for study *k,* and the weights were the inverse of the standard error of each RESI estimate. The sample mean, SD of the age covariate, and the proportion of males in each study were included as nonlinear terms estimated using natural splines with 3 degrees of freedom, and a binary variable indicating the design type (cross-sectional or longitudinal) was also included.

These meta-analyses were performed for each of the global and regional brain measures. Inferences were performed using robust standard errors^61^. In the partial regression plots, the expected standardized ESs for the age effects were estimated from the meta-analysis model after fixing mean age at 45, SD of age at 7 and/or skewness at 0; the expected standardized ESs for the sex effects were estimated from the meta-analysis model after fixing mean age at 45, SD of age at 7 and/or proportion of males at 0.5.

#### Sampling schemes for age in the UKB and ADNI

We used bootstrapping to evaluate the effect of different sampling schemes with different target sample covariate distributions on the standardized ESs and replicability in the cross-sectional UKB and longitudinal ADNI datasets. For a given sample size and sampling schemes, 1,000 bootstrap replicates were conducted. The standardized ES was estimated as the mean standardized ES (i.e., RESI) across the bootstrap replicates. The 95% confidence interval (CI) for the standardized ES was estimated using the lower and upper 2.5% quantiles across the 1,000 estimates of the standardized ES in the bootstrap replicates. Power was calculated as the proportion of bootstrap replicates producing *p*-values less than or equal to 5% for those associations that were significant at 0.05 in the full sample. In the UKB, only one region was not significant for age in each of GMV and CT, and in ADNI, only 1 and 4 regions were not significant for age in GMV and CT, respectively. Replicability in prior work has been defined as having a significant *p*-value and the same sign for the regression coefficient. Because we were fitting nonlinear effects, we defined replicability as the probability that two independent studies have significant *p*-values; this is equivalent to the definition of power squared. The 95% CIs for replicability were derived using Wilson’s method^62^.

In the UKB dataset, to modify the (between-subject) variability of the age variable we used the following three target sampling distributions (Fig. S3): bell-shaped, where the target distribution had most of the subjects distributed in the middle age range; uniform, where the target distribution had subjects equally distributed across the age range; and U-shaped, where the target distribution had most of the subjects distributed closer to the range limits of the age in the study. The samples with U-shaped age distribution had the largest sample variance of age, followed by the ones with uniform age distribution and the ones with bell-shaped age distribution. The bell-shaped and U-shaped functions were proportional to a quadratic function. To sample according to these distributions, each record was first inversely weighted by the frequency of the records with age falling in the range of +/- 0.5 years of the age for that record to achieve the uniform sampling distribution. Each record was then rescaled to derive the weights for bell- and U-shaped sampling distributions. The records with age <50 or >78 years were winsorized at 50 or 78 years when assigning weights, respectively, to limit the effects of outliers on the weight assignment, but the actual age values were used when analyzing each bootstrapped data.

In each bootstrap from the ADNI dataset, each subject was sampled to have two records. We modified the between- and within-subject variability of age, respectively, by making the “baseline age” follow one of the three target sampling distributions used for the UKB dataset and the “age change” independently follow one of three new distributions: decreasing; uniform; and increasing (Fig. S5). The increasing and decreasing functions were proportional to an exponential function. The samples with “increasing” distribution of “age change” had the largest within-subject variability of age, followed by the ones with the uniform distribution of “age change” and the ones with “decreasing” distribution of age change.

To modify the “baseline age” and the “age change” from baseline independently, we first created all combinations of the baseline record and one follow-up from each participant, and derived the “baseline age” and “age change” for each combination. The “bivariate” frequency of each combination was obtained as the number of combinations with values of “baseline age” and “age change” falling in the range of +/- 0.5 years of the values of “baseline age” and “age change” for this combination. Then each combination was inversely weighted by its “bivariate” frequency to target a uniform bivariate distribution of baseline age and age change. The weight for each combination was then rescaled to make the baseline age and age change follow different sampling distributions independently. The combinations with baseline age <65 or >85 years were winsorized at 65 or 85 years, and the combinations with age change greater than 5 years were winsorized at 5 years when assigning weights to limit the effects of outliers on the weight assignment, but the actual ages were used when analyzing each bootstrapped data.

The sampling methods could be easily extended to the scenario where each subject had three records (and more than three) in the bootstrap data by making combinations of the baseline and two follow-ups. Each combination was inversely weighted to achieve uniform baseline age and age change distributions, respectively, by the “trivariate” frequency of the combinations with baseline age and the two age changes from baseline for the two follow-ups falling into the range of +/- 0.5 years of the corresponding values for this combination. Since we only investigated the effect of modifying the number of measurements per subject under uniform between- and within-subject sampling schemes (Fig. 3c-d), we did not need to consider rescaling the weights here to achieve other sampling distributions but they could be done similarly. For the scenario where each subject only had one measurement (Fig. 3c-d), the standardized ESs and replicability were estimated only using the baseline measurements.

All site effects were removed using ComBat or LongComBat prior to performing the bootstrap analysis.

#### Sampling schemes for other non-brain covariates in ABCD

We used bootstrapping to study how different sampling strategies affect the RESI in the ABCD dataset. Each subject in the bootstrap data had two measurements. We applied the same weight assignment method described above for the ADNI dataset to modify the between- and within-subject variability of a covariate. We made the baseline covariate and the change in covariate follow bell-shaped and/or U-shaped distributions, to let the sample have larger or smaller between- and/or within-subject variability of the covariate, respectively. The baseline covariate and change of covariate were winsorized at the upper and lower 5% quantiles to limit the effect of outliers on sampling. For each cognitive variable, only the subjects with non-missing baseline measurements and at least one non-missing follow-up were included.

Generalized linear models (GLMs) and GEEs were fitted to estimate the effect of each non-brain covariate on the structural brain measures after controlling for age and sex,

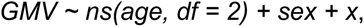

where *x* denotes one of the non-brain covariates. For the GEEs, we used an exchangeable correlation structure as the working structure and identity linkage function.

Only the between-subject sampling schemes were applied for the non-brain covariates that were stable over time (e.g., birth weight and handedness). In other words, the subjects were sampled based on their baseline covariate values, and then a follow-up was selected randomly for each subject. The sampling schemes to increase the between-subject variability in the covariate “handedness”, which was a binary variable (right-handed or not), was specified differently. The expected proportion of right-handed subjects in the bootstrap samples was 50% under the sampling scheme with “larger” between-subject variability and 10% under the sampling scheme with “smaller” between-subject variability.

For given between- and/or within-subject sampling schemes, We obtained 1,000 bootstrap replicates. The standardized ES was estimated as the mean standardized ES across the bootstrap replicates. The 95% CIs for standardized ES were estimated using the lower and upper 2.5% quantiles across the 1,000 estimates of the standardized ES in the bootstrap replicates. The sample sizes needed for 80% replicability were estimated based on the (mean) estimated standardized ES and *F*-distribution (see below).

#### Analysis of FC in ABCD

In a subset of the ABCD where we have preprocessed longitudinal FC data at two timepoints (baseline and 2-year follow-up), we only restricted to the subjects with non-missing measurements at both of the two times. In the GEEs used to estimate the effects of non-brain covariates on FC, the mean model was specified as below after LongComBat:

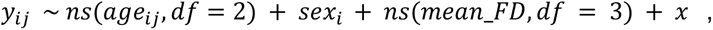

where *yij* was taken to be an FC outcome and *x* denotes a non-brain covariate. Mean frame-wise displacement (FD) (*mean_FD*) was also included as a covariate with natural cubic splines with 3 degrees of freedom. We used an exchangeable correlation structure as the working structure and identity linkage function in the GEEs. The frame count of each scan was used as the weights.

When evaluating the effect of different sampling schemes on the standardized ESs, we obtained 1,000 bootstrap replicates for given between- and/or within-subject sampling schemes. The standardized ES was estimated as the mean standardized ES across the bootstrap replicates. CIs were computed as described above. The sample sizes needed for 80% replicability were estimated based on the (mean) estimated standardized ESs and *F*-distribution (see below).

#### Sample size calculation for a target power or replicability with a given standardized effect size

After estimating the standardized ES for an association, the calculation of the corresponding sample size *N* needed for detecting this association with *γ* × 100% power at significance level of *α* was based on an *F*-distribution. Let *df* denote the total degree of freedom of the analysis model, *F*(*x*; *λ*) denotes the cumulative density function for the random variable *X*, which follows the (non-central) *F*-distribution with degrees of freedom being 1 and *N - df* and non-centrality parameter *λ*. The corresponding sample size *N* is:

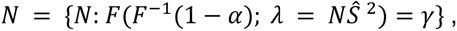

where *Ŝ* is the estimated RESI for the standardized ES. Power curves for the RESI are given in Figure 3 of Vandekar et al. (2002)^21^. Replicability was defined as the probability that two independent studies have significant *p*-values, which is equivalent to power squared.

#### Estimation of the between- and within-subject effects

For the non-brain covariates that were analyzed longitudinally in ABCD dataset, GEEs with exchangeable correlation structures were fitted to estimate their cross-sectional and longitudinal effects on structural and functional brain measures after controlling for age and sex, respectively. The mean model was specified as illustrated with GMV:

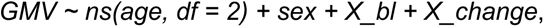

where *X_bl* denotes the subject-specific baseline covariate values, and the *X_change* denotes the difference of the covariate value at each visit to the subject-specific baseline covariate value (see Supplementary Information: Section 3.2). The subjects without baseline measures were not included in the modeling. The model coefficients for the terms “*X_bl*” and “*X_change*” represent the between- and within-subject effects of this non-brain covariate on total GMV, respectively. For the FC data, the same covariates and weighting were used as described above. Using the first time point as the between-subject term was a special case that ensured comparing the parameter using the baseline cross-sectional model was equal to the parameter for the between-subject effect in the longitudinal model. In this model, the between-subject variance was defined as the variance of the baseline measurement and the within-subject variance was the mean square of *X_change*. This model specification ensured that the sampling schemes independently affected the between- and within-subject variances separately (Supplementary Information: Section 3.2 Eqn (17)).

## Data availability

Participant-level data from many datasets are available according to study-level data access rules. Study-level model parameters are available at https://github.com/KaidiK/RESI_BWAS. We acknowledge the usage of several openly shared MRI datasets, which are available at the respective consortia websites and are subject to the sharing policies of each consortium: OpenNeuro (https://openneuro.org/), UK BioBank (https://www.ukbiobank.ac.uk/), ABCD (https://abcdstudy.org/), the Laboratory of NeuroImaging (https://loni.usc.edu/), data made available through the Open Science Framework (https://osf.io/), the Human Connectome Project (http://www.humanconnectomeproject.org/), and the OpenPain project (https://www.openpain.org). The ABCD data repository grows and changes over time. The ABCD data used in this report came from DOI 10.15154/1503209. Data used in this article were provided by the brain consortium for reliability, reproducibility and replicability (3R-BRAIN) (https://github.com/zuoxinian/3R-BRAIN). Data used in the preparation of this article was obtained from the Australian Imaging Biomarkers and Lifestyle flagship study of aging (AIBL) funded by the Commonwealth Scientific and Industrial Research Organisation (CSIRO) which was made available at the ADNI database (https://adni.loni.usc.edu/aibl-australian-imaging-biomarkers-and-lifestyle-study-of-ageing-18-month-data-now-released/). The AIBL researchers contributed data but did not participate in analysis or writing of this report. AIBL researchers are listed at https://www.aibl.csiro.au. Data used in preparation of this article were obtained from the Alzheimer’s Disease Neuroimaging Initiative (ADNI) database (https://adni.loni.usc.edu/). The investigators within the ADNI contributed to the design and implementation of ADNI and/or provided data but did not participate in analysis or writing of this report. A complete listing of ADNI investigators can be found at https://adni.loni.usc.edu/wp-content/uploads/how_to_apply/ADNI_Acknowledgement_List.pdf. More information on the ARWIBO consortium can be found at https://www.arwibo.it/. More information on CALM team members can be found at https://calm.mrc-cbu.cam.ac.uk/team/ and in the Supplementary Information. Data used in this article were obtained from the developmental component ‘Growing Up in China’ of the Chinese Color Nest Project (http://deepneuro.bnu.edu.cn/?p=163). Data used in the preparation of this article were obtained from the IConsortium on Vulnerability to Externalizing Disorders and Addictions (c-VEDA), India (https://cveda-project.org/). Data used in the preparation of this article were obtained from the Harvard Aging Brain Study (HABS P01AG036694) (https://habs.mgh.harvard.edu). Data used in the preparation of this article were obtained from the IMAGEN consortium (https://imagen-europe.com/). The POND network (https://pond-network.ca/) is a Canadian translational network in neurodevelopmental disorders, primarily funded by the Ontario Brain Institute.

The LBCC dataset used in the preparation of this article includes data obtained from the Alzheimer’s Disease Neuroimaging Initiative (ADNI) database (adni.loni.usc.edu). The ADNI was launched in 2003 as a public-private partnership, led by Principal Investigator Michael W. Weiner, MD. The primary goal of ADNI has been to test whether serial magnetic resonance imaging (MRI), positron emission tomography (PET), other biological markers, and clinical and neuropsychological assessment can be combined to measure the progression of mild cognitive impairment (MCI) and early Alzheimer’s disease (AD). Its data collection and sharing for this project was funded by the Alzheimer’s Disease Neuroimaging Initiative (ADNI) (National Institutes of Health Grant U01 AG024904) and DOD ADNI (Department of Defense award number W81XWH-12-2-0012). ADNI is funded by the National Institute on Aging, the National Institute of Biomedical Imaging and Bioengineering, and through generous contributions from the following: AbbVie, Alzheimer’s Association; Alzheimer’s Drug Discovery Foundation; Araclon Biotech; BioClinica, Inc.; Biogen; Bristol-Myers Squibb Company; CereSpir, Inc.; Cogstate; Eisai Inc.; Elan Pharmaceuticals, Inc.; Eli Lilly and Company; EuroImmun; F. Hoffmann-La Roche Ltd and its affiliated company Genentech, Inc.; Fujirebio; GE Healthcare; IXICO Ltd.; Janssen Alzheimer Immunotherapy Research & Development, LLC.; Johnson & Johnson Pharmaceutical Research & Development LLC.; Lumosity; Lundbeck; Merck & Co., Inc.; Meso Scale Diagnostics, LLC.; NeuroRx Research; Neurotrack Technologies; Novartis Pharmaceuticals Corporation; Pfizer Inc.; Piramal Imaging; Servier; Takeda Pharmaceutical Company; and Transition Therapeutics. The Canadian Institutes of Health Research is providing funds to support ADNI clinical sites in Canada. Private sector contributions are facilitated by the Foundation for the National Institutes of Health (www.fnih.org). The grantee organization is the Northern California Institute for Research and Education, and the study is coordinated by the Alzheimer’s Therapeutic Research Institute at the University of Southern California. ADNI data are disseminated by the Laboratory for Neuro Imaging at the University of Southern California.

## Code availability

All code used to produce the analyses presented in this paper is available at https://github.com/KaidiK/RESI_BWAS.

## Acknowledgments

S.N.V was supported by R01MH123563. A.A.B. and J.S. were partially supported by R01MH132934 and R01MH133843 . Data processing done at the University of Cambridge is supported by the NIHR Cambridge Biomedical Research Centre (BRC-1215-20014) and NIHR Applied Research Collaboration East of England. Any views expressed are those of the author(s) and not necessarily those of the funders, IHU-JU2, the NIHR or the Department of Health and Social Care.

## Extended Data Figures/Tables

### Extended Data Figures

**Fig. S1.**
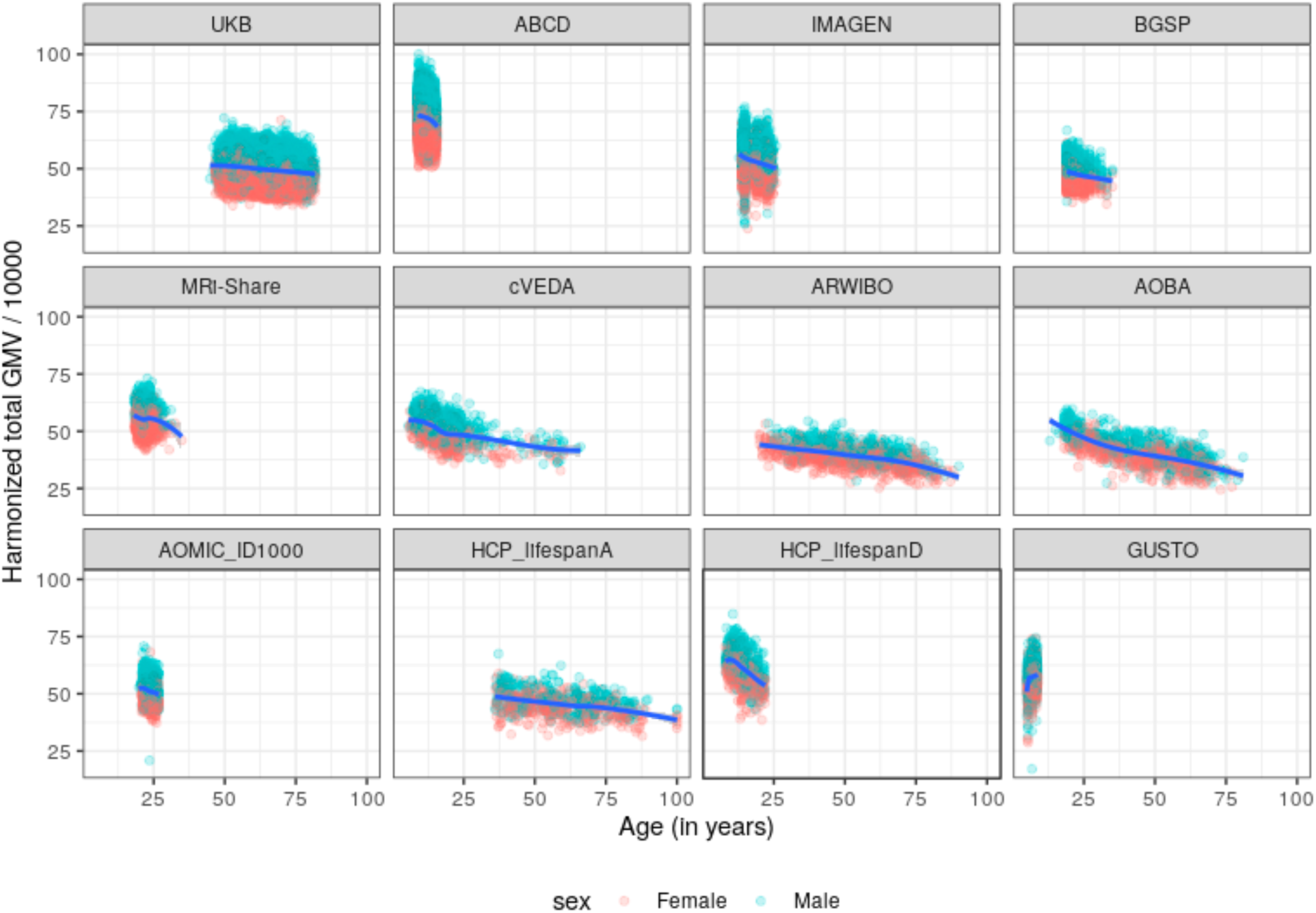
Scatterplots of the harmonized total gray matter volume (GMV) measures (i.e., total GMV after removing site effects using ComBat or longCombat) and age (in years) in the 12 largest studies among the 63 studies. The blue curves are the locally estimated scatterplot smoothing (LOESS) curves. The shaded areas are the 95% confidence bands. MRI datasets containing 77,695 scans from 63 studies with different age sample distributions were used. Nonlinear models are used to fit total GMV, white matter volume (WMV), subcortical gray matter volume (sGMV), and mean cortical thickness (CT) as a function of age (in years) and sex in each study.

**Fig. S2.**
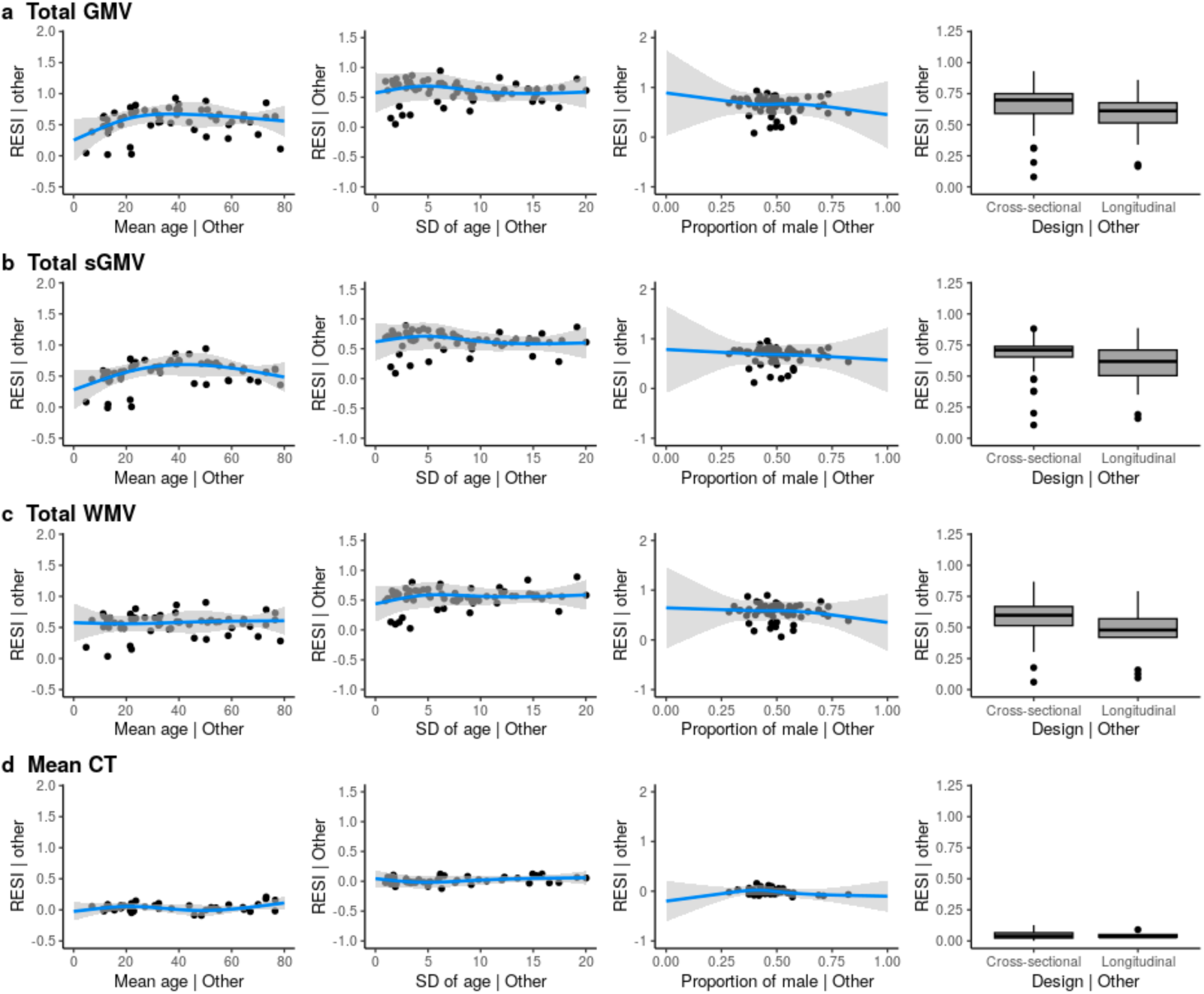
Meta-analysis results regarding the effects of study features on the associations between sex and different brain measures. (**a-d**) Partial regression plots of the meta-analyses of standardized effect sizes (i.e., RESI) for the associations between sex and global brain measures (**a**) total gray matter volume (GMV), (**b**) total subcortical gray matter volume (sGMV), (**c**) total white matter volume (WMV), and (**d**) mean cortical thickness (CT) (“| Other” means after fixing the other features at constant levels: design = cross-sectional, mean age = 45 years, sample age standard deviation (SD) = 7 and/or proportion of males = 0.5). The blue curves are the expected standardized effect size for sex. The gray areas are the 95% confidence bands.

**Fig. S3.**
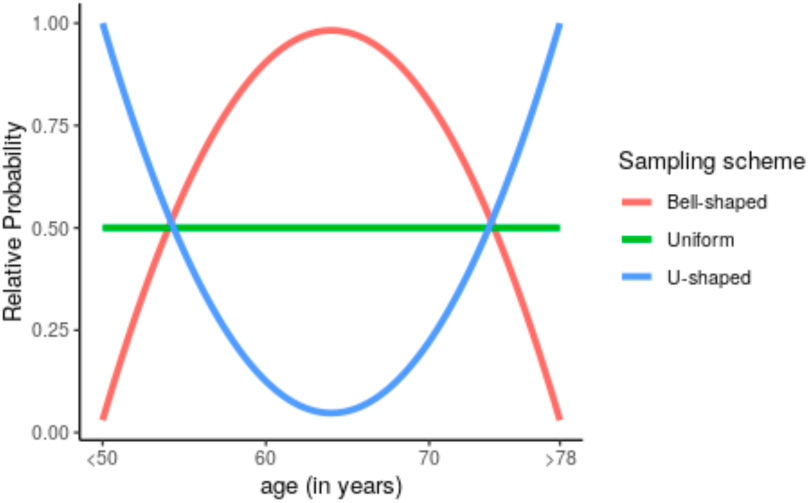
The sampling scheme implemented in UKB. The sampling schemes adjust the variability of age in the samples by assigning heavier or lighter weights to the subjects with age at the two tails of the population. The U-shaped scheme produces the largest variability of age in the samples, followed by uniform and bell-shaped sampling schemes.

**Fig. S4.**
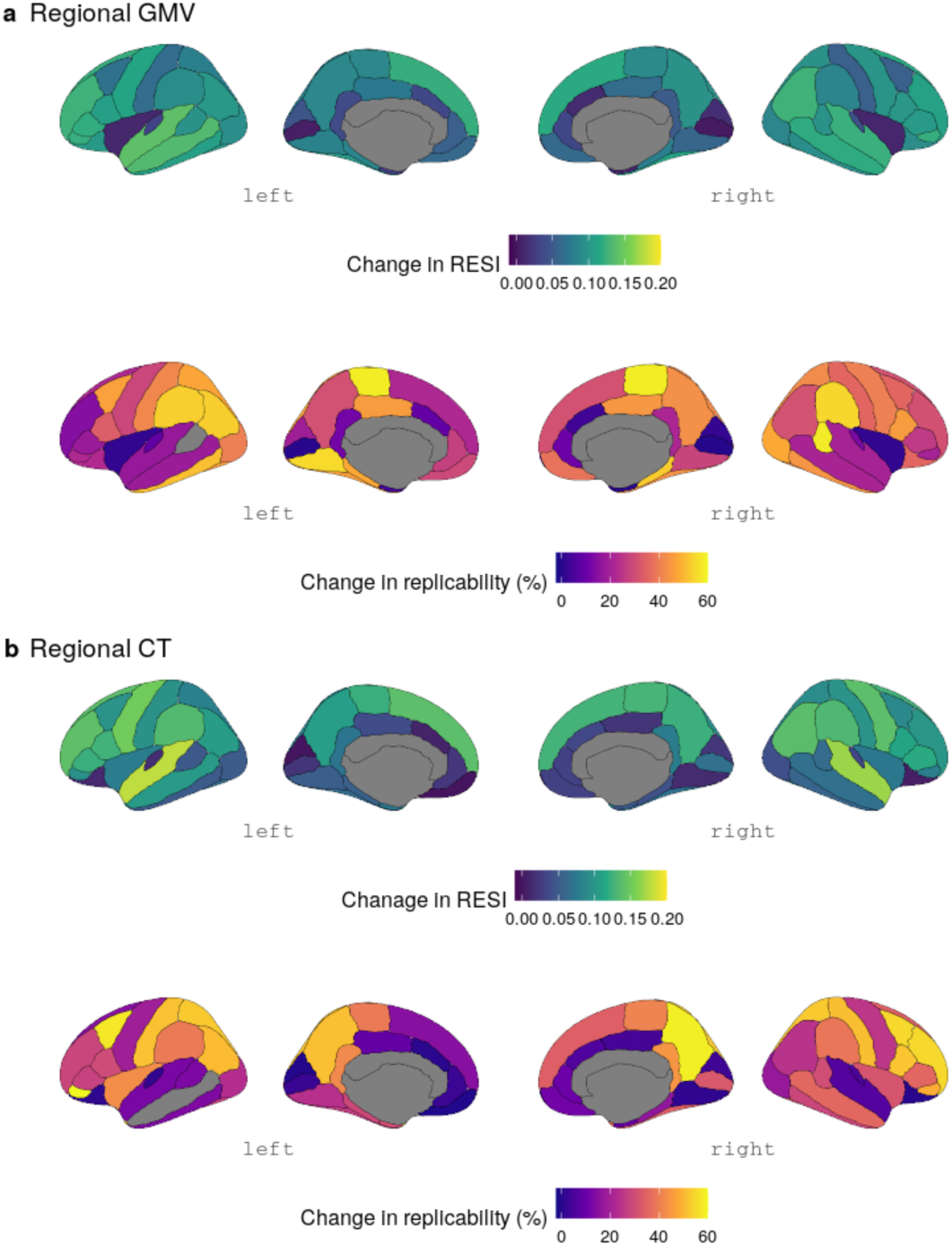
Region-specific improvement in the RESI and replicability in UKB for the association between age and (**a**) regional gray matter volume (GMV) and (**b**) regional cortical thickness (CT), respectively, by using U-shaped sampling scheme compared with bell-shaped sampling scheme, when *N* = 300.

**Fig. S5.**
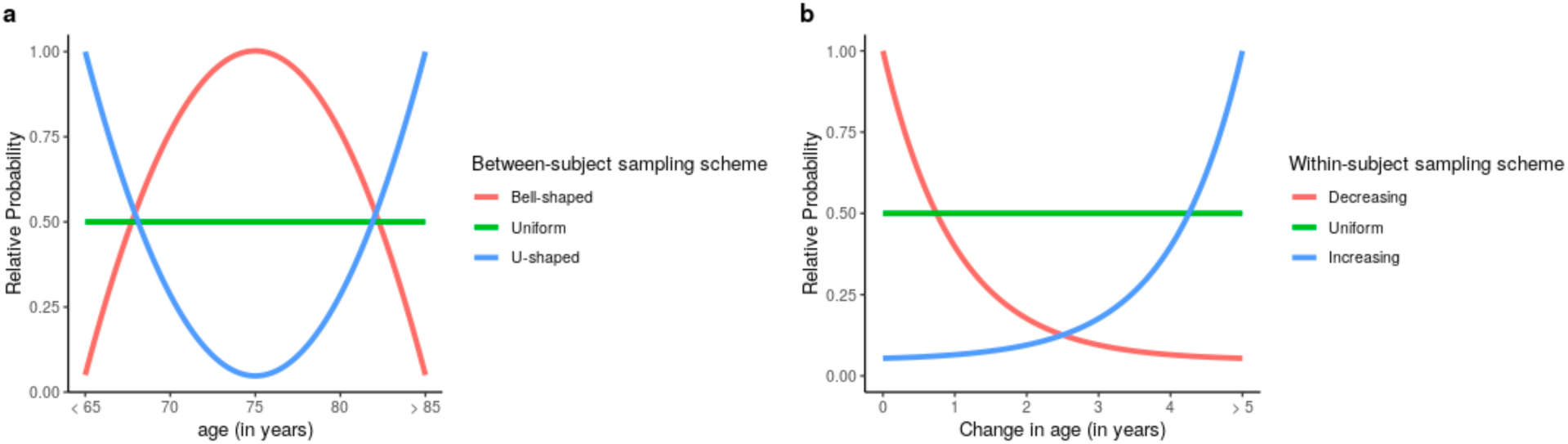
Between- and within-subject sampling schemes implemented in ADNI. (**a**) The between-subject variability of age is adjusted by assigning heavier or lighter weights to the subjects with baseline age closer to the two tails of the population baseline age distribution. (**b**) The within-subject variability in age is adjusted by increasing or decreasing the probability of selecting the follow-up observation(s) with a larger change in age since baseline.

**Fig. S6.**
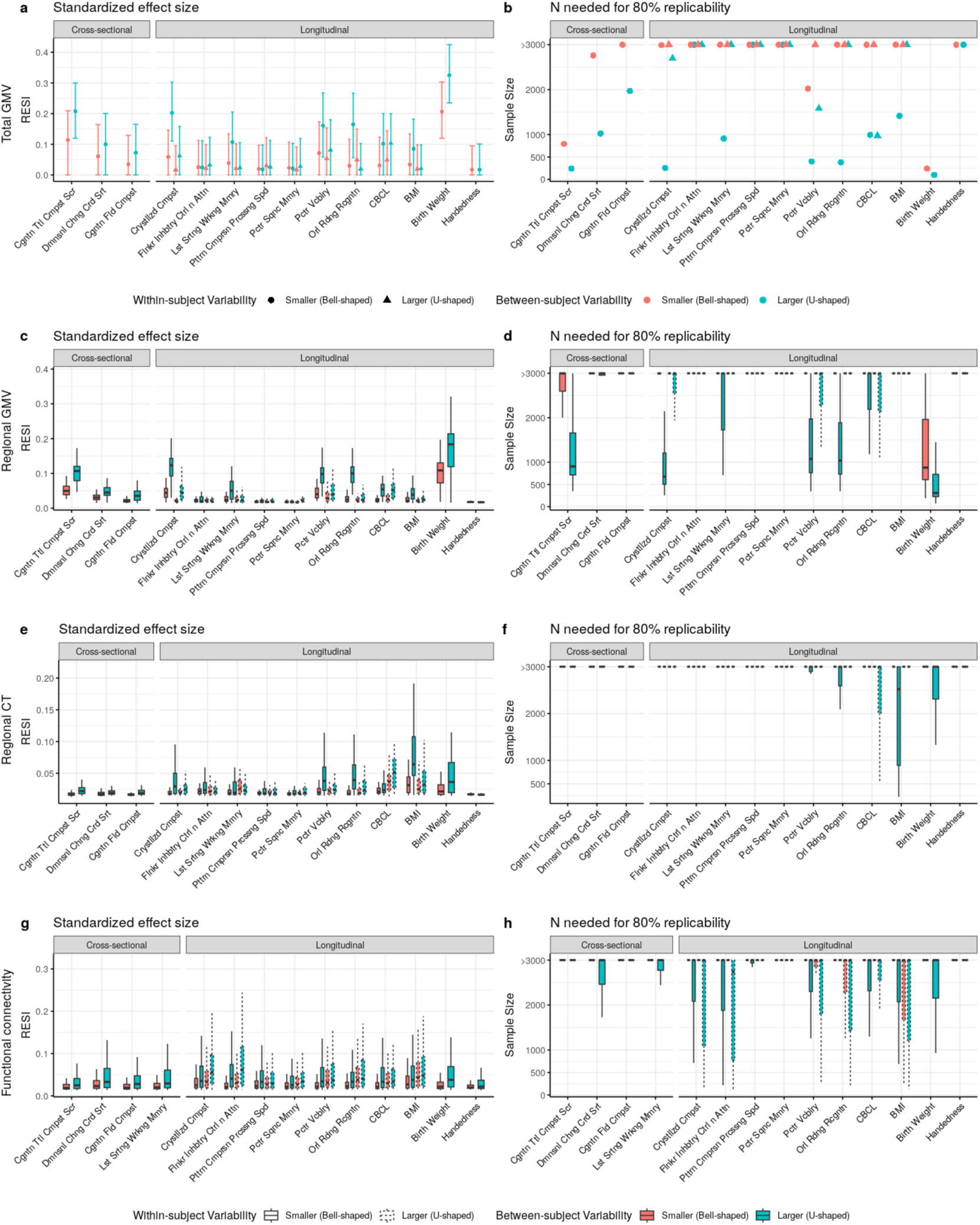
Heterogeneous improvement of standardized effect sizes (ESs) for cognitive, mental health, and demographic associations with structural and functional brain measures in the ABCD study with bootstrapped samples of *N* = 500. (**a**) U-shaped between-subject sampling scheme (blue) that increases between-subject variability of the non-brain covariate produces larger standardized ESs and (**b**) reduces the number of participants scanned to obtain 80% replicability in total gray matter volume (GMV). The points and triangles are the average standardized ESs across bootstraps and the whiskers are the 95% confidence intervals. Increasing within-subject sampling (triangles) can reduce standardized ESs. A similar pattern holds in (**c-d**) regional GMV and (**e-f**) regional cortical thickness (CT); boxplots show the distributions of the standardized ESs across regions. In contrast, (**g**) regional pairwise functional connectivity (FC) standardized ESs are improved by increasing between-(blue) and within-subject variability (dashed borders) with a corresponding reduction in the (**h**) number of participants scanned for 80% replicability.

**Fig. S7.**
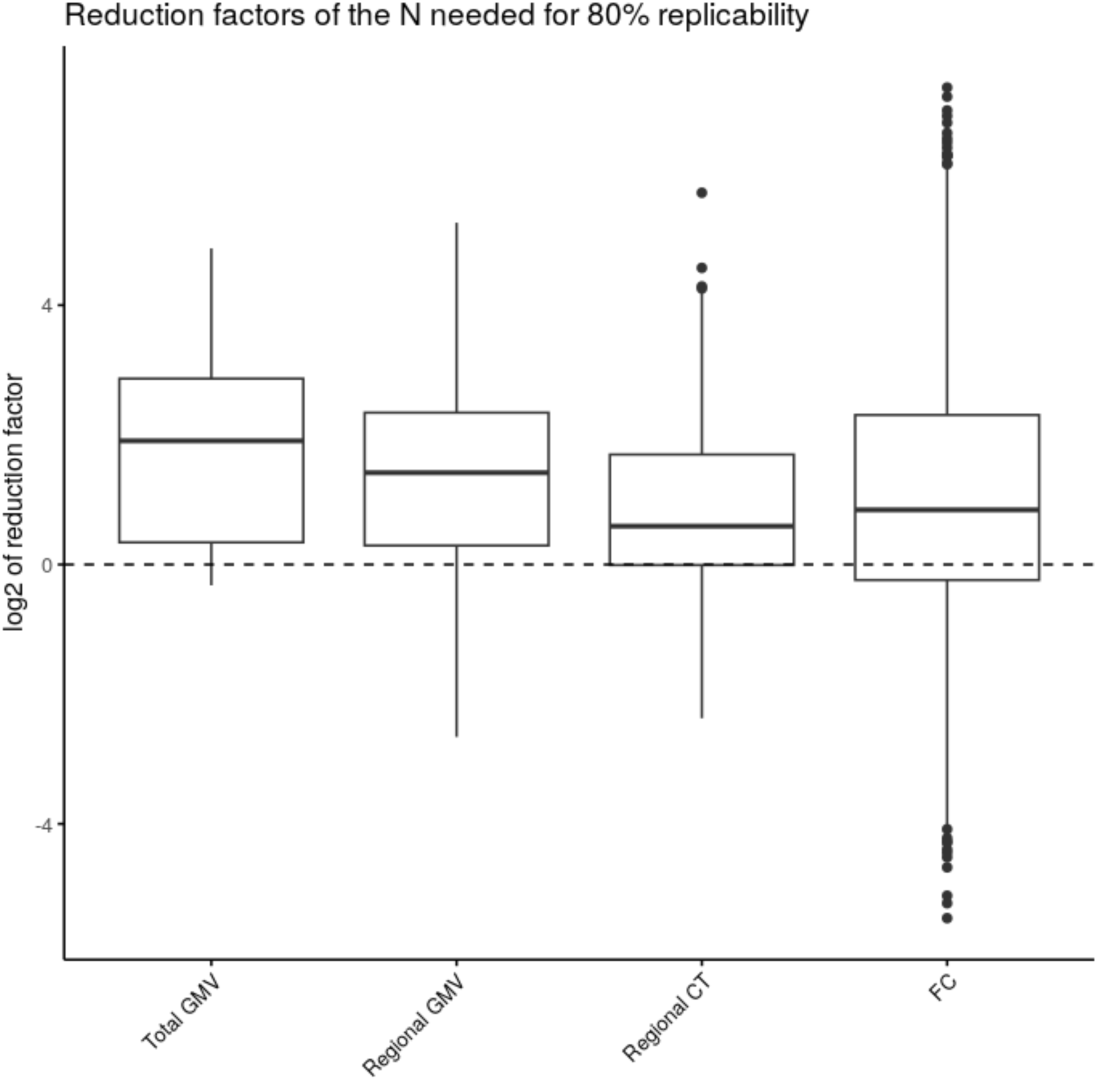
The boxplots showing the distributions of (log2 of) reduction factors of the sample size *N* needed for 80% replicability by increasing between-subject variability of the covariates across all the associations with each of the outcomes in Fig. 4. The reduction factors are derived by comparing the sample sizes needed for 80% replicability with U-shaped to the one with bell-shaped between-subject sampling scheme when the within-subject sampling scheme is bell-shaped (Fig. S6).

**Fig. S8.**
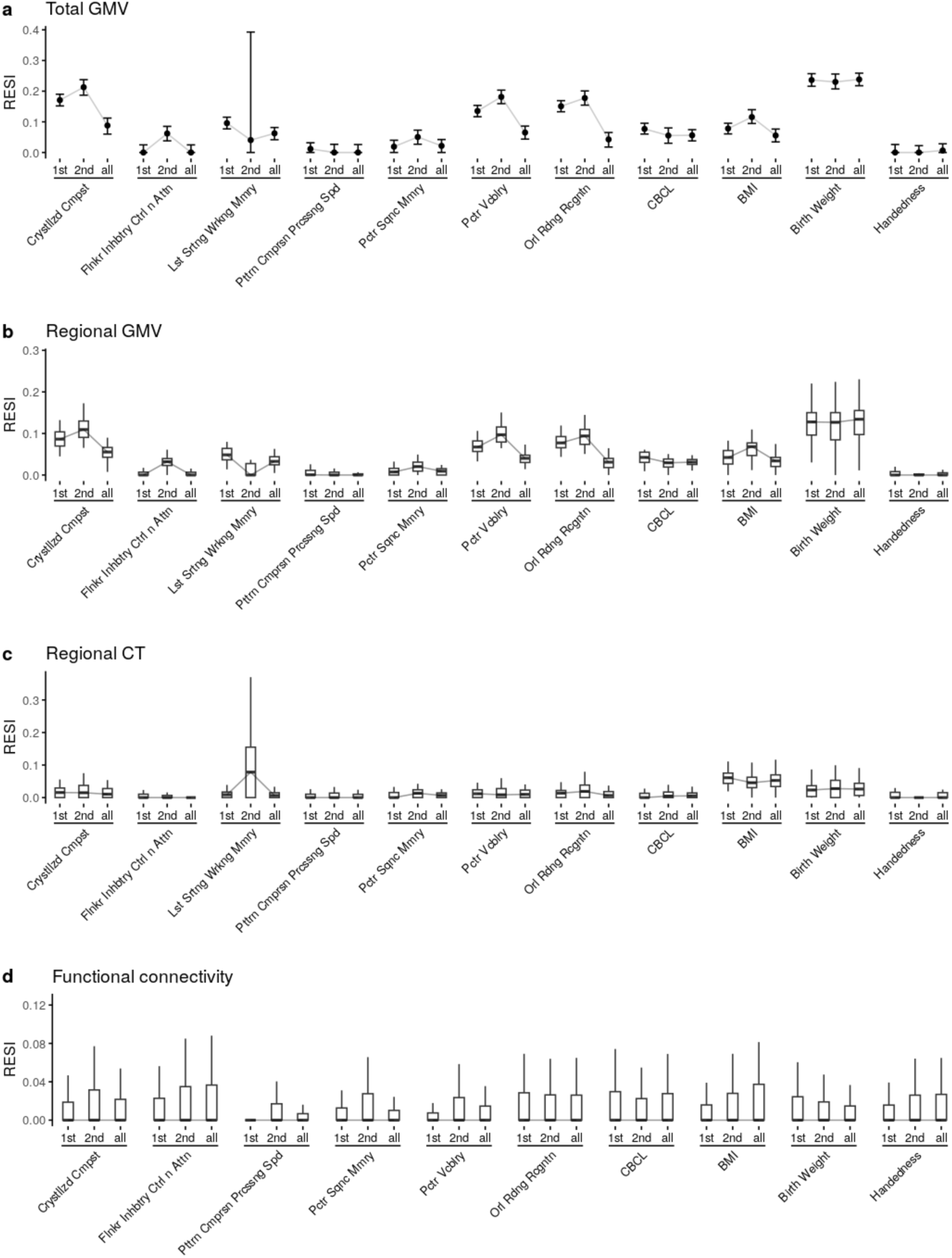
Longitudinal study designs can reduce standardized effect sizes (ESs) and replicability. Boxplots show the distributions of the standardized ESs across regions. The cross-sectional analyses use only the baseline or the 2nd measures (indicated by “1st”s or “2nd”s on the x-axes, respectively). The longitudinal analyses use the full longitudinal data (indicated by “all”s on the x-axes). (**a-c**) Cross-sectional analyses can have larger standardized ESs than the same longitudinal analyses for structural brain measures in ABCD. (**d**) The functional connectivity (FC) measures have a slight benefit of longitudinal modeling.

**Fig. S9.**
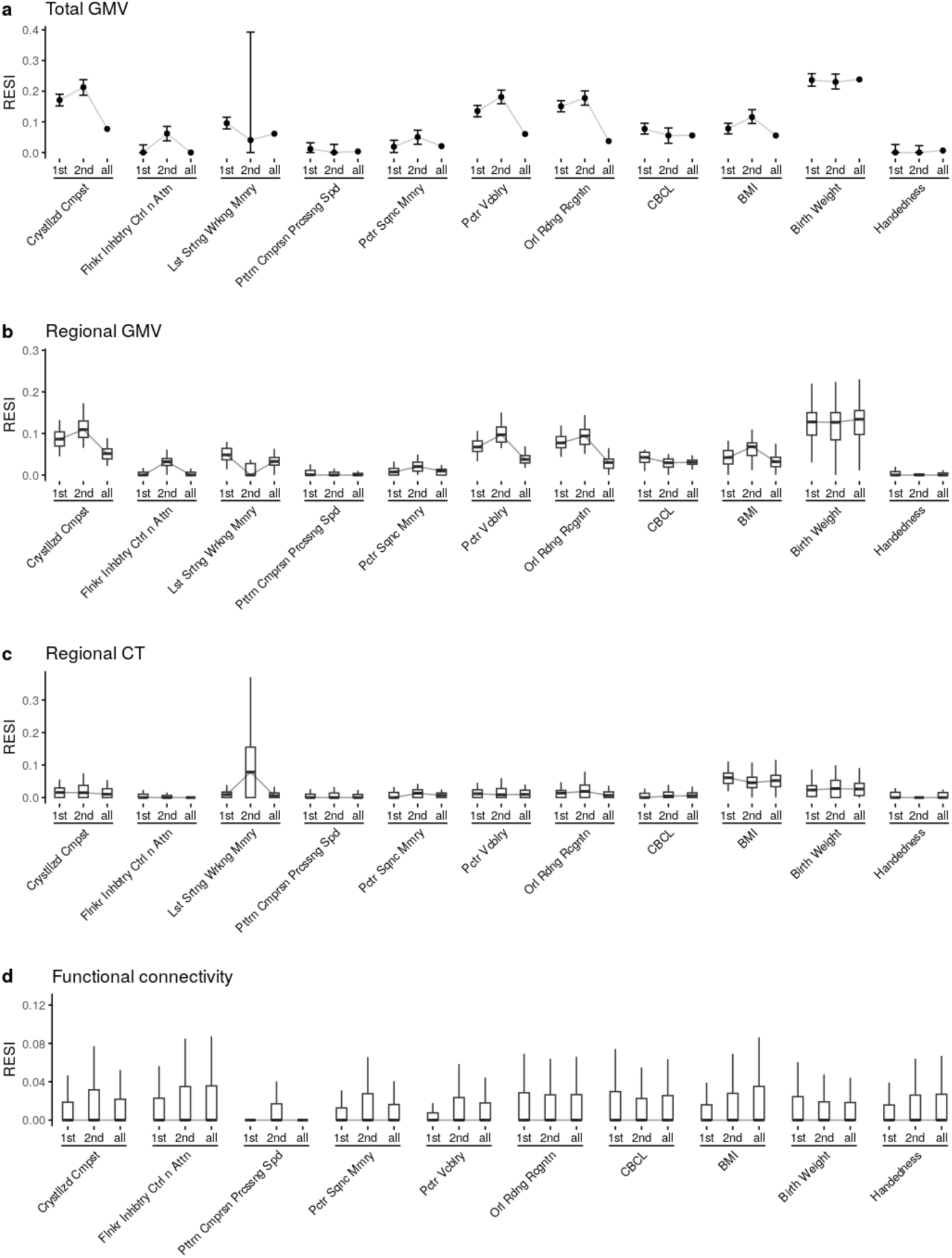
Replication of Fig. S8 using linear mixed models (LMMs) with individual-specific random intercepts. Boxplots show the distributions of the standardized effect sizes (ESs) across regions. The estimates for the RESI from cross-sectional analyses that only use the baseline measures and the first follow-up measures after baseline are indicated by “1st”s and “2nd”s on the x-axes, respectively; the estimates for RESI from longitudinal analyses that use the full longitudinal data are indicated by “all”s on the x-axes. The results are similar using generalized estimating equations (GEEs) and LMMs and show that longitudinal study designs can reduce standardized ESs for most structural brain associations. Only the point estimates of RESI are obtained as the bootstrap confidence intervals (CIs) in LMMs have not been evaluated.

**Fig. S10.**
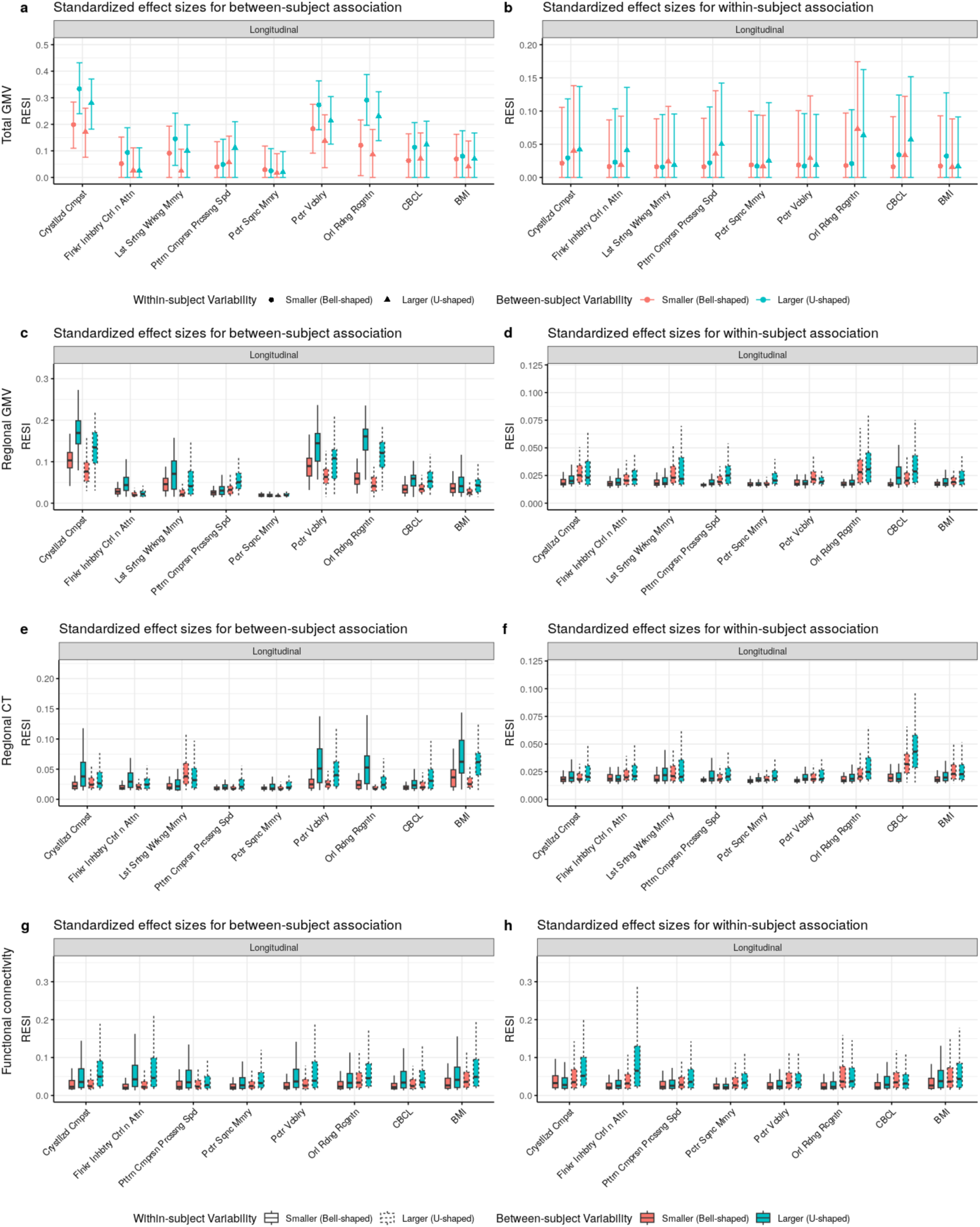
The influence of sampling schemes on the standardized effect sizes (ESs) for between- and within-subject associations, respectively, of cognition, mental health, and demographic covariates with different brain measures in the ABCD study at *N* = 500. Boxplots show the distribution of the standardized ESs across regions. Between-subject standardized ESs are predominantly affected by the between-subject variance, whereas within-subject standardized ESs are predominantly affected by the within-subject variance. The results for covariates birthweight and handedness, which do not vary within subjects, are not included as the within-subject sampling schemes do not apply to them.

**Fig. S11.**
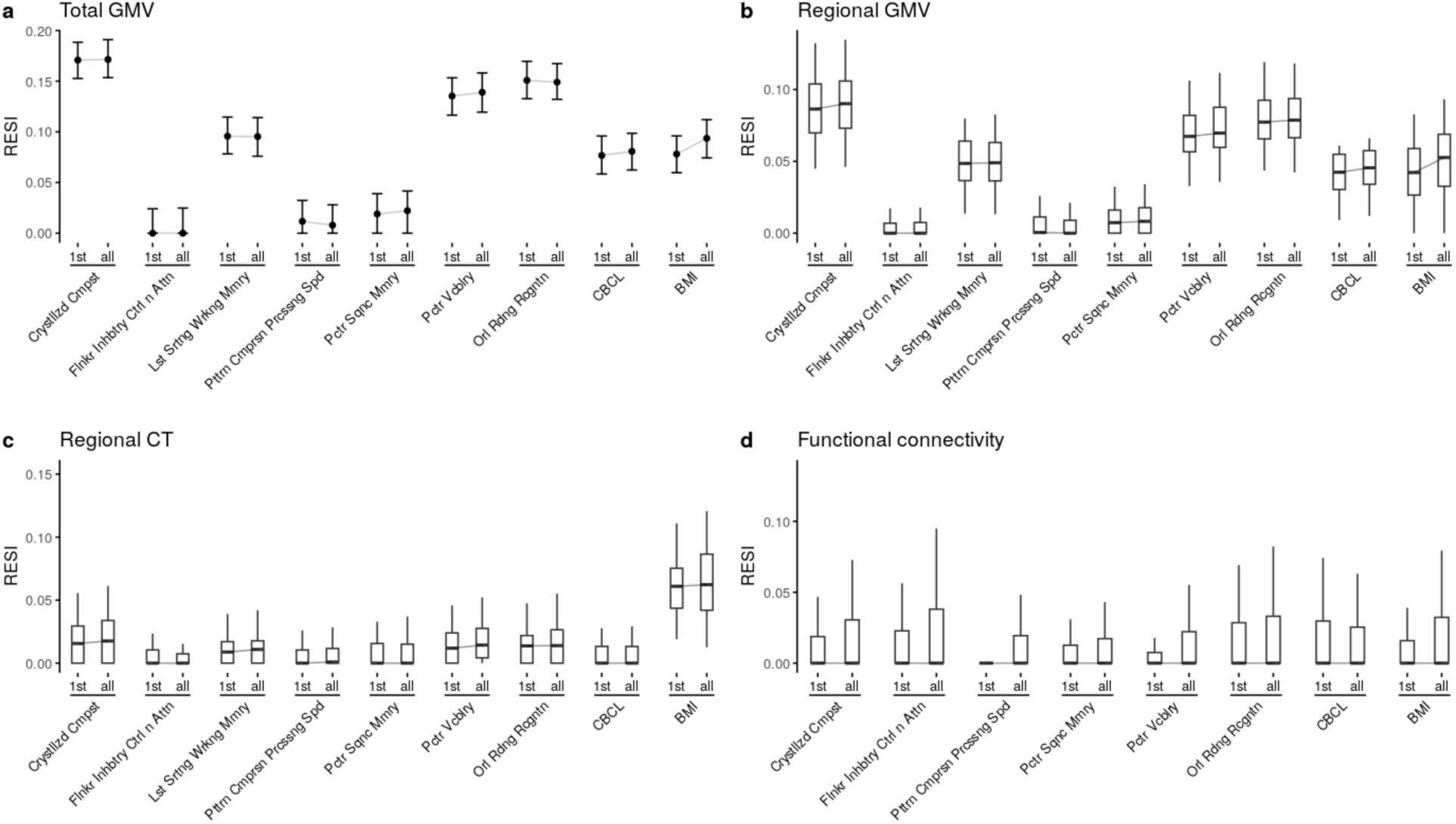
The estimated standardized effect sizes (ESs) from cross-sectional and longitudinal analyses, respectively, for the between-subject associations for cognition, mental health, and demographic covariates with different brain measures in the ABCD study. The estimated RESIs for cross-sectional analyses (that only use the baseline measures) are indicated by “1st”s on the x-axes; the estimated RESIs for the between-subject effects from longitudinal analyses (that use the full longitudinal data and a specification of separate between- and within-subject effects (see Methods: *Estimation of the between- and within-subject effects*) are indicated by “all”s on the x-axes. By separating the between- and within-subject effects in the longitudinal model, we can avoid averaging the different between- and within-subject effects and maintain the benefit of longitudinal designs on the estimated RESIs for the between-subject effects. The results for covariates birthweight and handedness are not included, as they do not vary within-subjects so only their between-subject effects can be estimated (which have already been shown in Fig. S8).

**Fig. S12.**
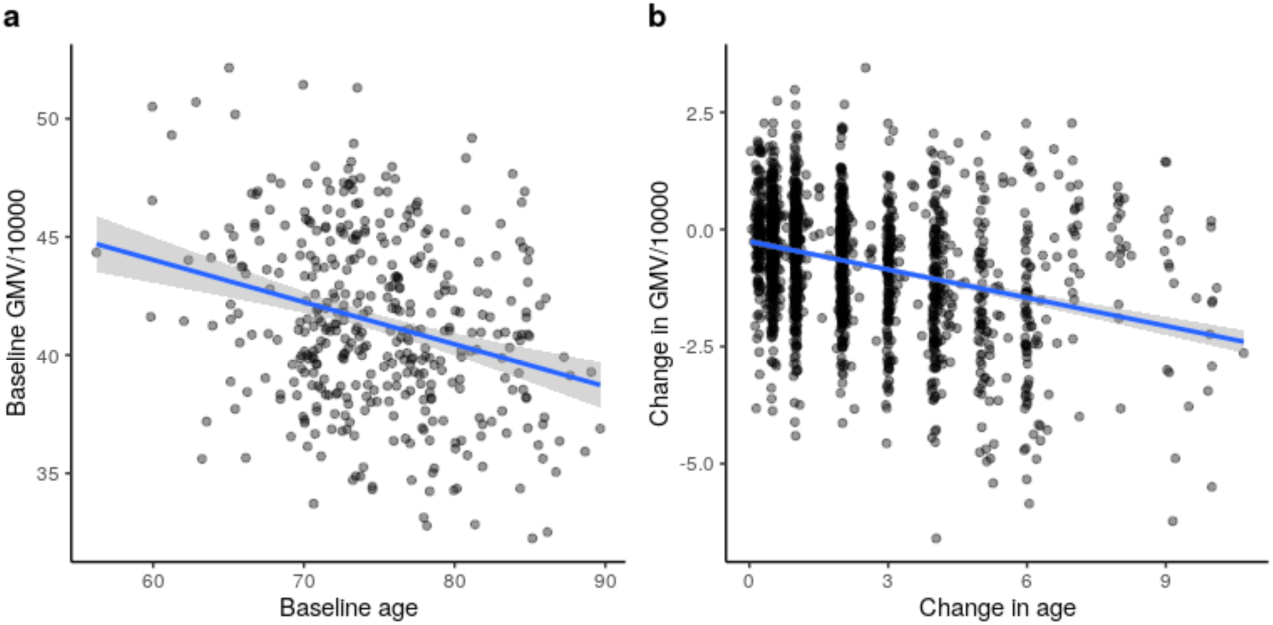
Visualization of the strengths of crude between- and within-subject associations between age and total gray matter volume (GMV) in ADNI. There are strong associations (**a**) between baseline total GMV and baseline age and (**b**) between change in total GMV and change in age since baseline, indicating there exist strong between- and within-subject effects of age on total GMV in ADNI study. The blue lines are the simple linear regression lines and the gray areas are the 95% confidence bands.

**Fig. S13:**
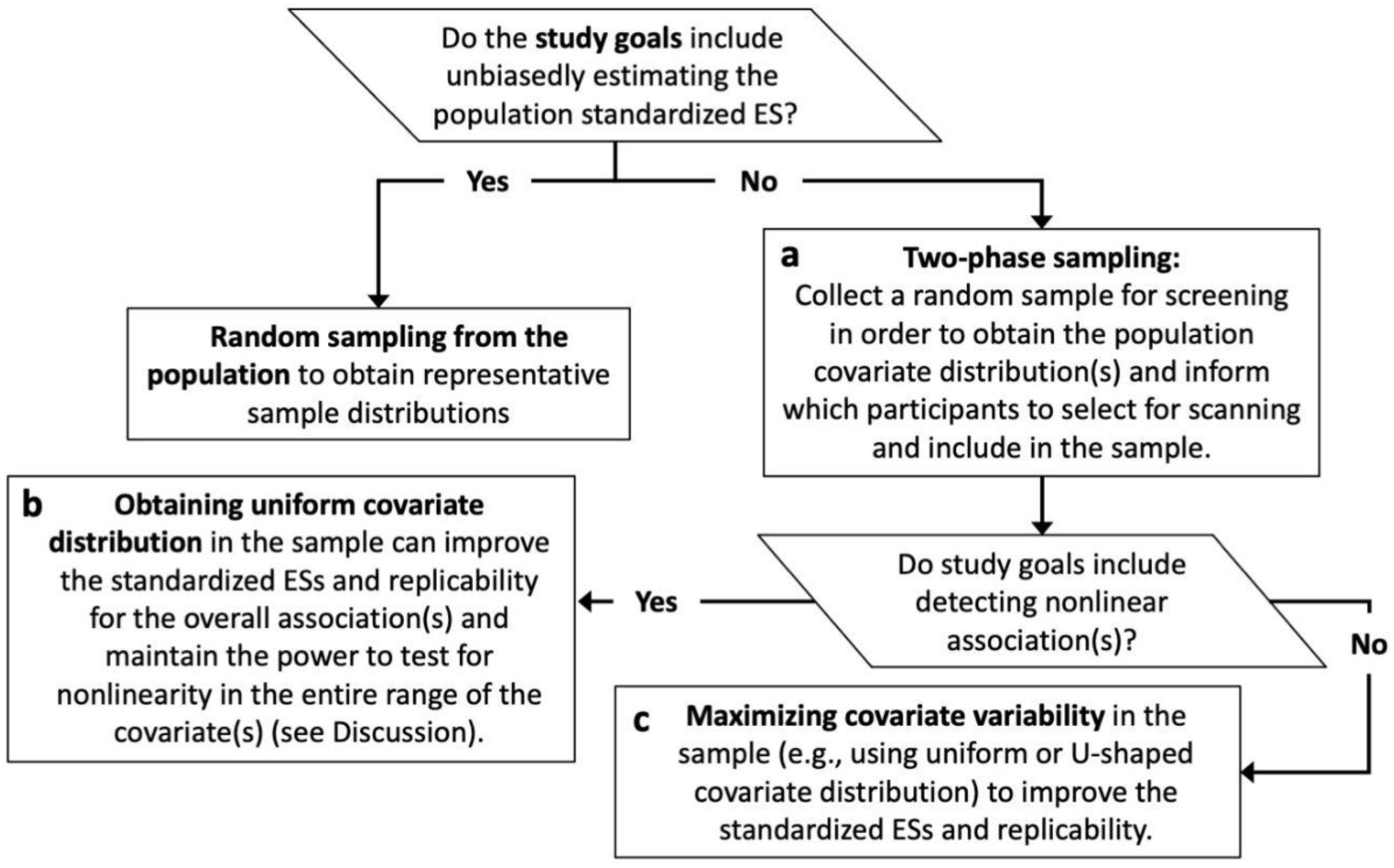
Decision tree for modified sampling strategy for a single primary covariate. Random/representative sampling is needed to unbiasedly estimate the variance of the covariate distribution in the population in order to obtain unbiased standardized effect size (ES) estimates in the population. (**a**) A two-phase design is needed to modify the covariate distribution(s) in the sample to improve the standardized ES and replicability, where random sampling is performed first in a larger dataset to collect covariate values and biased sampling a subset based on collected covariates values to optimize the standardized ESs and replicability; unbiased population standardized ES estimates still can be obtained using weighted estimation (see Discussion: *Sampling strategies can increase replicability*). (**b**) If the distribution(s) of the covariate(s) in the population is bell-shaped, a uniform covariate distribution in the sample can still increase the standardized ES and replicability in detecting the overall association. (**c**) The particular target distribution will depend on the difficulty of collecting participants in the tail of the distributions (see Supplementary Information: Section 2.1).

**Fig. S14:**
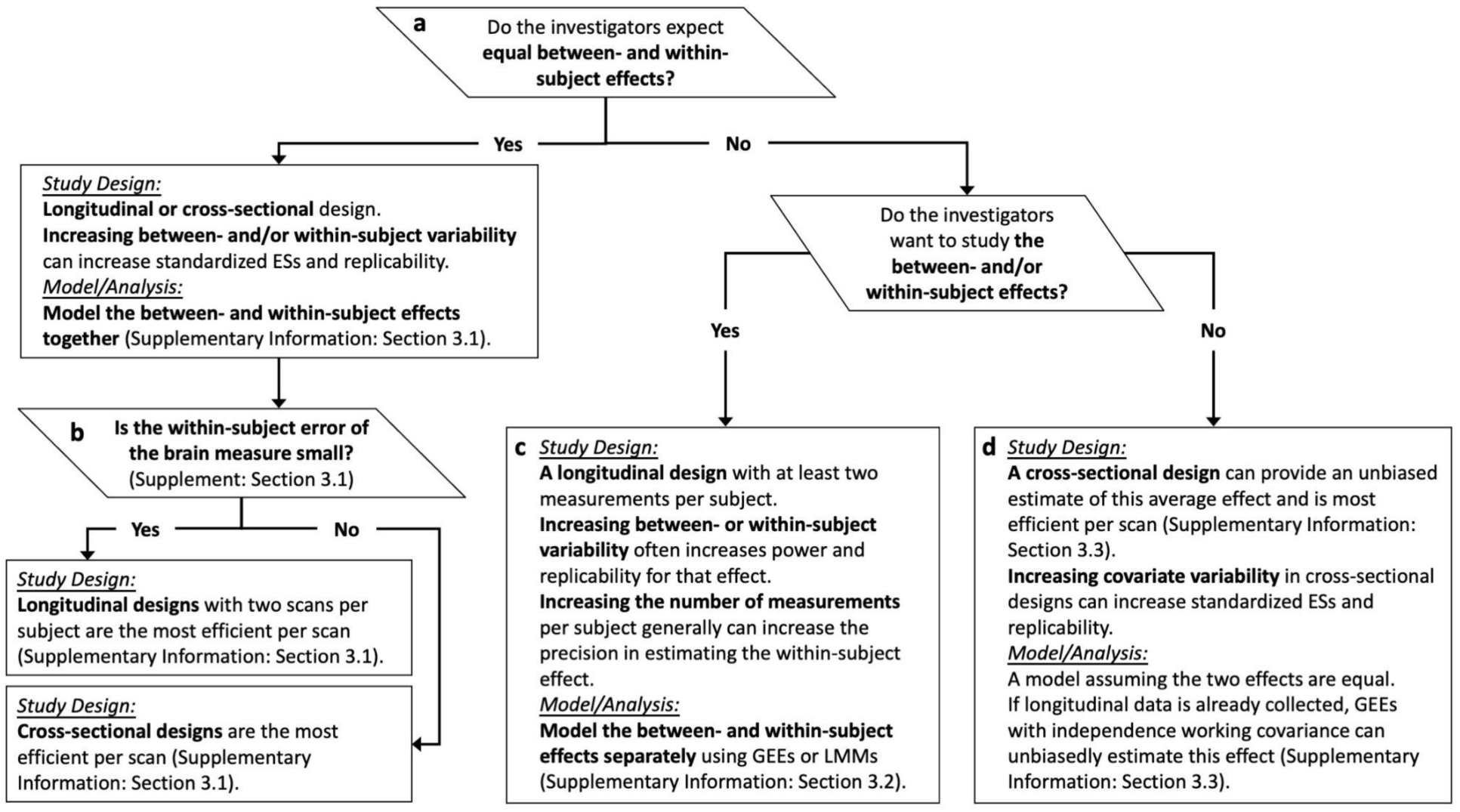
Optimal study design and analysis depends on characteristics of the hypothesized association(s). (**a**) Visualization can be performed in pilot or study data to evaluate this assumption as in Fig. S12 (Supplementary Information: Section 3). (**b**) If the between- and within-subject effects are hypothesized to be equal, either a cross-sectional or longitudinal design can be applied, but the efficiency per scan depends on the size of the within-subject error of the brain measure; pilot/study data can be used to evaluate this question (Supplementary Information: Section 3.1). (**c**) If estimating the between- and within-subject effects separately, a longitudinal design is required and common longitudinal data analysis tools such as generalized estimating equations (GEEs) and linear mixed models (LMMs) with separate between- and within-subject effects are required to unbiasedly estimate these effects (see Supplement: Section 3.2). (**d**) If there are different between- and within-subject effects, the investigators may still use a model to target the average effect (i.e., a weighted average of the underlying between- and within-subject effects) if they have cross-sectional data, or if they want results from a longitudinal study that are consistent for the same biological effect as cross-sectional studies. For longitudinal studies, a GEE with independence working covariance structure targets the same average effect as the cross-sectional model, but it is less statistically efficient than the cross-sectional model (see Supplement: Section 3.3). All recommendations are based on the empirical findings in the paper and the theory for exchangeable covariance longitudinal linear models in the Supplementary Information.

### Extended Data Tables

**Table S1.**
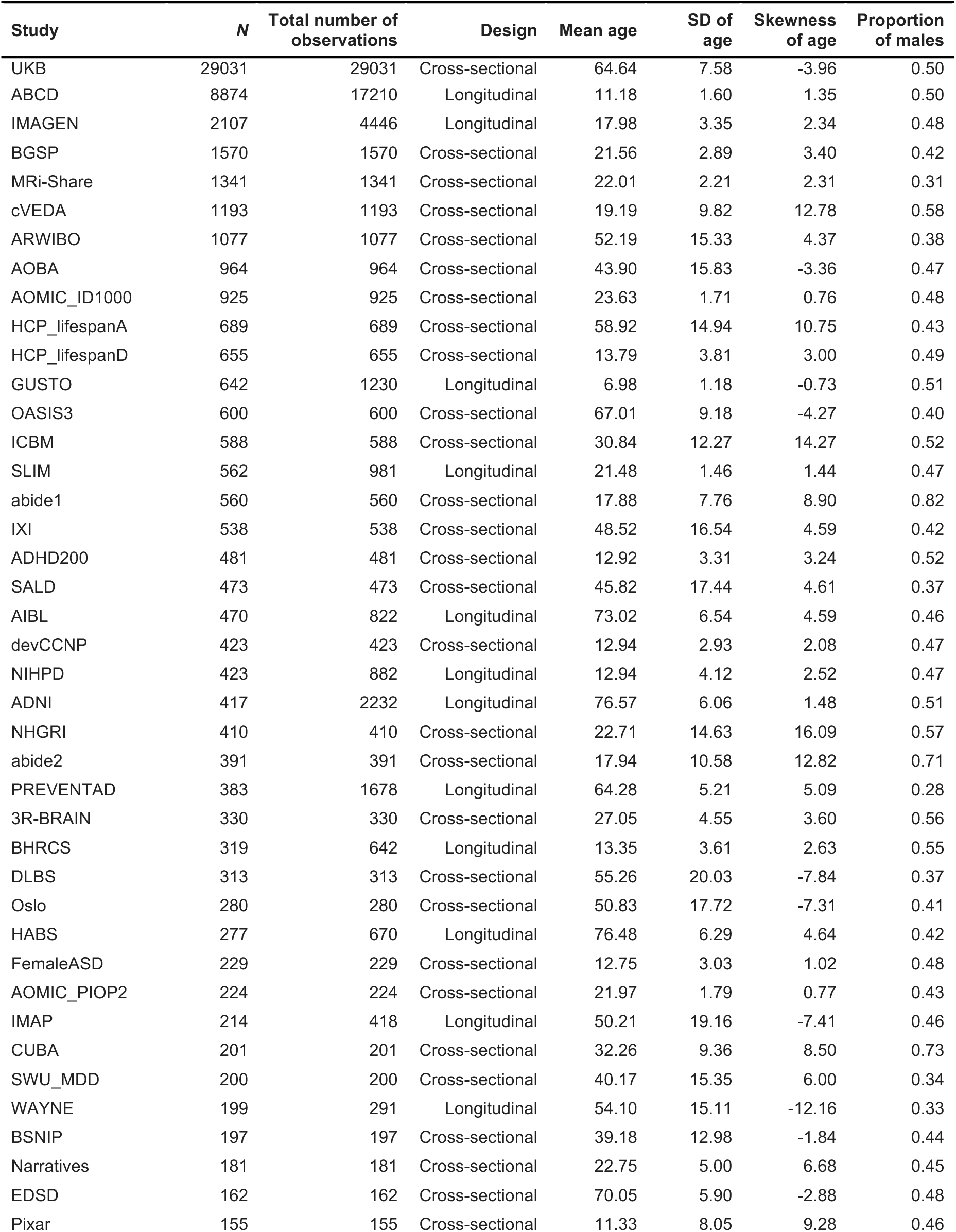
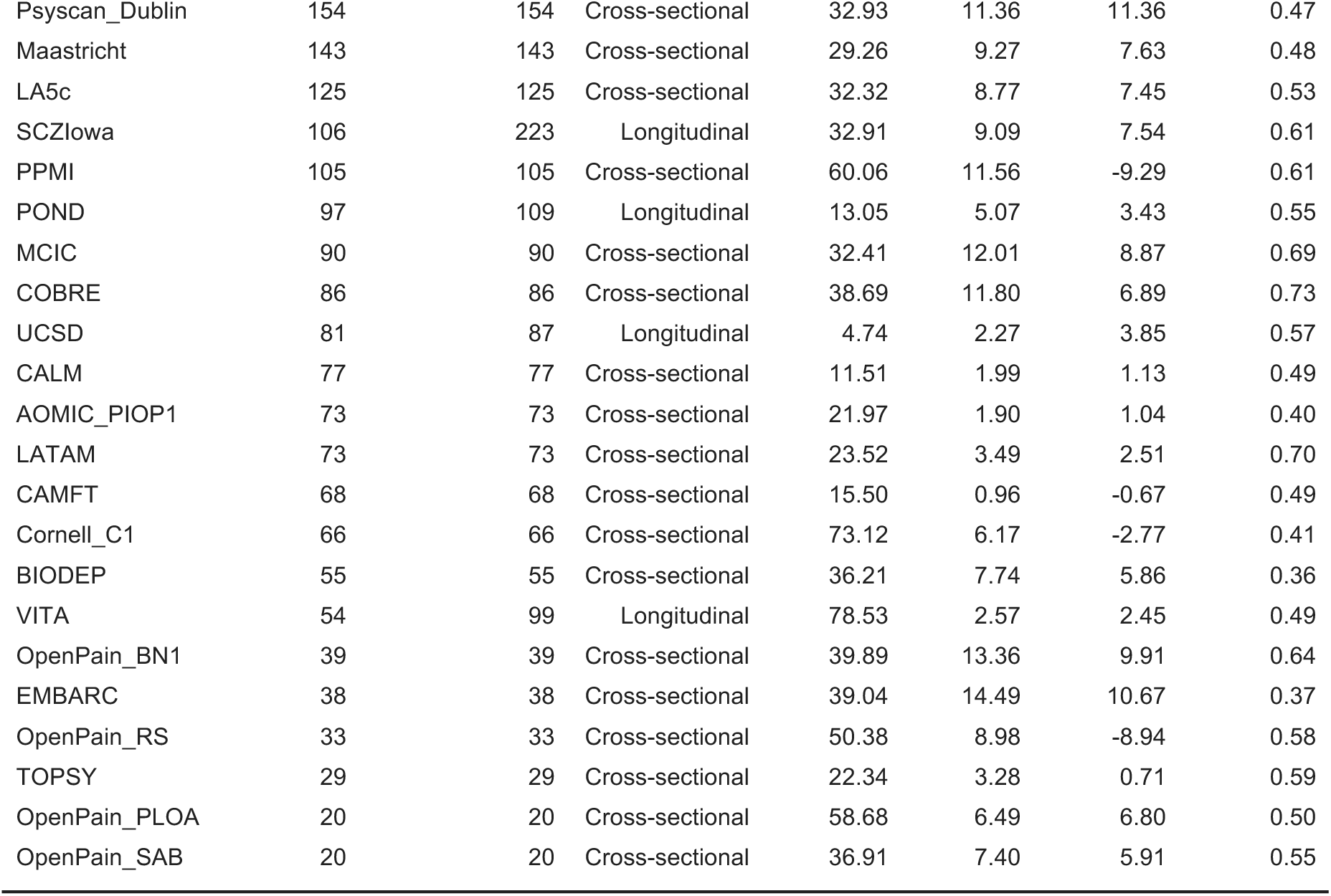
The descriptive summary of the 63 neuroimaging datasets used in the meta-analyses for the total gray matter volume (GMV), subcortical gray matter volume (sGMV) and white matter volume (WMV).

**Table S2.**
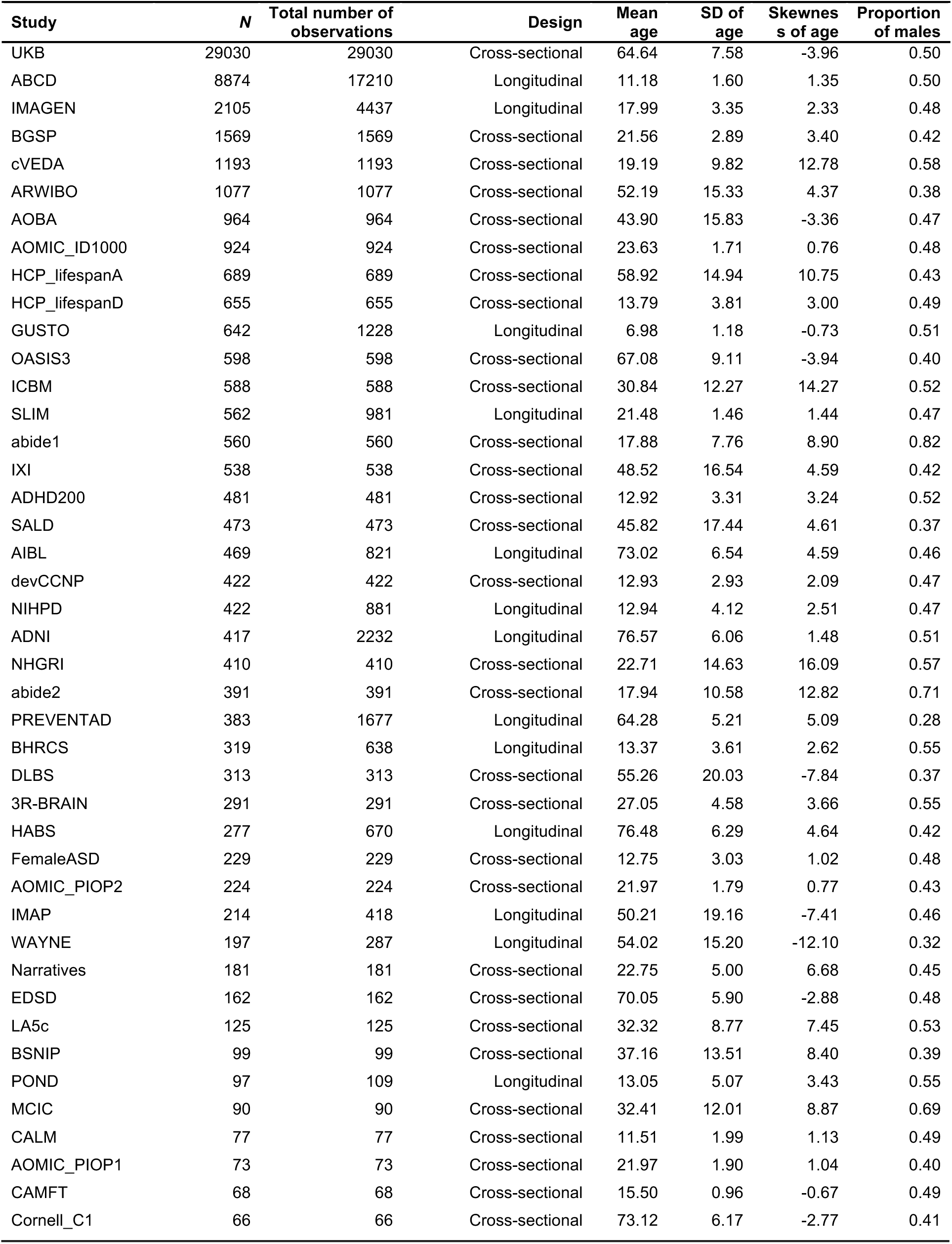
Descriptive summary table for the 43 studies used in the meta-analyses for mean cortical thickness (CT), regional gray matter volume (GMV), and regional CT.

**Table S3.**
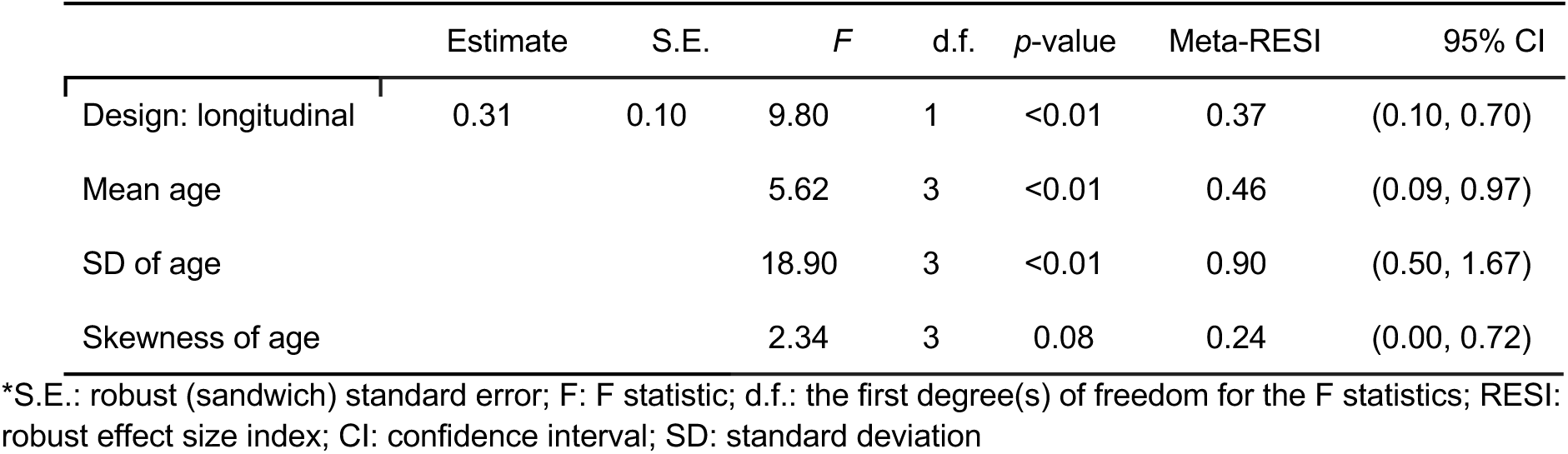
Meta-analysis results for the RESI for age on total gray matter volume (GMV). The residual degrees of freedom are 52. The regression model was weighted by the inverse standard errors of the RESIs.

|  | Estimate | S.E. | *F* | d.f. | *p*-value | Meta-RESI | 95% CI |
| --- | --- | --- | --- | --- | --- | --- | --- |
| Design: longitudinal | 0.31 | 0.10 | 9.80 | 1 | <0.01 | 0.37 | (0.10, 0.70) |
| Mean age |  |  | 5.62 | 3 | <0.01 | 0.46 | (0.09, 0.97) |
| SD of age |  |  | 18.90 | 3 | <0.01 | 0.90 | (0.50, 1.67) |
| Skewness of age |  |  | 2.34 | 3 | 0.08 | 0.24 | (0.00, 0.72) |

\*S.E.: robust (sandwich) standard error; *F*: *F* statistic; d.f.: the first degree(s) of freedom for the *F* statistics; RESI: robust effect size index; CI: confidence interval; SD: standard deviation

**Table S4.** Meta-analysis results for the RESI for age on total subcortical gray matter volume (sGMV). The residual degrees of freedom are 52. The regression model is weighted by the inverse standard errors of the RESIs.

|  | Estimate | S.E. | *F* | d.f. | *p*-value | Meta-RESI | 95% CI |
| --- | --- | --- | --- | --- | --- | --- | --- |
| Design: longitudinal | 0.23 | 0.11 | 4.92 | 1 | 0.03 | 0.24 | (0.00, 0.78) |
| Mean age |  |  | 1.61 | 3 | 0.20 | 0.16 | (0.00, 0.67) |
| SD of age |  |  | 6.98 | 3 | <0.01 | 0.52 | (0.00, 1.18) |
| Skewness of age |  |  | 1.27 | 3 | 0.29 | 0.10 | (0.00, 0.63) |

\*S.E.: robust (sandwich) standard error; *F*: *F* statistic; d.f.: the first degree(s) of freedom for the *F* statistics; RESI: robust ES index; CI: confidence interval; SD: standard deviation

**Table S5.** Meta-analysis results for the RESI for age on total white matter volume (WMV). The residual degrees of freedom are 52. The regression model is weighted by the inverse standard errors of the RESIs.

|  | Estimate | S.E. | *F* | d.f. | *p*-value | Meta-RESI | 95% CI |
| --- | --- | --- | --- | --- | --- | --- | --- |
| Design: longitudinal | 0.42 | 0.13 | 11.11 | 1 | <0.01 | 0.39 | (0.21, 0.88) |
| Mean age |  |  | 7.50 | 3 | <0.01 | 0.54 | (0.33, 1.39) |
| SD of age |  |  | 1.19 | 3 | 0.32 | 0.08 | (0.00, 0.75) |
| Skewness of age |  |  | 0.30 | 3 | 0.83 | 0.00 | (0.00, 0.50) |

\*S.E.: robust (sandwich) standard error; *F*: *F* statistic; d.f.: the first degree(s) of freedom for the *F* statistics; RESI: robust ES index; CI: confidence interval; SD: standard deviation

**Table S6.** Meta-analysis results for the RESI for age on mean cortical thickness (CT). The residual degrees of freedom are 32. The regression model is weighted by the inverse standard errors of the RESIs.

|  | Estimate | S.E. | *F* | d.f. | *p*-value | Meta-RESI | 95% CI |
| --- | --- | --- | --- | --- | --- | --- | --- |
| Design: longitudinal | 0.25 | 0.19 | 1.79 | 1 | 0.19 | 0.13 | (0.00, 0.82) |
| Mean age |  |  | 2.30 | 3 | 0.10 | 0.28 | (0.00, 1.26) |
| SD of age |  |  | 4.27 | 3 | 0.01 | 0.46 | (0.10, 1.55) |
| Skewness of age |  |  | 0.15 | 3 | 0.93 | 0.00 | (0.00, 0.67) |

\*S.E.: robust (sandwich) standard error; *F*: *F* statistic; d.f.: the first degree(s) of freedom for the *F* statistics; RESI: robust ES index; CI: confidence interval; SD: standard deviation

**Table S7.** Meta-analysis results for the RESI for sex on total gray matter volume (GMV). The residual degrees of freedom are 52. The regression model is weighted by the inverse standard errors of the RESIs.

|  | Estimate | S.E. | *F* | d.f. | *p*-value | Meta-RESI | 95% CI |
| --- | --- | --- | --- | --- | --- | --- | --- |
| Design: longitudinal | -0.03 | 0.11 | 0.06 | 1 | 0.80 | 0.00 | (0.00, 0.31) |
| Mean age |  |  | 0.45 | 3 | 0.72 | 0.00 | (0.00, 0.71) |
| SD of age |  |  | 0.40 | 3 | 0.75 | 0.00 | (0.00, 0.50) |
| Proportion of male |  |  | 0.18 | 3 | 0.91 | 0.00 | (0.00, 0.49) |

\*S.E.: robust (sandwich) standard error; *F*: *F* statistic; d.f.: the first degree(s) of freedom for the *F* statistics; RESI: robust ES index; CI: confidence interval; SD: standard deviation

**Table S8:** The meta-analysis results for the RESI for sex on total subcortical gray matter volume (sGMV). The residual degrees of freedom are 52. The regression model is weighted by the inverse standard errors of the RESIs.

|  | Estimate | S.E. | *F* | d.f. | *p*-value | Meta-RESI | 95% CI |
| --- | --- | --- | --- | --- | --- | --- | --- |
| Design: longitudinal | -0.07 | 0.12 | 0.27 | 1 | 0.60 | 0 | (0, 0.32) |
| Mean age |  |  | 0.60 | 3 | 0.62 | 0 | (0, 0.69) |
| SD of age |  |  | 0.59 | 3 | 0.62 | 0 | (0, 0.49) |
| Proportion of male |  |  | 0.12 | 3 | 0.95 | 0 | (0, 0.39) |

\*S.E.: robust (sandwich) standard error; *F*: *F* statistic; d.f.: the first degree(s) of freedom for the *F* statistics; RESI: robust ES index; CI: confidence interval; SD: standard deviation

**Table S9.**
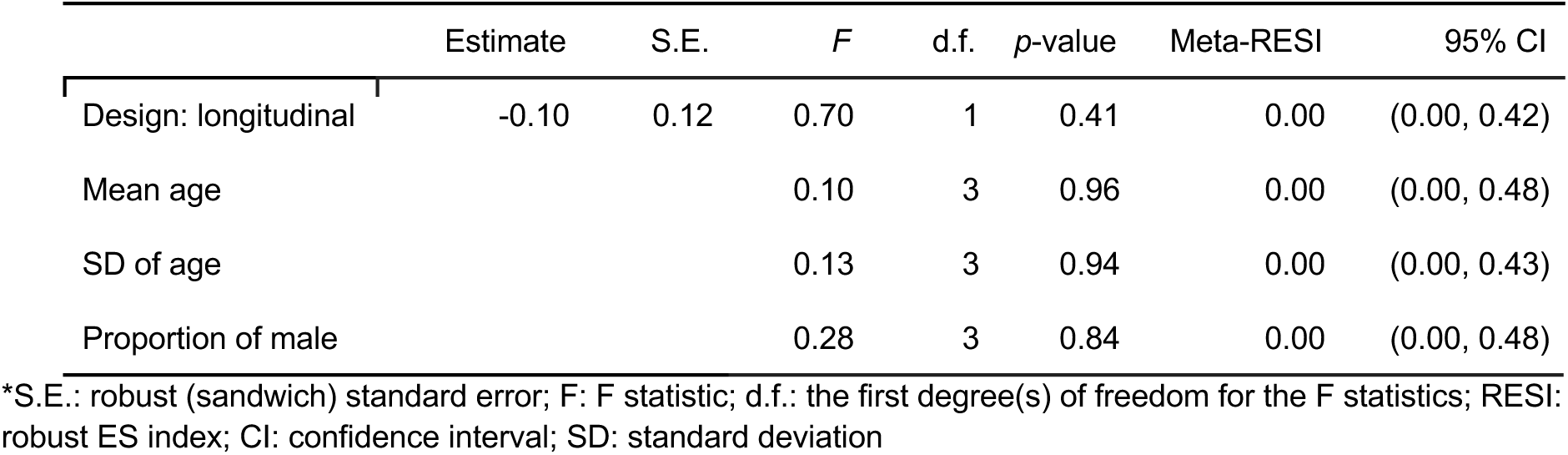
The meta-analysis results for the RESI for sex on total white matter volume (WMV). The residual degrees of freedom are 52. The regression model is weighted by the inverse standard errors of the RESIs.

**Table S10.**
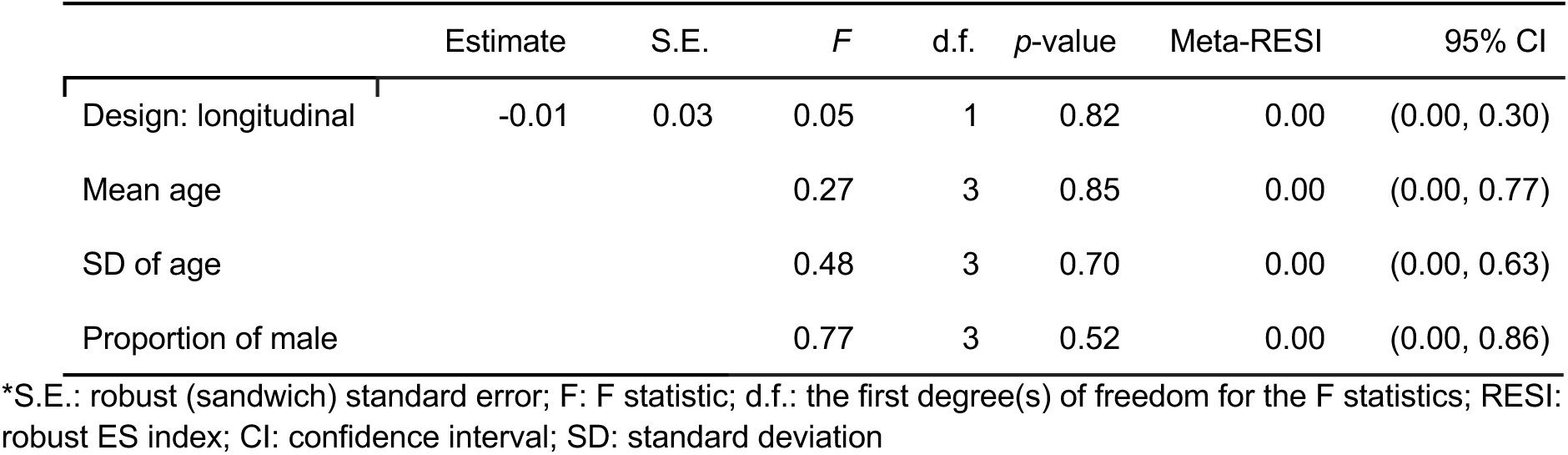
The meta-analysis results for the RESI for sex on mean cortical thickness (CT). The residual degrees of freedom are 32. The regression model is weighted by the inverse standard errors of the RESIs.

|  | Estimate | S.E. | *F* | d.f. | *p*-value | Meta-RESI | 95% CI |
| --- | --- | --- | --- | --- | --- | --- | --- |
| Design: longitudinal | -0.01 | 0.03 | 0.05 | 1 | 0.82 | 0.00 | (0.00, 0.30) |
| Mean age |  |  | 0.27 | 3 | 0.85 | 0.00 | (0.00, 0.77) |
| SD of age |  |  | 0.48 | 3 | 0.70 | 0.00 | (0.00, 0.63) |
| Proportion of male |  |  | 0.77 | 3 | 0.52 | 0.00 | (0.00, 0.86) |

\*S.E.: robust (sandwich) standard error; *F*: *F* statistic; d.f.: the first degree(s) of freedom for the *F* statistics; RESI: robust ES index; CI: confidence interval; SD: standard deviation

**Table S11.**
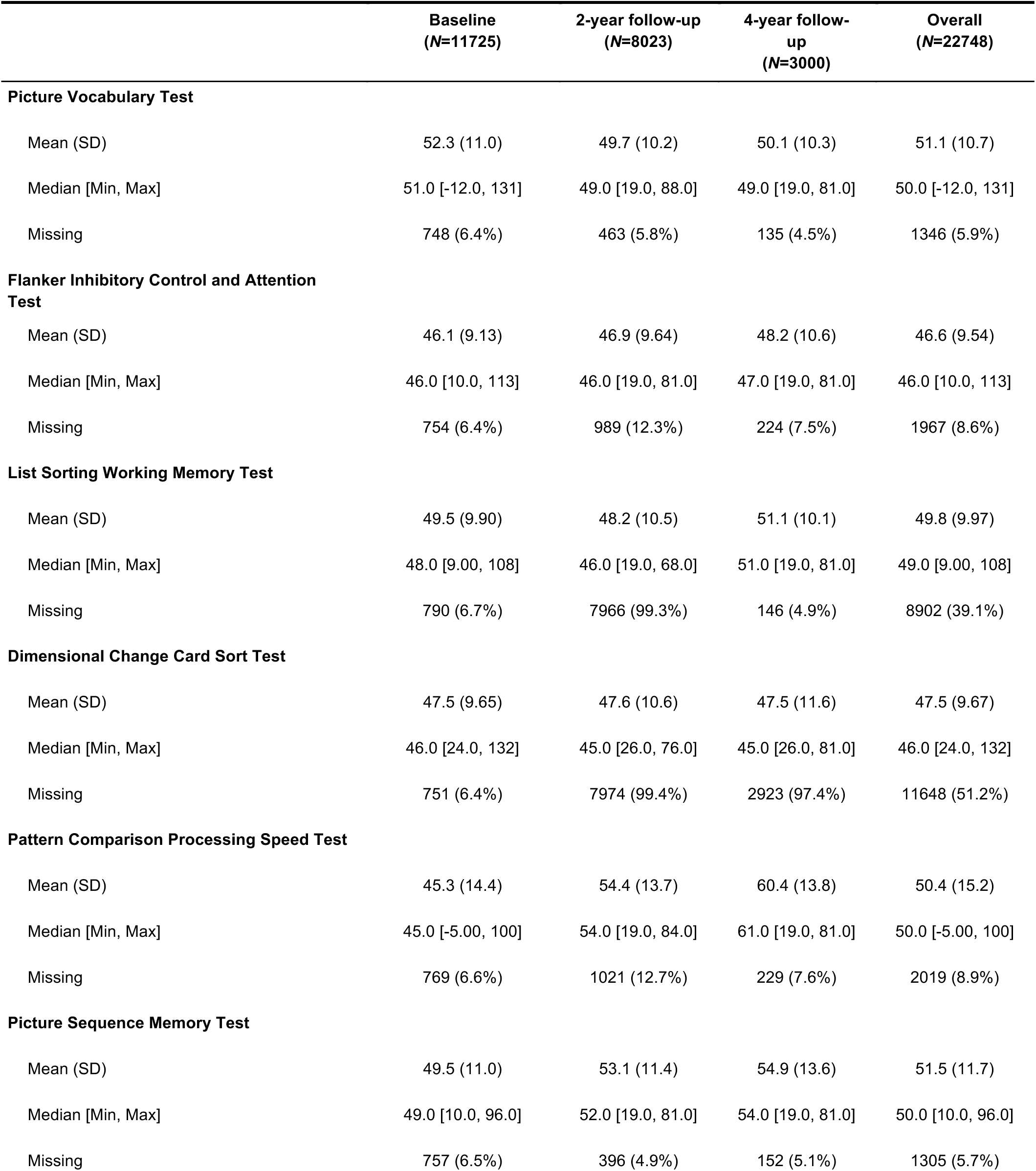
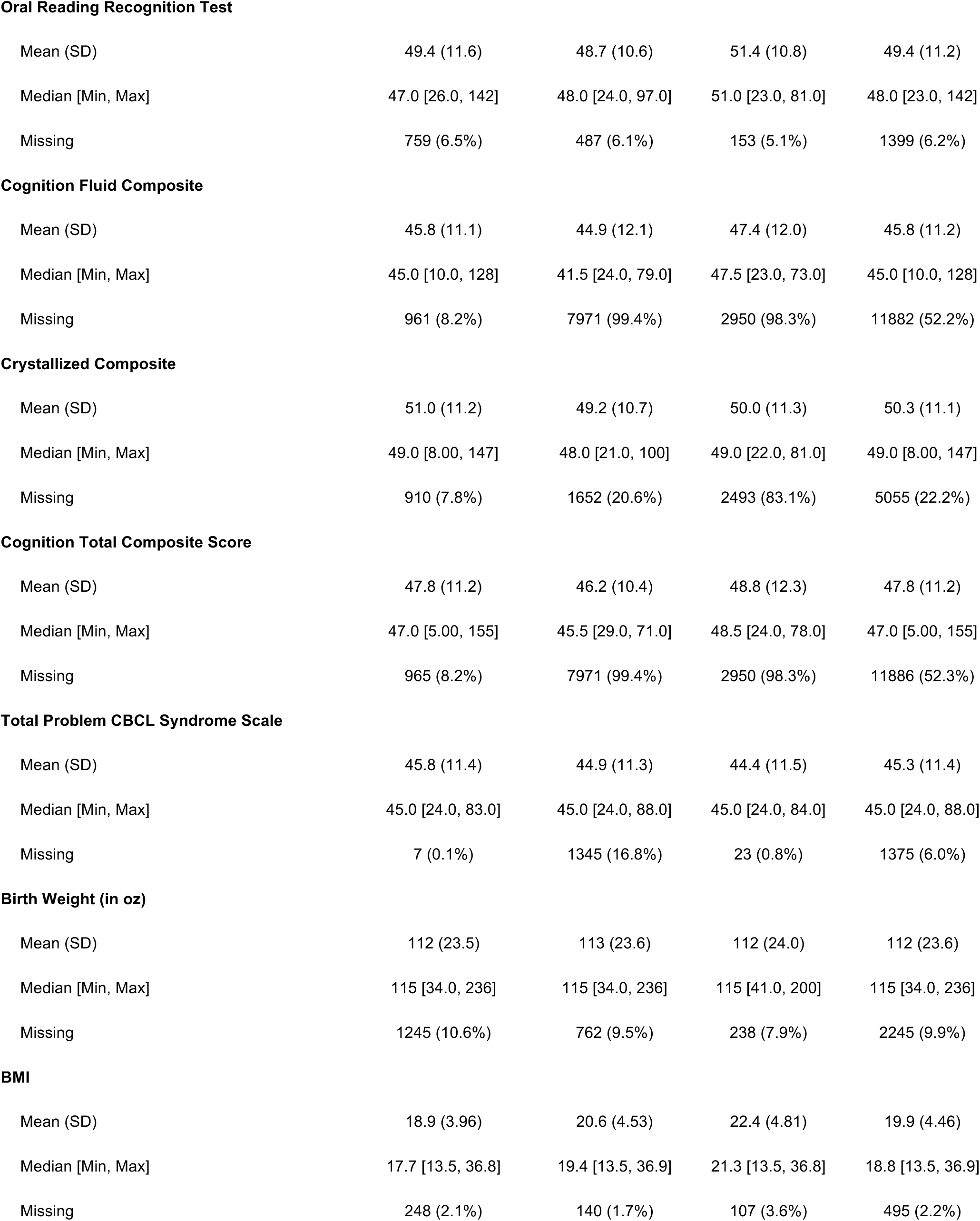
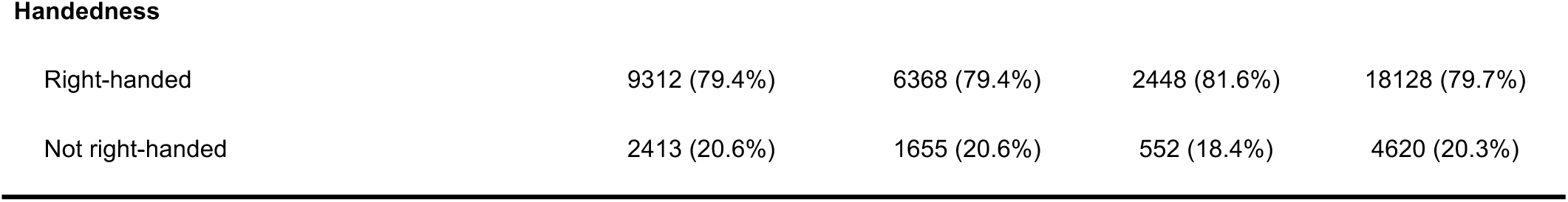
The descriptive summary of the non-brain measures in ABCD study used for the structural brain outcome analyses. The records with missing global and regional brain measures and the subjects with missing baseline records are removed.

|  | Baseline<br>(N=11725) | 2-year follow-up<br>(N=8023) | 4-year follow-up<br>(N=3000) | Overall<br>(N=22748) |
| --- | --- | --- | --- | --- |
| **Picture Vocabulary Test** |  |  |  |  |
| Mean (SD) | 52.3 (11.0) | 49.7 (10.2) | 50.1 (10.3) | 51.1 (10.7) |
| Median [Min, Max] | 51.0 [-12.0, 131] | 49.0 [19.0, 88.0] | 49.0 [19.0, 81.0] | 50.0 [-12.0, 131] |
| Missing | 748 (6.4%) | 463 (5.8%) | 135 (4.5%) | 1346 (5.9%) |
| **Flanker Inhibitory Control and Attention Test** |  |  |  |  |
| Mean (SD) | 46.1 (9.13) | 46.9 (9.64) | 48.2 (10.6) | 46.6 (9.54) |
| Median [Min, Max] | 46.0 [10.0, 113] | 46.0 [19.0, 81.0] | 47.0 [19.0, 81.0] | 46.0 [10.0, 113] |
| Missing | 754 (6.4%) | 989 (12.3%) | 224 (7.5%) | 1967 (8.6%) |
| **List Sorting Working Memory Test** |  |  |  |  |
| Mean (SD) | 49.5 (9.90) | 48.2 (10.5) | 51.1 (10.1) | 49.8 (9.97) |
| Median [Min, Max] | 48.0 [9.00, 108] | 46.0 [19.0, 68.0] | 51.0 [19.0, 81.0] | 49.0 [9.00, 108] |
| Missing | 790 (6.7%) | 7966 (99.3%) | 146 (4.9%) | 8902 (39.1%) |
| **Dimensional Change Card Sort Test** |  |  |  |  |
| Mean (SD) | 47.5 (9.65) | 47.6 (10.6) | 47.5 (11.6) | 47.5 (9.67) |
| Median [Min, Max] | 46.0 [24.0, 132] | 45.0 [26.0, 76.0] | 45.0 [26.0, 81.0] | 46.0 [24.0, 132] |
| Missing | 751 (6.4%) | 7974 (99.4%) | 2923 (97.4%) | 11648 (51.2%) |
| **Pattern Comparison Processing Speed Test** |  |  |  |  |
| Mean (SD) | 45.3 (14.4) | 54.4 (13.7) | 60.4 (13.8) | 50.4 (15.2) |
| Median [Min, Max] | 45.0 [-5.00, 100] | 54.0 [19.0, 84.0] | 61.0 [19.0, 81.0] | 50.0 [-5.00, 100] |
| Missing | 769 (6.6%) | 1021 (12.7%) | 229 (7.6%) | 2019 (8.9%) |
| **Picture Sequence Memory Test** |  |  |  |  |
| Mean (SD) | 49.5 (11.0) | 53.1 (11.4) | 54.9 (13.6) | 51.5 (11.7) |
| Median [Min, Max] | 49.0 [10.0, 96.0] | 52.0 [19.0, 81.0] | 54.0 [19.0, 81.0] | 50.0 [10.0, 96.0] |
| Missing | 757 (6.5%) | 396 (4.9%) | 152 (5.1%) | 1305 (5.7%) |
| **Oral Reading Recognition Test** |  |  |  |  |
| Mean (SD) | 49.4 (11.6) | 48.7 (10.6) | 51.4 (10.8) | 49.4 (11.2) |
| Median [Min, Max] | 47.0 [26.0, 142] | 48.0 [24.0, 97.0] | 51.0 [23.0, 81.0] | 48.0 [23.0, 142] |
| Missing | 759 (6.5%) | 487 (6.1%) | 153 (5.1%) | 1399 (6.2%) |
| **Cognition Fluid Composite** |  |  |  |  |
| Mean (SD) | 45.8 (11.1) | 44.9 (12.1) | 47.4 (12.0) | 45.8 (11.2) |
| Median [Min, Max] | 45.0 [10.0, 128] | 41.5 [24.0, 79.0] | 47.5 [23.0, 73.0] | 45.0 [10.0, 128] |
| Missing | 961 (8.2%) | 7971 (99.4%) | 2950 (98.3%) | 11882 (52.2%) |
| **Crystallized Composite** |  |  |  |  |
| Mean (SD) | 51.0 (11.2) | 49.2 (10.7) | 50.0 (11.3) | 50.3 (11.1) |
| Median [Min, Max] | 49.0 [8.00, 147] | 48.0 [21.0, 100] | 49.0 [22.0, 81.0] | 49.0 [8.00, 147] |
| Missing | 910 (7.8%) | 1652 (20.6%) | 2493 (83.1%) | 5055 (22.2%) |
| **Cognition Total Composite Score** |  |  |  |  |
| Mean (SD) | 47.8 (11.2) | 46.2 (10.4) | 48.8 (12.3) | 47.8 (11.2) |
| Median [Min, Max] | 47.0 [5.00, 155] | 45.5 [29.0, 71.0] | 48.5 [24.0, 78.0] | 47.0 [5.00, 155] |
| Missing | 965 (8.2%) | 7971 (99.4%) | 2950 (98.3%) | 11886 (52.3%) |
| **Total Problem CBCL Syndrome Scale** |  |  |  |  |
| Mean (SD) | 45.8 (11.4) | 44.9 (11.3) | 44.4 (11.5) | 45.3 (11.4) |
| Median [Min, Max] | 45.0 [24.0, 83.0] | 45.0 [24.0, 88.0] | 45.0 [24.0, 84.0] | 45.0 [24.0, 88.0] |
| Missing | 7 (0.1%) | 1345 (16.8%) | 23 (0.8%) | 1375 (6.0%) |
| **Birth Weight (in oz)** |  |  |  |  |
| Mean (SD) | 112 (23.5) | 113 (23.6) | 112 (24.0) | 112 (23.6) |
| Median [Min, Max] | 115 [34.0, 236] | 115 [34.0, 236] | 115 [41.0, 200] | 115 [34.0, 236] |
| Missing | 1245 (10.6%) | 762 (9.5%) | 238 (7.9%) | 2245 (9.9%) |
| **BMI** |  |  |  |  |
| Mean (SD) | 18.9 (3.96) | 20.6 (4.53) | 22.4 (4.81) | 19.9 (4.46) |
| Median [Min, Max] | 17.7 [13.5, 36.8] | 19.4 [13.5, 36.9] | 21.3 [13.5, 36.8] | 18.8 [13.5, 36.9] |
| Missing | 248 (2.1%) | 140 (1.7%) | 107 (3.6%) | 495 (2.2%) |
| **Handedness** |  |  |  |  |
| Right-handed | 9312 (79.4%) | 6368 (79.4%) | 2448 (81.6%) | 18128 (79.7%) |
| Not right-handed | 2413 (20.6%) | 1655 (20.6%) | 552 (18.4%) | 4620 (20.3%) |

**Table S12.** The descriptive summary of the non-brain measures in the subset of ABCD study used for the functional connectivity (FC) outcome analyses. Only the subjects with both baseline and 2-year follow-up are included.

|  | Baseline<br>(N=1192) | 2-year follow-up<br>(N=1192) | Overall<br>(N=2384) |
| --- | --- | --- | --- |
| **Picture Vocabulary Test** |  |  |  |
| Mean (SD) | 52.8 (11.2) | 50.2 (10.4) | 51.5 (10.9) |
| Median [Min, Max] | 52.0 [14.0, 112] | 49.0 [19.0, 88.0] | 50.0 [14.0, 112] |
| Missing | 62 (5.2%) | 75 (6.3%) | 137 (5.7%) |
| **Flanker Inhibitory Control and Attention Test** |  |  |  |
| Mean (SD) | 46.0 (9.17) | 46.9 (9.40) | 46.5 (9.29) |
| Median [Min, Max] | 45.5 [10.0, 113] | 46.0 [19.0, 81.0] | 46.0 [10.0, 113] |
| Missing | 62 (5.2%) | 130 (10.9%) | 192 (8.1%) |
| **List Sorting Working Memory Test** |  |  |  |
| Mean (SD) | 50.1 (9.55) | 52.5 (12.0) | 50.1 (9.58) |
| Median [Min, Max] | 49.0 [26.0, 91.0] | 48.0 [34.0, 68.0] | 49.0 [26.0, 91.0] |
| Missing | 67 (5.6%) | 1181 (99.1%) | 1248 (52.3%) |
| **Dimensional Change Card Sort Test** |  |  |  |
| Mean (SD) | 47.9 (9.78) | 47.1 (14.4) | 47.9 (9.81) |
| Median [Min, Max] | 46.0 [26.0, 98.0] | 41.0 [33.0, 71.0] | 46.0 [26.0, 98.0] |
| Missing | 63 (5.3%) | 1184 (99.3%) | 1247 (52.3%) |
| **Pattern Comparison Processing Speed Test** |  |  |  |
| Mean (SD) | 45.9 (13.9) | 54.4 (13.3) | 50.0 (14.3) |
| Median [Min, Max] | 46.0 [-2.00, 88.0] | 54.0 [19.0, 81.0] | 50.0 [-2.00, 88.0] |
| Missing | 64 (5.4%) | 137 (11.5%) | 201 (8.4%) |
| **Picture Sequence Memory Test** |  |  |  |
| Mean (SD) | 49.8 (11.2) | 53.6 (11.4) | 51.7 (11.4) |
| Median [Min, Max] | 49.0 [15.0, 96.0] | 52.0 [19.0, 81.0] | 51.0 [15.0, 96.0] |
| Missing | 63 (5.3%) | 66 (5.5%) | 129 (5.4%) |
| **Oral Reading Recognition Test** |  |  |  |
| Mean (SD) | 49.6 (11.3) | 49.4 (10.9) | 49.5 (11.1) |
| Median [Min, Max] | 48.0 [28.0, 118] | 48.0 [24.0, 97.0] | 48.0 [24.0, 118] |
| Missing | 64 (5.4%) | 78 (6.5%) | 142 (6.0%) |
| **Cognition Fluid Composite** |  |  |  |
| Mean (SD) | 46.4 (11.2) | 49.8 (17.9) | 46.5 (11.3) |
| Median [Min, Max] | 46.0 [20.0, 113] | 47.0 [29.0, 79.0] | 46.0 [20.0, 113] |
| Missing | 81 (6.8%) | 1182 (99.2%) | 1263 (53.0%) |
| **Crystallized Composite** |  |  |  |
| Mean (SD) | 51.3 (11.2) | 49.9 (10.9) | 50.7 (11.1) |
| Median [Min, Max] | 50.0 [22.0, 110] | 49.0 [23.0, 100] | 49.0 [22.0, 110] |
| Missing | 80 (6.7%) | 201 (16.9%) | 281 (11.8%) |
| **Cognition Total Composite Score** |  |  |  |
| Mean (SD) | 48.4 (11.2) | 50.7 (12.8) | 48.4 (11.2) |
| Median [Min, Max] | 48.0 [18.0, 127] | 53.5 [29.0, 70.0] | 48.0 [18.0, 127] |
| Missing | 84 (7.0%) | 1182 (99.2%) | 1266 (53.1%) |
| **Total Problem CBCL Syndrome Scale** |  |  |  |
| Mean (SD) | 45.4 (11.1) | 44.3 (11.0) | 44.9 (11.1) |
| Median [Min, Max] | 45.0 [24.0, 81.0] | 44.5 [24.0, 78.0] | 45.0 [24.0, 81.0] |
| Missing | 0 (0%) | 160 (13.4%) | 160 (6.7%) |
| **Birth Weight (in oz)** |  |  |  |
| Mean (SD) | 112 (23.8) | 112 (23.8) | 112 (23.8) |
| Median [Min, Max] | 114 [35.0, 208] | 114 [35.0, 208] | 114 [35.0, 208] |
| Missing | 121 (10.2%) | 116 (9.7%) | 237 (9.9%) |
| **BMI** |  |  |  |
| Mean (SD) | 18.0 (3.27) | 19.5 (3.67) | 18.7 (3.56) |
| Median [Min, Max] | 17.1 [13.2, 30.7] | 18.7 [13.2, 32.2] | 17.9 [13.2, 32.2] |
| Missing | 26 (2.2%) | 21 (1.8%) | 47 (2.0%) |
| **Handedness** |  |  |  |
| Right-handed | 968 (81.2%) | 968 (81.2%) | 1936 (81.2%) |
| Not right-handed | 224 (18.8%) | 224 (18.8%) | 448 (18.8%) |

**Table S13.**
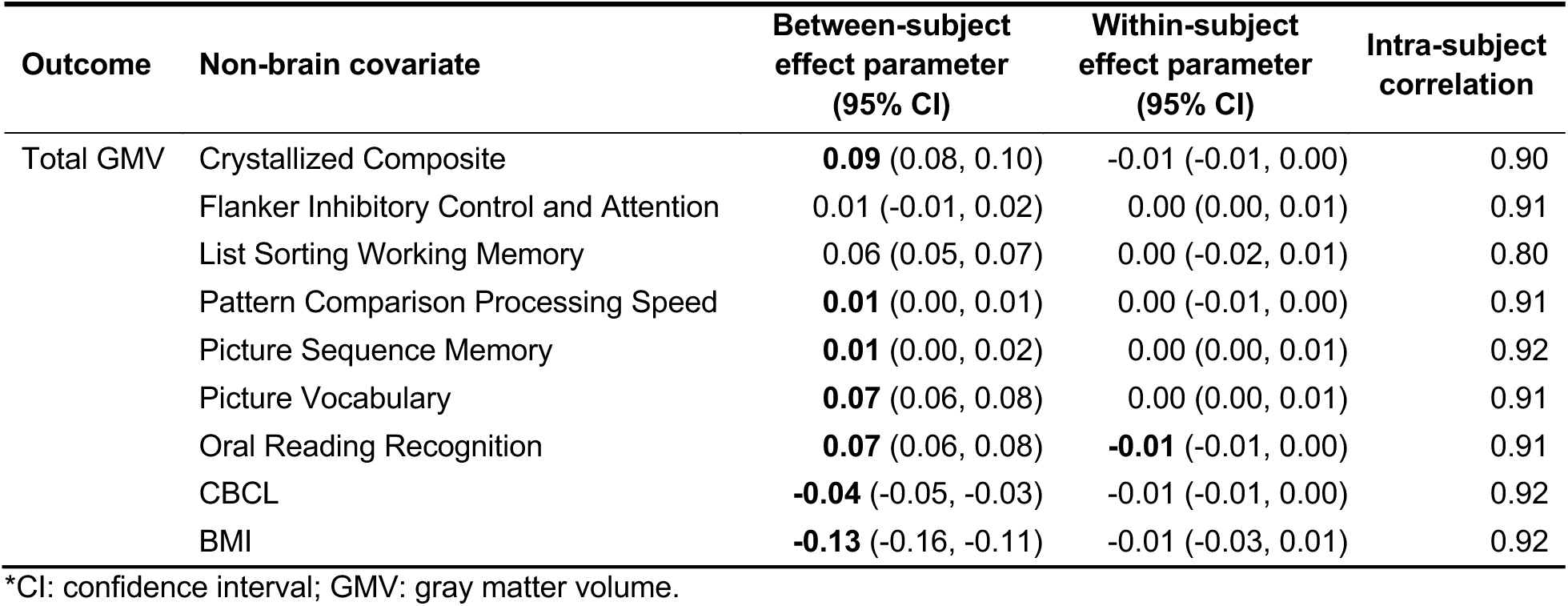
The estimated cross-sectional and longitudinal parameters of the non-brain covariates on total gray matter volume (GMV), respectively, in the ABCD study. See Methods for more details about the estimation of the between- and within-subject effect parameters. The parameter estimates that are significantly different from zero are in bold. The intra-subject correlation of the outcome total GMV is estimated from the GEEs (generalized estimating equations) with an exchangeable correlation structure used for the modeling.

