## Supplementary Information for "Study design features increase replicability in cross-sectional and longitudinal brain-wide association studies"

### 1 Robust Effect Size Index (RESI) in Longitudinal Analysis

Standardized effect size (ES) indices are measures quantifying the strength of association between a variable and an outcome of interest. They are unaffected by the sample size and play an important role in power analyses, sample size planning, meta-analyses, and replicability[1–3]. Reporting ES estimates with confidence intervals (CIs) can simultaneously communicate the strength of evidence (sample size independent) as well as the precision in the observed evidence (sample size dependent).

We recently proposed a robust ES index (RESI) in the setting of cross-sectional data analysis [4, 5], which is a standardized parameter describing the deviation of the true parameter value from a reference value. It has several advantages over previously proposed indices [1, 6–9] because: it is widely applicable to many types of data since it is constructed from M-estimators, which are generally defined; it is robust to possible model misspecification as it can use a heteroskedasticity consistent covariance estimator [10]; it can accommodate the existence of nuisance covariates/parameters. We also provided a consistent estimator and a valid CI estimation method for the RESI [4, 5].

The correlation between observations makes the analysis of longitudinal data complicated. Generalized estimating equations (GEEs) [11], which estimate marginal associations, are commonly used in the analysis of longitudinal data. There are at least two advantages of GEEs: robustness to misspecification of the correlation structure of the repeatedly measured data; and computational simplicity. Technically, standardized ES indices designed for cross-sectional analysis can still be applied in longitudinal analysis, but the resulting standardized ES will incorporate any benefits of the longitudinal designs (i.e., the repeated measurements). The additional benefit of the longitudinal design introduces a systematic difference in the standardized ES estimates not only between longitudinal and cross-sectional data designs, but also between the longitudinal designs with different numbers of measurements per subject. Such a systematic difference compromises the comparability of the standardized ESs between studies with different designs. If we would like to compare the standardized ESs from studies studying similar scientific questions, but using different designs, it would be ideal to put them on the same "scale". To our knowledge, there is no ES index defined for longitudinal analysis models with this purpose.

In this section, we modify the RESI for longitudinal data analysis (with the modification called the "cross-sectional" RESI or CS-RESI) based on GEEs to make it comparable between longitudinal and cross-sectional studies and propose an estimator and CI estimation method for it. We conduct simulation studies to evaluate the consistency of the estimator, the nominal coverage of the CI estimation method, and the comparability between the CS-RESI in longitudinal analysis and the previously proposed RESI in an equivalent cross-sectional analysis.

#### 1.1 RESI for Generalized Estimating Equations

Let $Y_{ij}$ denote the outcome variable and $X_{ij}$ be a vector of length $p$ explanatory variables (i.e., covariates) observed at time $t_{ij}$ for observation $j = 1, \dots, n_i$ from $i = 1, \dots, N$ independent subjects. Let $n = \sum_{i=1}^{N} n_i$ be the total number of measurements in the sample. The set of repeated measurements from subject $i$ is a vector of length $n_i$, $Y_i = (Y_{i1}, Y_{i2}, \dots, Y_{in_i})$. We can assume the marginal mean model to be

$$\mathbb{E}(Y_{ij}|X_{ij}) = \mu_{ij}(\theta) = g^{-1}\left(X_{ij}^T\theta\right),$$

where $\mathbb{E}(Y_{ij}|X_{ij})$ is the expectation of $Y_{ij}$ conditional on $X_{ij}$ and $g$ is a known link function [11]. $\theta = (\alpha, \beta) \in \Theta \subset \mathbb{R}^p$ are the model parameters, where $\alpha \in \mathbb{R}^{p_0}$ is a vector of nuisance parameters, $\beta \in \mathbb{R}^{p_1}$ is a vector of target parameter, and $p = p_0 + p_1$ which is the total number of parameters.

Let $V_i(\rho) = \mathrm{Var}(Y_i)$ denote the working covariance structure of $Y_i$, which depends on the marginal means and additional correlation parameter $\rho$. The estimating function used by GEEs for $\theta$ is given by

$$U_\theta(\theta,\rho) = \sum_{i=1}^{N} D_i^T V_i^{-1}(\rho)(Y_i - \mu_i), \tag{1}$$

where $D_i = (\partial\mu_i/\partial\theta)^T$ and $\mu_i = (\mu_{i1}, \mu_{i2}, \ldots, \mu_{in_i})$, is the marginal mean for subject $i$ for the $n_i$ observations [11].

For a given value of $\rho$, the solution to $U_\theta(\hat{\theta},\rho) = 0$ defines an estimator $\hat{\theta}$. Under mild regularity conditions [11], the estimator for $\theta$ obtained from (1) is consistent and

$$\sqrt{N}(\hat{\theta} - \theta) \xrightarrow{d} \mathrm{MVN}_p\left(0, \Sigma_\theta\right).$$

where

$$\begin{aligned}
\Sigma_\theta &= A^{-1}BA^{-1} \qquad (2)\\
A &= \lim_{N\to\infty} N^{-1} \sum_{i=1}^{N} D_i^T V_i^{-1} D_i\\
B &= \lim_{N\to\infty} N^{-1} \sum_{i=1}^{N} D_i^T V_i^{-1} \mathrm{Cov}(Y_i) V_i^{-1} D_i
\end{aligned}$$

If the correlation model is correctly specified, $\mathrm{Cov}(Y_i) = V_i$, then $\Sigma_\theta = A^{-1}$.

When $\Sigma_\theta$ is known, the Wald-type statistic testing the null hypothesis $H_0 : \beta = \beta_0$, where $\beta_0$ is the reference values, follows a chi-squared distribution with $p_1$ degrees of freedom and non-centrality parameter (NCP) $N(\beta - \beta_0)^T \Sigma_\beta^{-1}(\theta)(\beta - \beta_0)$,

$$T^2 = N(\hat{\beta} - \beta_0)^T \Sigma_\beta^{-1}(\theta)(\hat{\beta} - \beta_0) \sim \chi^2_{p_1}\left(N(\beta - \beta_0)^T \Sigma_\beta^{-1}(\theta)(\beta - \beta_0)\right),$$

where $\hat{\beta}$ is the estimator for $\beta$ and $\Sigma_\beta(\theta)$ is the asymptotic covariance matrix of $\sqrt{N}(\hat{\beta} - \beta)$.

##### 1.1.1 Longitudinal RESI (L-RESI)

The original definition of the RESI for cross-sectional analysis [4] is still applicable here, and is defined as the square root of the component of the NCP that is due to the deviation of $\beta$ from the reference values,

$$S = \sqrt{(\beta - \beta_0)^T \Sigma_\beta^{-1}(\beta - \beta_0)}. \tag{3}$$

We also defined a signed version of the RESI for univariate parameters $\beta$ [12]

$$S_{\mathrm{sgn}} = \beta/\Sigma_\beta^{1/2}, \tag{4}$$

where $\Sigma_\beta \in \mathbb{R}$. The absolute value of $S_{\mathrm{sgn}}$ is equal to $S$.

The previously defined estimators for (3) and (4) are consistent in this context by the theory described in previous work [4, 5]:

$$\hat{S} = \left(\max\left\{0, \frac{\hat{T}^2 - p_1}{N}\right\}\right)^{\frac{1}{2}} \text{ and}$$
$$\hat{S}_{\mathrm{sgn}} = \frac{\hat{T}}{\sqrt{N}}.$$

The RESI is independent of the sample size $N$ but dependent on the within-subject correlation. This can be seen through the dependence of (3) and (4) on $\text{Cov}(Y_i)$ through (2). The RESI in equation (3) quantifies the standardized ES over the entire follow-up, therefore is influenced by the study design such as the number of measurements, length of follow-up time, and correlation structure. Throughout this document, we refer to the RESI defined in the setting of cross-sectional analysis but applied in longitudinal analysis as longitudinal RESI or L-RESI to indicate that it implicitly incorporates the information of the longitudinal design. This is different from the main paper, where we use the term "RESI" for this value regardless of cross-sectional or longitudinal analysis.

##### 1.1.2 "Cross-sectional" RESI in Longitudinal Analysis

The systematic difference in the standardized ESs between cross-sectional and longitudinal studies makes comparing and aggregating cross-sectional and longitudinal studies more challenging in meta-analysis. In order to allow more consistent comparisons of standardized ES estimates across cross-sectional and longitudinal studies, we modify the definition of the RESI to let it represent the standardized ES as if the study was conducted cross-sectionally, but using the longitudinal data. As $\beta$ quantifies the marginal effect, it is equivalent in cross-sectional and longitudinal studies. Thus, only the variance in (3) is different between cross-sectional and longitudinal studies, so the target parameter is

$$S_{cs} = \sqrt{(\beta - \beta_0)^T \Sigma_{\beta_{cs}}^{-1} (\beta - \beta_0)}, \tag{5}$$

where $\Sigma_{\beta_{cs}}$ is the asymptotic covariance matrix of $\sqrt{N}(\hat{\beta} - \beta)$ from an equivalent cross-sectional study. We refer to this as "CS-RESI" in a longitudinal study.

To relate the equivalent cross-sectional study to the longitudinal study, we assume

$$L_{cs} = RL, \tag{6}$$

where $L = [Y,\ X]$ is an $n \times (p+1)$ matrix of the full longitudinal data including $n_i$ measurements for each subject $i$, $L_{cs}$ is an $N \times (p+1)$ matrix of covariates and outcome $Y$, and $R$ is an $N \times n$ random block sampling matrix that samples one row for each subject in the longitudinal study with equal probability. We assume that $R$ is independent of $L$, so that the data are missing completely at random.

If the mean model is correctly specified, estimation of the CS-RESI for GEEs boils down to obtaining a consistent estimator for $\Sigma_{\beta_{cs}}$ using the longitudinal data. For identity link regression models, we propose the use of the estimator

$$\hat{\Sigma}_{\beta_{cs}} = (N^{-1}X^T W X)^{-1}(N^{-1}X^T W \hat{\Delta} W X)(N^{-1}X^T W X)^{-1} := \hat{A}_{cs}^{-1}\hat{B}_{cs}\hat{A}_{cs}^{-1}, \tag{7}$$

where $W$ is a diagonal matrix that weights each observation by the inverse of the number of measurements for each subject, $\hat{\Delta}$ is a diagonal matrix with elements $\hat{\Delta}_{ii} = (Y_i - X_i\hat{\beta})^2/(1-H_{ii})^2$ and $H_{ii}$ is the $i$-th diagonal element of the hat matrix [10].

**Theorem 1.** *Let all objects be as defined above and assume that for all* $a, b = 1, \ldots, p_1$, $\text{Cov}\left(\hat{\epsilon}_{ij}^2, \hat{\epsilon}_{ik}^2\right)(x_{ija}x_{ijb})(x_{ika}x_{ikb})$ *and* $n_i$ *are bounded for all* $i,\ j, k = 1, \ldots, n_i$. *Then the estimator*

$$\hat{S} = \left(\max\left\{0, \frac{T_{cs}^2 - p_1}{N}\right\}\right)^{\frac{1}{2}}, \tag{8}$$

*is consistent for* (5), *where* $T_{cs}^2 = N(\hat{\beta} - \beta_0)^T \hat{\Sigma}_{\beta_{cs}}^{-1}(\hat{\beta} - \beta_0)$, *and* $\hat{\beta}$ *is the estimator for* $\beta$ *from a GEE.*

*Proof.* Since the marginal parameters are estimated in GEEs using identify link function, the parameter estimates are consistent for the same parameter in the cross-sectional and longitudinal models. Therefore, the estimator (8) is a consistent estimator for the CS-RESI (5) as long as

$$\hat{\Sigma}_{\beta_{cs}} \xrightarrow{p} \Sigma_{\beta_{cs}},$$

where

$$\Sigma_{\beta_{cs}} = \lim_{N\to\infty} (N^{-1}X^TWX)^{-1}(N^{-1}X^TW\Delta WX)(N^{-1}X^TWX)^{-1} := A_{cs}^{-1}B_{cs}A_{cs}^{-1}.$$

It is sufficient to show that $\hat{A}_{cs} \xrightarrow{p} A_{cs}$ and $\hat{B}_{cs} \xrightarrow{p} B_{cs}$ then $\hat{\Sigma}_{\beta_{cs}} \xrightarrow{p} \Sigma_{\beta_{cs}}$ will follow by Slutsky's theorem.

The element on the $a$-th row and $b$-th column of $B_{cs}$ is

$$\begin{aligned} B_{cs,(a,b)} &= \lim_{N\to\infty} \frac{1}{N}\mathbb{E}_R\left(\sum_{i=1}^{N}\sum_{j=1}^{n_i} R_{ij}x_{ija}x_{ijb}\sigma_{ij}^2\right) \\ &= \lim_{N\to\infty} \frac{1}{N}\sum_{i=1}^{N}\frac{1}{n_i}\sum_{j=1}^{n_i} x_{ija}x_{ijb}\sigma_{ij}^2. \end{aligned}$$

$\hat{B}_{cs,(a,b)}$ converges in probability if it is asymptotically unbiased and $\mathrm{Var}(\hat{B}_{a,b}) \to 0$ as $N \to \infty$ for $a$, $b = 1, \ldots, p_1$. The corresponding element of $\hat{B}_{cs}$ is $\frac{1}{N}\sum_{i=1}^{N}\frac{1}{n_i}\sum_{j=1}^{n_i} x_{ija}x_{ijb}\hat{\epsilon}_{ij}^2$, where $x_{ija}$ is the value of the $a$-th covariates (including the intercept) of the $j$-th observation from the $i$-th subject. The asymptotic unbiasedness of $\hat{B}_{cs,(a,b)}$ follows because $\hat{\epsilon}_{ij}^2 = \frac{(Y_{ij}-X_{ij}\hat{\beta})^2}{1-h_{ij,ij}}$ is unbiased estimator for $\sigma_{ij}^2$. The variance of $\hat{B}_{cs,(a,b)}$ is

$$\begin{aligned} \mathrm{Var}(\hat{B}_{cs,(a,b)}) &= \mathrm{Var}\left(\frac{1}{N}\sum_{i=1}^{N}\sum_{j=1}^{n_i}\frac{1}{n_i}x_{ija}x_{ijb}\hat{\epsilon}_{ij}^2\right) \\ &= \frac{1}{N^2}\sum_{i=1}^{N}\frac{1}{n_i^2}\sum_{j=1}^{n_i}\sum_{k=1}^{n_i}(x_{ija}x_{ijb})(x_{ika}x_{ikb})\mathrm{Cov}\left(\hat{\epsilon}_{ij}^2, \hat{\epsilon}_{ik}^2\right) \\ &= \frac{1}{N^2} \times \sum_{i=1}^{N}\frac{1}{n_i^2}\left[\sum_{j=1}^{n_i}(x_{ija}x_{ijb})^2\mathrm{Var}(\hat{\epsilon}_{ij}^2) + 2\sum_{j\le k}\mathrm{Cov}\left(\hat{\epsilon}_{ij}^2, \hat{\epsilon}_{ik}^2\right)(x_{ija}x_{ijb})(x_{ika}x_{ikb})\right] \\ &= \frac{1}{N^2}\mathcal{O}\left(\sum_{i=1}^{N}\frac{1}{n_i}\right) + \frac{2}{N^2}\mathcal{O}\left(\sum_{i=1}^{N}\frac{1}{n_i}\right) \\ &= \mathcal{O}\left(\frac{1}{N^2}\sum_{i=1}^{N}\frac{1}{n_i}\right) \end{aligned}$$

thus, $\mathrm{Var}(\hat{B}_{cs,(a,b)}) \to 0$ as $N \to \infty$.

Similarly, the element on the $a$-th row and $b$-th column of $\hat{A}_{cs}$ and $A_{cs}$ are

$$\hat{A}_{cs,(a,b)} = \frac{1}{N}\sum_{i=1}^{N}\frac{1}{n_i}\sum_{j=1}^{n_i} x_{ija}x_{ijb}$$

and

$$\begin{aligned} A_{cs,(a,b)} &= \frac{1}{N}\sum_{i=1}^{N}\sum_{j=1}^{n_i}\mathbb{E}_R(R_{ij})x_{ija}x_{ijb} \\ &= \frac{1}{N}\sum_{i=1}^{N}\frac{1}{n_i}\sum_{j=1}^{n_i} x_{ija}x_{ijb} \end{aligned}$$

Thus, for given $\boldsymbol{X}$, $Var(\hat{A}_{cs,(a,b)}) = 0$ and $\mathbb{E}(\hat{A}_{cs,(a,b)}) - A_{cs,(a,b)} \to 0$, thus we have $\hat{A}_{cs} \xrightarrow{p} A_{cs}$. Then $\hat{\Sigma}_{\beta_{cs}} \xrightarrow{p} \Sigma_{cs}$ by Slutsky's theorem, which completes the proof.

□

The CS-RESI estimate assumes that the parameters estimated in the cross-sectional and longitudinal models are equivalent. This is violated when the actual between- and within-subject effects are not equal and the longitudinal analysis model (either linear mixed model (LMM) or GEE) assumes a non-independence working covariance structure (see Section 3.3).

##### 1.1.3 Confidence Interval Construction

As shown in our previous work, the distribution of the test statistic may deviate from the theoretical chi-square or $F$ distributions when an estimator is used for the covariance and/or the design matrix is random [5]. In these situations, the confidence intervals (CIs) constructed based on the theoretical distributions will fail to provide nominal coverage. Bootstrapping can be used to approximate the sampling distribution of the RESI estimators. We consider the standard nonparametric bootstrap by sampling the subjects with replacement $r$ times and estimating the corresponding L-RESI and CS-RESI for each resampled data. The lower and upper $\alpha/2 \times 100$ percentiles of the $r$ estimated RESIs are the bootstrapped lower and upper bounds of the $(1-\alpha) \times 100$ bootstrap CI for the estimated L-RESI or CS-RESI.

#### 1.2 Simulation Studies

In the previous sections, we define the L-RESI and CS-RESI, propose corresponding estimators, and the non-parametric bootstrap for CI construction. Here, we use simulations to evaluate the influence of fixed/random covariates, and model misspecification on the performance of the estimators and CIs.

##### 1.2.1 Simulation Setup

We simulate longitudinal data for each subject using a linear model $Y_i = \beta_0 + \beta_g \times \text{group} + \beta_t \times \text{time} + \epsilon_i$, which contains a time-independent binary variable "group" and a time-dependent variable "time" across 5 visits, for 1,000 independent simulations. To simulate the within-subject correlation, we specify $\text{Cov}(\epsilon_i)$ to have either exchangeable or first-order autoregressive (AR-1) correlation structure. When using the exchangeable correlation structure,

$$\text{Cov}(\epsilon_i) = \begin{bmatrix} 1 & 0.5 & \cdots & 0.5 \\ 0.5 & 1 & \cdots & 0.5 \\ \vdots & & \ddots & \vdots \\ 0.5 & 0.5 & \dots & 1 \end{bmatrix},$$

and when using the AR-1 structure,

$$\text{Cov}(\epsilon_i) = \begin{bmatrix} 1 & 0.5 & 0.5^2 & 0.5^3 & 0.5^4 \\ 0.5 & 1 & & \cdots & 0.5^3 \\ \vdots & & & \ddots & \vdots \\ 0.5^4 & 0.5^3 & 0.5^2 & \dots & 1 \end{bmatrix}.$$

We vary the sample size $N \in \{50, 250, 500\}$ across three different true L-RESI values $S \in \{0, 0.5, 1\}$. The true L-RESIs of the group variable and the time variable are specified to be the same in each scenario. The true CS-RESI values are calculated accordingly. To show the influence of experimental and observational designs where the covariates are fixed or random, the values of the covariates are generated in two ways: (1) when being fixed, $\pi = 0.5$ and the random ceiling($N\pi$) of the $N$ individual subjects have their group variable with value 1 and the remaining have value 0, and everyone has 5 different observations at timepoints $= \{0, 1, \cdots, 4\}$; (2) when being random, group $\sim$ Bernoulli($\pi$) is sampled from a Bernoulli distribution with parameter $\pi = 0.5$ for each subject, and everyone has 5 observations at 5 timepoints randomly chosen in the range of 0 to 4. The working correlation structure is specified to be "exchangeable" or "AR1", regardless of the true correlation structure to evaluate the influence of model misspecification on the performance of the estimators and CIs. For each bootstrap CI, 1,000 bootstraps are used.

To evaluate the consistency of the CS-RESI between the longitudinal analysis and its equivalent cross-sectional analysis, we use block sampling to create the equivalent artificial cross-sectional sample by randomly selecting one observation from each subject in each of the simulated longitudinal datasets. In each of the simulations, the CS-RESI is estimated using the GEE model in the longitudinal data and the regular RESI designed for cross-sectional studies is estimated using a linear model with the same model formula as the GEE in the equivalent cross-sectional data. The mean difference between the CS-RESI from the longitudinal data and the regular RESI from the cross-sectional data (defined as the CS-RESI in the longitudinal data minus the regular RESI in the cross-sectional data) is estimated across the simulations. This mean difference should approach zero at large sample sizes if CS-RESI has the desired comparability.

##### 1.2.2 Simulation Results

Simulations are performed to assess the bias of the estimators and the coverage of the CIs. We consider different cases where there are random/fixed covariates and/or misspecification of the model correlation structure.

In small samples ($N = 50$), the estimators for the CS-RESI and the longitudinal RESI are both slightly biased for both the group and time variables, but the bias goes to zero as sample size increases (Figure SI1 and SI2). The estimators for the CS-RESI and L-RESI show consistency in all scenarios, including when the correlation structure is mistakenly specified (Figure SI1B and C; Figure SI2B and C). The variable randomness doesn't show an influence on the estimator biasedness.

The bootstrap CI provides nominal coverage for the CS-RESI and L-RESI under all scenarios when the models are correctly specified (Figure SI3A and D; SI4A and D). When the models are misspecified (Figure SI3B and C; SI4B and C), the CI sometimes provides coverage lower than the nominal level, but the coverages approach the nominal level as sample size increases. The variable randomness doesn't show an influence on the coverage of bootstrap CI.

As to the consistency/comparability of the CS-RESI in the longitudinal study to the regular RESI in the equivalent cross-sectional study, Figure SI5 shows the mean difference between them across all simulations. The difference is defined as CS-RESI minus regular RESI. In small samples ($N = 50$), the CS-RESI in longitudinal data slightly overestimates the effect of the group variable and underestimates the effect of the time variable in cross-sectional studies when $S = 0$, but the mean difference approaches zero in large sample sizes. When $S \neq 0$ and $N = 50$, the CS-RESI tends to overestimate the effect of both the group and time variables, but the mean difference goes to zero in large sample sizes. The results indicate that the CS-RESI in longitudinal studies essentially estimates the same thing as if the studies are conducted cross-sectionally and the consistency is not affected by model misspecification and variable randomness.

#### 1.3 Comparison of L-RESI and CS-RESI quantifies benefit of longitudinal design

While the systematic difference in the standardized ES between cross-sectional and longitudinal study designs compromises the comparability of standardized ESs between studies, it is also

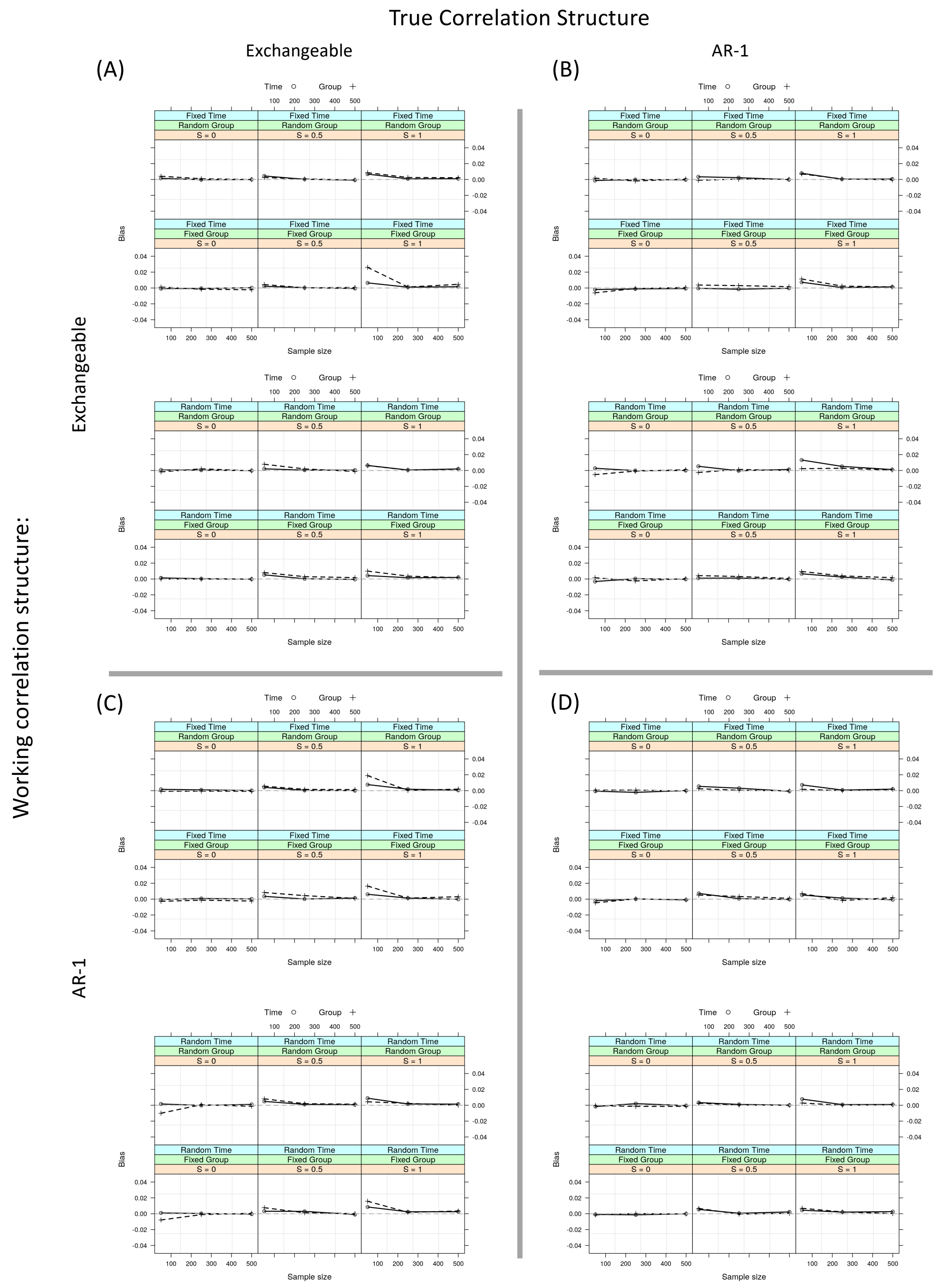


Figure SI1: The bias of the CS-RESI estimator for both group and time variables under the simulation settings described above. The bias of the CS-RESI estimator goes to zero as sample size increases in all scenarios, indicating it is an asymptotic unbiased estimator. $S$ denotes the L-RESI, which is specified the same for both the group and time variables; the true values of CS-RESI are calculated accordingly in each scenario.

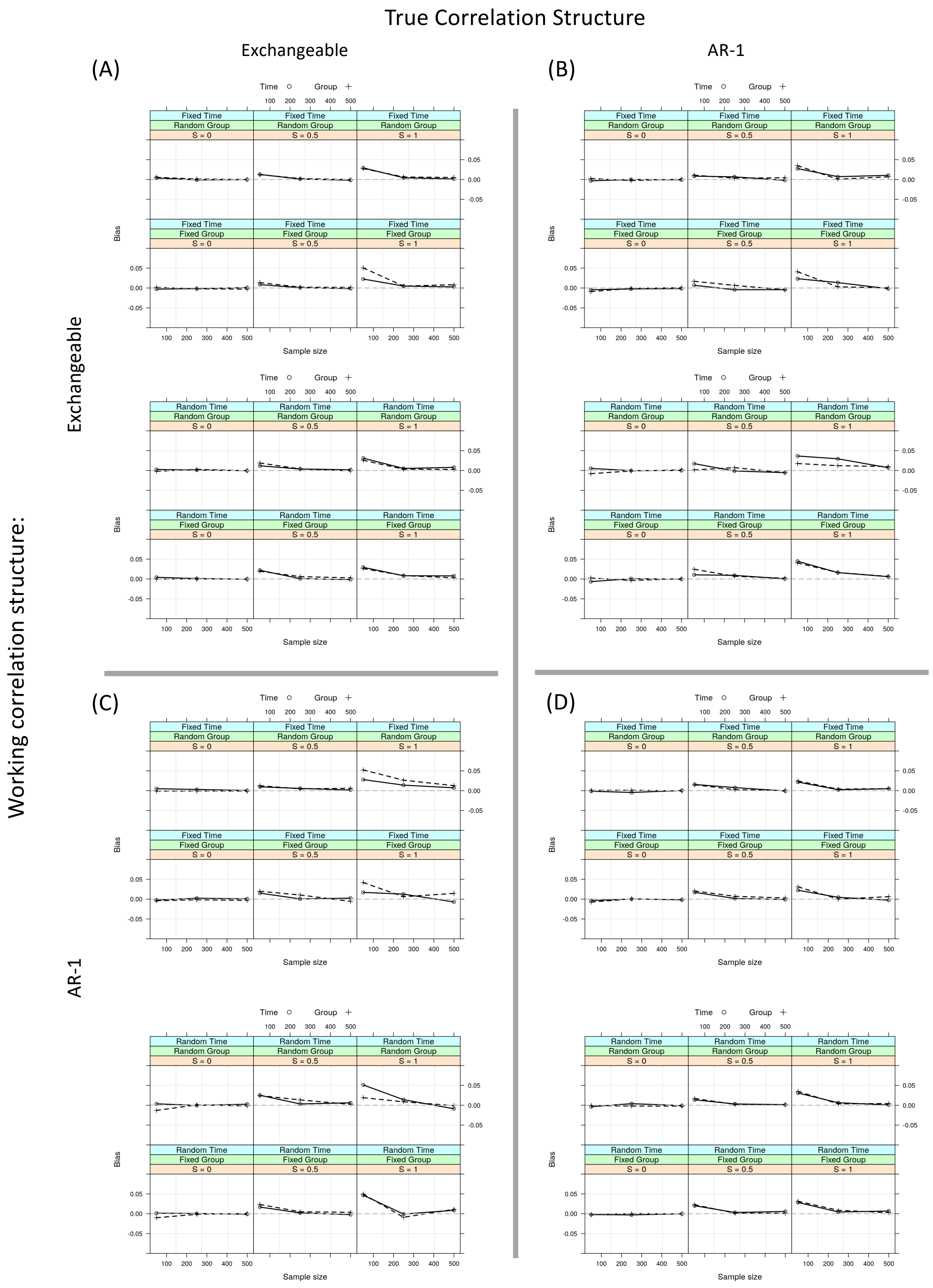


Figure SI2: The bias of the L-RESI estimator for both group and time variables under the simulation settings described above. The bias of the L-RESI estimator goes to zero as sample size increases in all scenarios, indicating it's an asymptotic unbiased estimator. $S$ denotes the longitudinal RESI, which is specified the same for both the group and time variables.

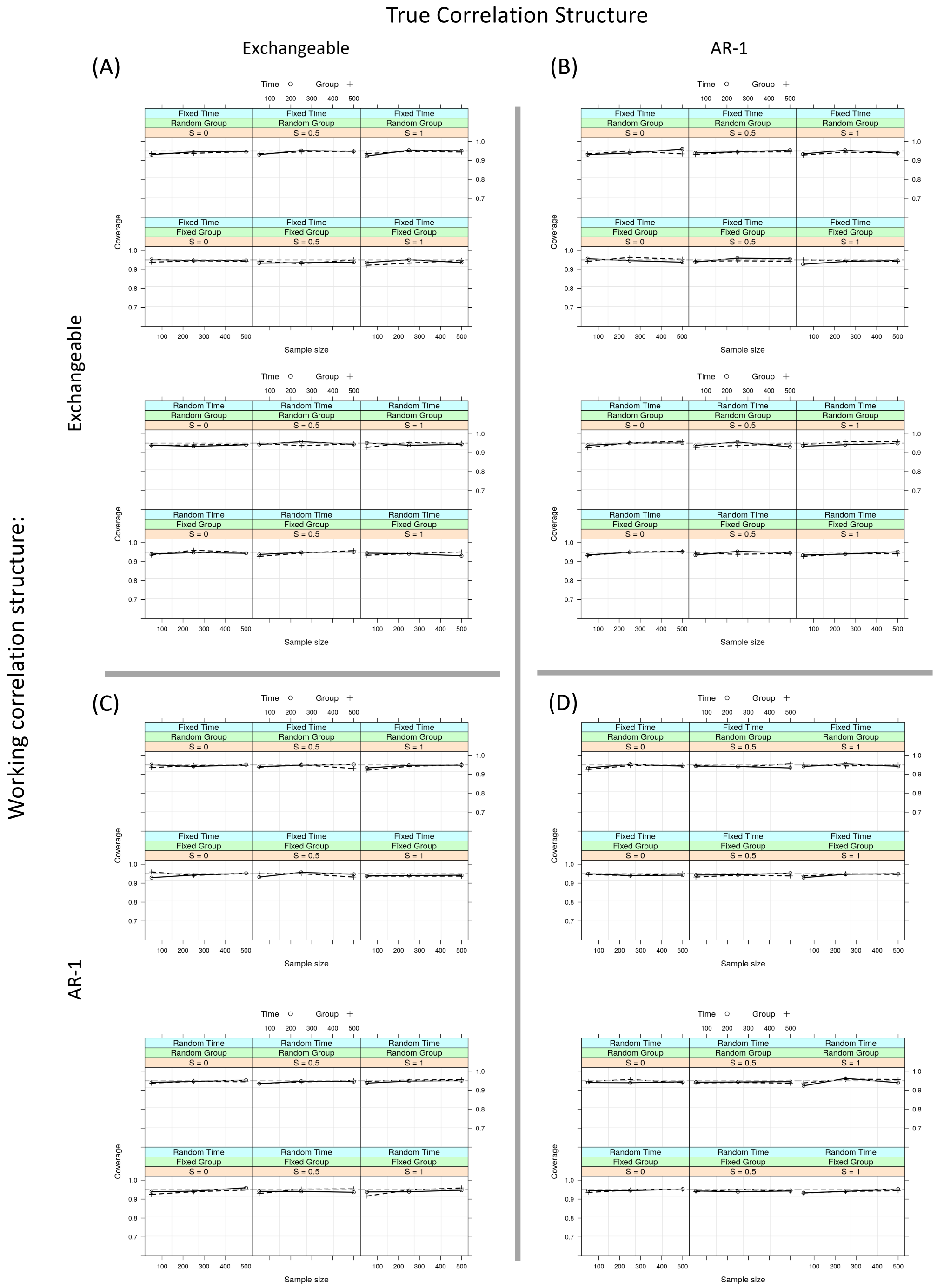


Figure SI3: The coverage of bootstrap CI for the CS-RESI estimator under the simulation settings described above. The bootstrap CI asymptotically provides nominal coverage for the CS-RESI under all scenarios even when the models are misspecified. $S$ denotes the L-RESI, which is specified the same for both the group and time variables; the true values of CS-RESI are calculated accordingly in each scenario.

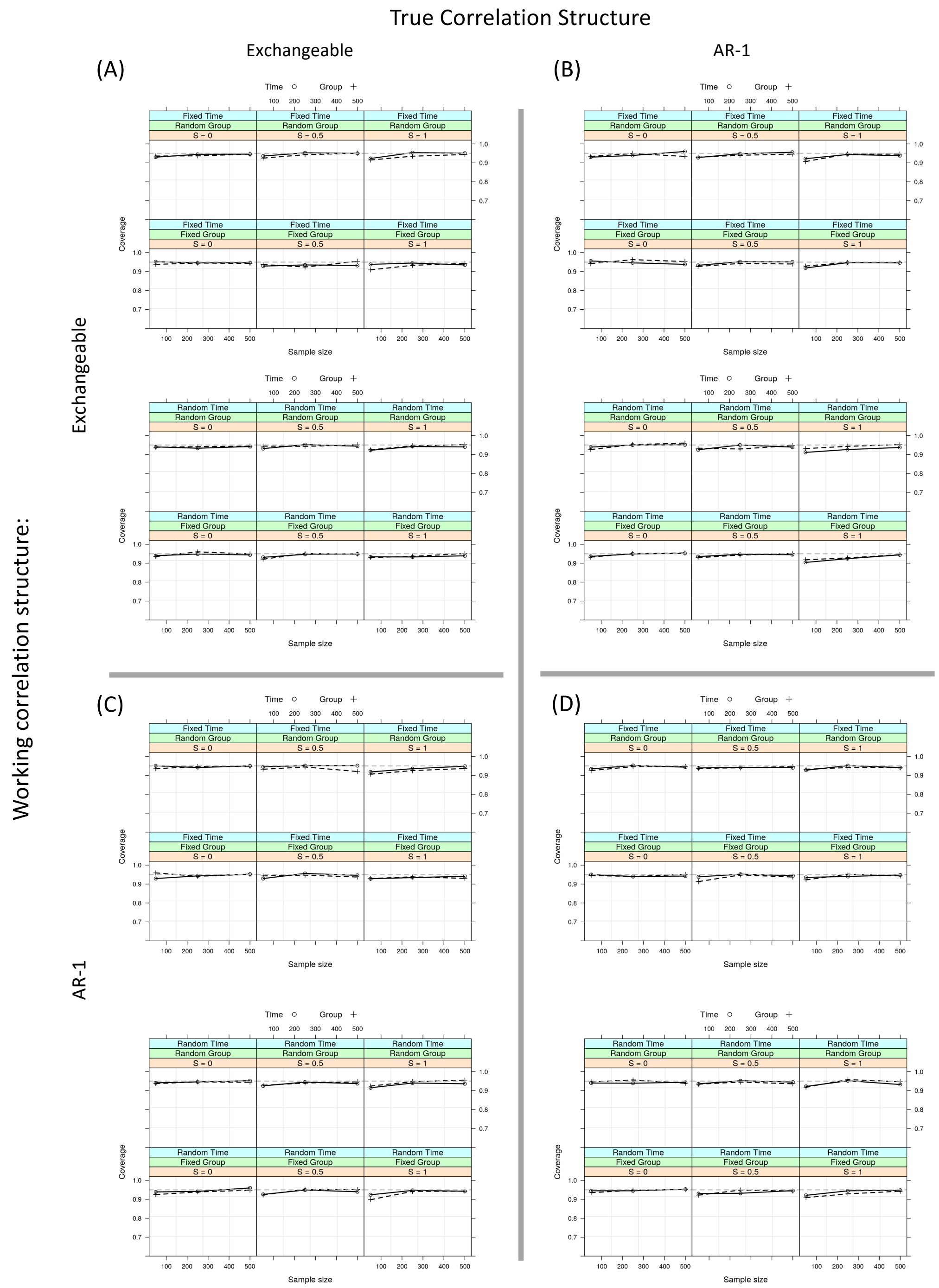


Figure SI4: The coverage of bootstrap CI for the L-RESI estimator under the simulation settings described above. The bootstrap CI asymptotically provides nominal coverage for the L-RESI and under all scenarios even when the models are misspecified. $S$ denotes the L-RESI, which is specified the same for both the group and time variables.

a feature that we can leverage to quantify the benefit of using a longitudinal design in a single study. With this goal in mind, we compare the difference between the L-RESI and CS-RESI in each of the longitudinal studies from the LBCC analyzed in the main paper to quantify the benefit of using the longitudinal study design.

To demonstrate the difference in standardized ES by study design and validate the CS-RESI estimator, we use non-parametric bootstrapping to create longitudinal bootstrap samples by randomly sampling subjects with replacement (1,000 bootstraps per sample size) in each of the 16 longitudinal neuroimaging datasets we use in the main paper. In this analysis and subsequent bootstrap analyses, the full dataset represents the unknown population, and bootstrap replicates represent random samples from the population. At a given sample size, after obtaining a longitudinal bootstrap sample (including all measurements from the selected subjects), we create an equivalent cross-sectional version of the longitudinal bootstrap sample by randomly selecting one measurement from each selected subject. In each replicate, we fit the same GEEs and linear regression models of total GMV we use in the main paper to the longitudinal and cross-sectional bootstrap samples, and quantify the effects of age and sex on total GMV. L-RESI and CS-RESI are computed in the longitudinal bootstrap samples and compared to the RESI computed in the cross-sectional bootstrap samples. The differences of the RESI estimated in the cross-sectional sample to the CS-RESI and L-RESI estimated in the longitudinal sample are calculated in each replicate, and the mean differences are derived across the replicates. All site effects are removed using LongComBat before performing the bootstrap analysis.

When comparing the L-RESI estimated in the longitudinal samples to the RESI estimated in cross-sectional samples, the longitudinal standardized ESs for age and sex are typically larger (Figure SI6a). The positive differences represent the amount of benefit of using longitudinal designs to quantify the age or sex effect for the two studies shown (ADNI and GUSTO). Age shows an increase of 0.16 in GUSTO and 0.25 in ADNI, while sex shows an expectedly smaller increase of 0.02 in GUSTO and ADNI (other studies shown in Figure SI7). The difference we note in longitudinal standardized ESs is not restricted to the L-RESI, as Cohen's $d$ for sex computed in a longitudinal study also shows the same slightly positive estimate (Figure SI6a). In contrast, the mean difference between the CS-RESI computed in the longitudinal samples and the RESI computed in the cross-sectional samples approaches zero as the sample size increases (Figure SI6b), indicating that the systematic difference in the standardized ESs between longitudinal and cross-sectional datasets is removed. As a further comparison, we repeat the meta-analyses in the main paper on the standardized ESs from all studies but now use the CS-RESIs to represent the standardized ESs in longitudinal studies, demonstrating a reduction in the design effect (Tab. SI1, SI2, SI3, and SI4).

| | Estimate | S.E. | $F$ | d.f. | $p$-value | Meta-RESI | 95% CI |
|---|---|---|---|---|---|---|---|
| Design: Longitudinal | 0.02 | 0.08 | 0.06 | 1 | 0.81 | 0.00 | (0.00, 0.26) |
| Mean age | | | 3.91 | 3 | 0.01 | 0.36 | (0.18, 1.02) |
| SD of age | | | 20.23 | 3 | <0.01 | 0.94 | (0.58, 1.72) |
| Skewness of age | | | 0.77 | 3 | 0.52 | 0.00 | (0.00, 0.61) |

*S.E.: robust (sandwich) standard error; F: F statistic; d.f.: the first degree(s) of freedom for the F statistics; RESI: robust effect size index; L-RESI: longitudinal RESI; CS-RESI: cross-sectional RESI; CI: confidence interval; SD: standard deviation

Table SI1: Meta-analysis results for the RESI for age on total gray matter volume (GMV). The residual degrees of freedom are 52. The regression model is weighted by the inverse standard errors of the RESIs. For cross-sectional studies, the outcome is RESI; for longitudinal studies, the outcome is CS-RESI.

The benefit of conducting the longitudinal design can be quantified in each study by comparing the CS-RESI estimates with the L-RESI estimates (Figure SI6c; Figure SI8 and Figure SI9). Longitudinal designs result in a 0.31 average increase in the standardized ES for the total GMV-

| | Estimate | S.E. | $F$ | d.f. | $p$-value | Meta-RESI | 95% CI |
|---|---|---|---|---|---|---|---|
| Design: Longitudinal | -0.03 | 0.07 | 0.20 | 1 | 0.65 | 0.00 | (0.00, 0.3) |
| Mean age | | | 1.10 | 3 | 0.36 | 0.05 | (0.00, 0.77) |
| SD of age | | | 7.09 | 3 | <0.01 | 0.53 | (0.06, 1.11) |
| Skewness of age | | | 0.17 | 3 | 0.91 | 0.00 | (0.00, 0.47) |

*S.E.: robust (sandwich) standard error; F: F statistic; d.f.: the first degree(s) of freedom for the F statistics; RESI: robust effect size index; L-RESI: longitudinal RESI; CS-RESI: cross-sectional RESI; CI: confidence interval; SD: standard deviation

Table SI2: Meta-analysis results for the RESI for age on total subcortical gray matter volume (sGMV). The residual degrees of freedom are 52. The regression model is weighted by the inverse standard errors of the RESIs. For cross-sectional studies, the outcome is RESI; for longitudinal studies, the outcome is CS-RESI.

| | Estimate | S.E. | $F$ | d.f. | $p$-value | Meta-RESI | 95% CI |
|---|---|---|---|---|---|---|---|
| Design: Longitudinal | 0.00 | 0.06 | 0.00 | 1 | 0.95 | 0.00 | (0.00, 0.28) |
| Mean age | | | 4.88 | 3 | <0.01 | 0.42 | (0.21, 1.23) |
| SD of age | | | 3.36 | 3 | 0.03 | 0.33 | (0.00, 0.92) |
| Skewness of age | | | 0.46 | 3 | 0.71 | 0.00 | (0.00, 0.60) |

Table SI3: Meta-analysis results for the RESI for age on total white matter volume (WMV). The residual degrees of freedom are 52. The regression model is weighted by the inverse standard errors of the RESIs. For cross-sectional studies, the outcome is RESI; for longitudinal studies, the outcome is CS-RESI.

| | Estimate | S.E. | $F$ | d.f. | $p$-value | Meta-RESI | 95% CI |
|---|---|---|---|---|---|---|---|
| Design: Longitudinal | 0.05 | 0.17 | 0.08 | 1 | 0.79 | 0.00 | (0.00, 0.50) |
| Mean age | | | 1.84 | 3 | 0.16 | 0.22 | (0.00, 1.27) |
| SD of age | | | 4.11 | 3 | 0.01 | 0.45 | (0.00, 1.44) |
| Skewness of age | | | 0.05 | 3 | 0.99 | 0.00 | (0.00, 0.69) |

Table SI4: Meta-analysis results for the RESI for age on mean cortical thickness (CT). The residual degrees of freedom are 32. The regression model is weighted by the inverse standard errors of the RESIs. For cross-sectional studies, the outcome is RESI; for longitudinal studies, the outcome is CS-RESI.

age association among the 16 longitudinal neuroimaging datasets (Tab. S3A in the main paper), corresponding to a moderate increase in standardized ES [4]. Notably, in some studies (e.g., POND and SCZIowa), the L-RESI is smaller than the CS-RESI, which implies that they lose efficiency by using longitudinal designs. We investigate this counterintuitive result further in our subsequent analyses of cognitive variables in the ABCD study in the main paper.

### 2 Modified Sampling Design Considerations

#### 2.1 Sampling estimates for targeted sampling

As we show in the main paper, we can use modified sampling designs to obtain a sample with a particular histogram shape to increase covariate variance, standardized ESs and replicability. The decision to pursue this design can be determined by the choices in Fig. S13. An estimate of the additional number of subjects needed to reach a particular sample size using the modified design can be computed if the population distribution of the covariate and modified sampling distribution are known. We assume that we can decide whether to scan each participant, whose covariate values are collected first. To compute the expected number of participants needed to collect, we assume that the data are binned on a consistent grid such as that used to make a histogram of the empirical data and that, to obtain a sample from the target modified distribution $g$, we will randomly sample from the true distribution $f$. In this case, both $f$ and $g$ can be treated as multinomial distributions. Assume that each distribution has $K$ bins on the same grid, then the probability the total sample size, $N$ is larger than $m$, when trying to obtain a sample of size $n$ from the modified distribution is

$$\mathbb{P}(N > m) = \mathbb{P}(N_k < n\pi_{gk} \text{ for one } k = 1, \dots, K) \tag{9}$$

$$= 1 - \mathbb{P}(N_k \geq n\pi_{gk} \text{ for all } k = 1, \dots, K) \tag{10}$$

where $(N_1, \dots, N_K) \sim \text{multinom}(\pi_{f1}, \dots, \pi_{fK}, m)$ are multinomial. Summing this over all possible values of $m$ gives the expected total sample size, $N$, required to obtain $n$ samples that exactly have the target distribution using the integrated tail probability formula (see e.g., [13]). Code to evaluate this probability is available in Section 4.

| $n$ | N |
|---|---|
| 50 | 298 |
| 100 | 516 |
| 250 | 1155 |
| 500 | 2201 |

Table SI5: Required total sample size, $N$, when true distribution of covariate is a normal distribution in order to obtain a sample size of $n$ from a uniform distribution for the covariate.

Besides optimizing study efficiency, there are non-statistical factors that should be considered when choosing a study design, such as the practical challenges in recruiting the study population. Because of the complexity of resources at different study sites, it is hard to provide specific details of how to execute the modified sampling without information about the covariate and how the imaging data are collected. For example, if the covariate varies slowly over time, the measurement can be ascertained in an initial screening and then a subset can be selected for imaging on a later date based on their measured values. If the covariate varies with cognitive state, then it might be necessary to administer cognitive testing and immediately scan a subset or to schedule the imaging and re-administer the cognitive test at scanning. Such complex design issues should be determined on a case-by-case basis.

#### 2.2 Implications for multivariate BWAS

It is important to distinguish between univariate and multivariate BWAS. In univariate BWAS, investigators treat the brain measure as the outcome and model these measurements with a non-brain covariate. In contrast, multivariate BWAS treat the non-brain measurement as the outcome and use machine learning methods to predict the outcome variable as a function of the imaging variable. Multivariate BWAS tend to have larger standardized ESs than univariate BWAS because the signal in neuroimaging data is likely to be dispersed across networks of brain regions. Our analyses focus on univariate BWAS analyses with tens to thousands of brain measurements; our findings directly generalize to higher-dimensional mass-univariate analyses because the standardized ESs do not depend on the number of comparisons although can be biased after selecting regions using multiplicity adjustment. Our findings may generalize to the multivariate BWAS setting, where the imaging variable is used as the covariate, based on previous research that optimizes power using outcome-dependent sampling and two-phase study designs, but further research is merited.

### 3 Longitudinal Design and Analysis Considerations

Two factors affect standardized ESs such as equation (3): the value of the target parameter (determined through the term involving $\beta$) and the efficiency of the estimator of the target parameter (determined by $\Sigma_\beta$). In the main paper, we discuss how between- and within-subject effects may differ across brain non-brain association pairs. When this is true, but the longitudinal model falsely assumes they are equal, then the expected value of the estimated parameter is a weighted average of the two. To understand how the expected value of the parameter estimate is affected, we assume the true model

$$\begin{aligned} Y_{ij} &= \beta_0 + \beta_b \bar{x}_i + \beta_w (x_{ij} - \bar{x}_i) + \epsilon_{ij} \\ \mathrm{Var}(\epsilon_{ij}) &= \Sigma_\epsilon, \end{aligned} \tag{11}$$

where we treat $\Sigma_\epsilon$ as a known exchangeable covariance matrix. $i = 1, \ldots, n$ indicates the participant and $j = 1, \ldots, m$ indicates the number of measurements, which is assumed to be the same across all participants. This model can be fit using LMMs and GEEs. $\beta_b$ represents the between-subject effects and $\beta_w$ represents the within-subject effects. See [14] for the distinction of random effects and between- and within-subject effects.

Unless explicitly parameterized, as in (11), the fitted model for longitudinal data in BWAS often assumes a mean model with $\beta_b = \beta_w$,

$$Y_{ij} = \beta_0 + \beta x_{ij} + \epsilon_{ij}. \tag{12}$$

This is the most common type of model specification used in BWAS. We hypothesized in the main paper that the detrimental effects of increasing within-subject variance or using a longitudinal study were due to the use of model (12) when model (11) was correct. When incorrectly specified with model (12), the expected value of the estimator for $\beta$ is a combination of the underlying true between- and within-subject effects given by

$$\begin{aligned} \mathbb{E}\hat{\beta} = \beta_b &\left[ \frac{(1-\rho)\rho_x^2}{(1+(m-1)\rho)(1-\rho_x^2) + (1-\rho)\rho_x^2} \right] \\ + \beta_w &\left[ \frac{(1+(m-1)\rho)(1-\rho_x^2)}{(1+(m-1)\rho)(1-\rho_x^2) + (1-\rho)\rho_x^2} \right] \end{aligned} \tag{13}$$

where $\rho_x^2 = \frac{\sigma_b^2}{\sigma^2}$, which represents the within-subject correlation strength of the covariate $x_{ij}$, the $\sigma_b^2$ and $\sigma^2$ denote the between-subject and total variance of the covariate, and $\rho$ is the correlation of the errors or outcome $\epsilon_{ij}$. Derivation of this value and subsequent values are given in Section 3.4. Assuming the variance matrix $\Sigma_\epsilon$ is known, the variance of $\sqrt{n}\hat{\beta}$ is

$$\mathrm{Var}(\sqrt{n}\hat{\beta}) = \left[ m \frac{(1+(m-1)\rho)\sigma_w^2 + (1-\rho)\sigma_b^2}{\sigma_\epsilon^2 (1+(m-1)\rho)(1-\rho)} \right]^{-1},$$

where $\sigma_w^2$ is the within-subject variance of the covariate $x_{ij}$, $\sigma_\epsilon^2$ is the error variance, and all other objects are as defined above. The signed RESI ES (4) is a function of the expected value of $\hat{\beta}$ and its variance.

$$\begin{aligned} S_{\mathrm{sgn}} = \frac{\mathbb{E}\hat{\beta}}{\sqrt{\mathrm{Var}(\sqrt{n}\hat{\beta})}} &= m^{1/2}\{(1+(m-1)\rho)\sigma_w^2 + (1-\rho)\sigma_b^2\}^{-1/2}\sigma_y^{-1/2} \\ &\times \left\{ \beta_b \sigma_b^2 \left[ \frac{(1-\rho)}{1+(m-1)\rho} \right]^{1/2} + \beta_w \sigma_w^2 \left[ \frac{(1+(m-1)\rho)}{(1-\rho)} \right]^{1/2} \right\}. \end{aligned} \tag{14}$$

Equation (13) implies that, when using a single parameter to model the between- and within-subject effects, modifications of the longitudinal study design can change the target parameter, as well as the estimation efficiency, so can affect the numerator and denominator of the ES in equation (14). The formula is visualized in Figure SI10. The expected value of the parameter estimator is a convex combination of the between- and within-subject parameters. The expected value of the ES estimator is a linear combination of $\beta_b$ and $\beta_w$, where the weights are complicated functions of the within-subject correlation of the outcome, the number of measurements, and the between- and within-subject variance of the covariate. As shown in the main paper (Fig. 6e-f), the between- and within-subject effects $\beta_b$ and $\beta_w$ can differ. In the paper, because our sampling schemes control the variance at the first timepoint and the relative squared distance from the first timepoint, the parameters we estimate are those given in equation (16).

When fitting a cross-sectional version of model (12) to a randomly selected observation from each subject, as in (6), the expected value of the parameter is

$$\mathbb{E}(\hat{\beta}_{CS}) = \beta_b \rho_x^2 + \beta_w (1 - \rho_x^2).$$

This is derived in Section 3.4, but is also obtained by setting $\rho = 0$ and $m = 1$ in (13). These results imply that the longitudinal model (12) is estimating a different parameter than an equivalent cross-sectional model because it incorporates the correlation of the outcome into the weighting. Changing features of a longitudinal design change the target parameter of (12), which will also affect the standardized ES. In most cases, if the between- and within-subject associations are different, they should be estimated with separate parameters. We propose three different strategies that can be used to model the association between the brain measure and non-brain covariate in BWAS. The first assumes that the between- and within-subject effects are equal and the second and third assume they are different.

#### 3.1 Equal between- and within-subject effects

When $\beta_b = \beta_w = \beta$ then there is no model misspecification and increasing the between- or within-subject variance of the covariate increases ESs for the parameter $\beta$. In this case, the parameters $\beta_b$ and $\beta_w$ can be distributed out of the sum in (14) and the whole ES increases linearly with respect $\sigma_w$ and $\sigma_b$.

When designing a study to estimate $\beta$, it is useful to know which study design is more efficient per subject and per measurement (such as per MRI scan). To do this we compare the ratio of the variances of the estimator obtained from a cross-sectional design $\hat{\beta}_{CS}$, to that obtained in a longitudinal design $\hat{\beta}_L$, while assuming the total number of scans in each study is the same, i.e. $n = Nm$.

$$\begin{aligned} \frac{\mathrm{var}(\hat{\beta}_{\mathrm{CS}})}{\mathrm{var}(\hat{\beta}_{\mathrm{L}})} &= \left[ (Nm) \frac{(1 + (m-1)\rho)\sigma_w^2 + (1-\rho)\sigma_b^2}{\sigma_\epsilon^2 (1 + (m-1)\rho)(1-\rho)} \right] \times \frac{\sigma_\epsilon^2}{n\sigma^2} \\ &= \frac{(1 + (m-1)\rho)(1 - \rho_x^2) + (1-\rho)\rho_x^2}{(1 + (m-1)\rho)(1-\rho)}. \end{aligned} \tag{15}$$

Setting this ratio greater than one and solving for $\rho$, we have

$$\rho > \frac{(m\rho_x^2 - 1)}{(m-1)},$$

which indicates that if the within-subject correlation of the outcome conditional on the covariate is large (i.e., if the within-subject error of brain measure is low) relative to $m$ times the within-subject correlation of the covariate, then the longitudinal design will be more efficient.

Here we provide some example code of how to estimate the within-subject correlation of the outcome (i.e., the parameter $\rho$ above) and the within-subject correlation of the covariate (i.e., the parameter $\rho_x^2$ above) in GEEs using the R package `geeglm`. We regress the age and sex effects on total GMV in ADNI data for illustration, and we only restrict to the first and last visits of the ADNI participants (i.e., to set $m = 2$). In the following code, the object `ADNI` is a data frame containing all the variables needed, with the continuous variable age (in years) and binary variable sex under the variable names `age_years` and `sex_01`, respectively. The outcome variable `GMV_10000_combat` is the total GMV after removing the site effect using longComBat[15] and scaled by 10,000.

```
# Fit the GEE model with an exchangeable correlation structure
library(geepack)
fit <- geeglm(GMV_10000_combat ~ sex_01 + age_years,
              data = ADNI, corstr = "exchangeable",
              id = participant)
summary(fit)
```

From the output produced by `summary(fit)` below, we can find the estimate for the parameter within-subject correlation of the outcome to be 0.902 by looking at the estimate of `alpha` under `Estimated Correlation Parameters`.

```
Call:
geeglm(formula = GMV_10000_combat ~ sex_01 + age_years, data = ADNI,
    id = participant, corstr = "exchangeable")

 Coefficients:
            Estimate Std.err Wald Pr(>|W|)
(Intercept)  58.6207  1.1957 2404   <2e-16 ***
sex_01        3.7981  0.3030  157   <2e-16 ***
age_years    -0.2574  0.0156  272   <2e-16 ***
---
Signif. codes:  0 ‘***’ 0.001 ‘**’ 0.01 ‘*’ 0.05 ‘.’ 0.1 ‘ ’ 1

Correlation structure = exchangeable
Estimated Scale Parameters:

            Estimate Std.err
(Intercept)     9.93   0.705
  Link = identity

Estimated Correlation Parameters:
      Estimate Std.err
alpha    0.902  0.0101
Number of clusters:   417  Maximum cluster size: 2
```

The within-subject correlation of covariate age can be estimated in R as below:

```
library(magrittr, dplyr)
# Calculate the within-subject correlation of covariate age
## 1. calcualte the between-subject variance of age
sigma2_b = ADNI %>% group_by(participant) %>%
    summarise(subj_mean_age = mean(age_years)) %>%
    ungroup() %>% summarise(sigma2_b = var(subj_mean_age)) %>%
    as.numeric
```

```
## 2. calculate the total variance of age
sigma2 = var(ADNI$age_years)
## the within-subject correlation of age
rho2_x = sigma2_b / sigma2
rho2_x
[1] 0.878
```

The estimated within-subject correlations of the outcome and the covariate are both pretty high, with $\hat{\rho} = 0.902$ and $\hat{\rho}_x^2 = 0.878$. Plugging them into the equation (15) along with $m = 2$ shows that the estimated ratio of efficiency is 1.71, indicating that, given a fixed total number of scans $n$, scanning $n/2$ participants twice and doing a longitudinal analysis is 71% more efficient than doing a cross-sectional analysis with scanning each of $n$ participants once when studying the total GMV-age association in situations of this study.

Differentiating (15) with respect to $m$ shows how the efficiency changes with respect to changes in the number of scans per subject, keeping the number of total scans fixed

$$\frac{d}{dm}\frac{\text{var}(\hat{\beta}_{\text{CS}})}{\text{var}(\hat{\beta}_{\text{L}})} = \frac{-\rho\rho_x^2}{(1+(m-1)\rho)^2}.$$

This function is non-positive for all values of $m > 1$, indicating a reduction in efficiency relative to the cross-sectional design with increasing number of measurements per participant, if the within-subject correlation of the outcome and the within-subject correlation of covariate are both fixed with increasing $m$. These formulas explain the results in Fig. 4d, where adding one additional measurement is beneficial, but more than one additional measurement has a diminishing benefit.

In this setting and if the data are cross-sectional, a model such as

$$Y_i = \beta_0 + \beta x_i + \epsilon_i,$$

is appropriate. If the data are longitudinal, a linear mixed model (LMM) or generalized estimating equation (GEE) as in (12) is appropriate.

#### 3.2 Modeling different between- and within-subject effects

When the between- and within-subject effects are different it is probably best to model them separately because it avoids averaging the larger effect with the smaller effect, and because the distinction between these two effects can inform our understanding of brain-behavior associations. In this case, increasing between- or within-subject variance independently increases the ES for the between- or within-subject parameter when (11) is the correct model. To fit this model, we include the original covariate $x_{ij}$ which varies over measurements from the same individual, and also include a covariate that is the mean of the original covariate $\bar{x}_i = n_i^{-1}\sum_{j=1}^{n_i} x_{ij}$, as in equation (11). The mean covariate will be the same value for all measurements within participants. An LMM or GEE can be used to model and estimate the parameters separately. The parameters can be interpreted separately as trait versus state associations with the outcome.

Various parameterizations are possible. They look similar but differ in their interpretation and mathematical formulation. For example, in the main paper, we use an alternative to (11) that models a separate effect for the baseline measurements, which influences all future measurements.

$$Y_{ij} = \beta_0^* + \beta_b^* x_{i1} + \beta_w^*(x_{ij} - x_{i1}) + \epsilon_{ij}. \tag{16}$$

Here, the between- and within-subject variance are defined as

$$\begin{aligned}\sigma_b^2 &= \frac{1}{N}\sum_{i=1}^{N}(x_{i1} - \bar{x}_1)^2 \\ \sigma_w^2 &= \frac{1}{N}\sum_{i=1}^{N}\frac{1}{n_i}\sum_{j=1}^{n_i}(x_{ij} - x_{i1})^2\end{aligned} \tag{17}$$

This model makes sense if the first measurement in the study represents a particular epoch that is comparable across participants. In the paper, we used this formulation, because our sampling schemes are based on the first measurement and the change from first measurement, which separately affect the between- and within-subject variance shown in (17).

Here, as an example, we provide some example code for estimating the between- and within-subject effects of CBCL on the outcome total GMV (after controlling for age and sex) separately in the ABCD data, as the model specified in (16). First, we need to create variables indicating the subject-specific baseline covariate measures and the change of the covariate from the baselines.

```
# create a variable `x_bl` to indicate
# the subject-specific baseline CBCL values (i.e., x_i1)
ABCD %<>% group_by(temp_id) %>%
    mutate(x_bl = CBCL[eventname == "Baseline"]) %>% ungroup()
# create a variable `x_change` to indicate (x_ij - x_i1)
ABCD %<>% mutate(x_change = CBCL - x_bl)
```

The variables `x_bl` and `x_change` in the data frame object `ABCD` should look like the following:

```
   temp_id eventname        CBCL        x_bl x_change  ...
 1       1 Baseline         36            36  0
 2       2 Baseline         32            32  0
 3       3 Baseline         51            51  0
 4       4 Baseline         49            49  0
 5       5 Baseline         58            58  0
 6       5 2-year follow-up 61            58  3
 7       5 4-year follow-up 55            58 -3
 8       6 Baseline         51            51  0
 9       6 2-year follow-up 55            51  4
10       6 4-year follow-up 51            51  0
...
```

Then we can estimate the between- and within-subject effects of CBCL in a GEE:

```
# fit a GEE model
library(geeglm)
fit = geeglm(GMV_10000_combat ~ age_years + sex_01 + x_bl + x_change,
              id = temp_id, corstr = "exchangeable", data = ABCD)
summary(fit)

Call:
geeglm(formula = GMV_10000_combat ~ age_years + sex_01 + x_bl +
    x_change, data = ABCD, id = temp_id, corstr = "exchangeable")

 Coefficients:
```

```
             Estimate  Std.err     Wald Pr(>|W|)
(Intercept) 77.82355  0.25265 94884.60   <2e-16 ***
age_years   -0.61448  0.01004  3742.46   <2e-16 ***
sex_01       6.04305  0.10527  3295.50   <2e-16 ***
x_bl        -0.04259  0.00481    78.24   <2e-16 ***
x_change    -0.00766  0.00280     7.51   0.0061 **
---
Signif. codes:  0 ‘***’ 0.001 ‘**’ 0.01 ‘*’ 0.05 ‘.’ 0.1 ‘ ’ 1

Correlation structure = exchangeable
Estimated Scale Parameters:
            Estimate Std.err
(Intercept)     33.2    0.47
  Link = identity

Estimated Correlation Parameters:
      Estimate Std.err
alpha    0.915  0.0114
Number of clusters:   11718  Maximum cluster size: 3
```

In the output, the estimates for the between-subject effect and the within-subject effect (i.e., $\beta_b^*$ and $\beta_w^*$ in (16)) are the coefficient estimates for the terms `x_bl` and `x_change`.

###### 3.2.1 Interpretation of between- and within-subject effects

The interpretation of the between- and within-subject parameters will be dependent on the context and the parameterization. The parameters can be better understood by looking at the expected differences between two measurements within (18) and between (19) participants,

$$\mathbb{E}(Y_{ij} - Y_{ik}) = \beta_w(x_{ij} - x_{ik}) \tag{18}$$
$$\mathbb{E}(Y_{ij} - Y_{\ell k}) = \beta_b(\bar{x}_i - \bar{x}_k) + \beta_w \left\{(x_{ij} - \bar{x}_i) - (x_{\ell k} - \bar{x}_\ell)\right\}. \tag{19}$$

Comparing measurements for two values of the covariate within a subject is expected to have a different effect than comparing the same two covariate values between subjects. From (18), we can see that if we plot $(Y_{ij} - Y_{ik})$ versus $(x_{ij} - x_{ik})$, the slope captures only the within-subject effect, which can be used for visually evaluating the within-subject effect (see e.g., Fig. S12 in the main paper). This can occur if the covariate has high measurement variability, but the outcome variable does not, then we would expect a within-subject change in the covariate to be unrelated to within-subject changes in the outcome (i.e., $\beta_w = 0$). For psychometric measurements that can be affected by the subject's state at measurements, $\bar{x}_i$ in (11) represents subject-specific mean "trait" value of the covariate and the term $(x_{ij} - \bar{x}_i)$ is the within-subject deviation of measurement $j$ from this mean trait value. If the brain outcome measure (e.g., GMV) is expected not to vary by state, then it might be reasonable to believe that $\beta_w \approx 0$, whereas with FC data, it's likely that within-subject association varies across regions as the measurements may be similarly affected by the subject's stat at measurements.

For certain covariates, it does not make sense to consider different between- and within-subject effects, depending on the study design. For example, consider the parameterization in (16) for the GMV-age association in an observational study, it would be odd if the age that a participant entered the study had an additional effect on GMV above the effect of their age when the GMV measurement was taken. In this parameterization, the absence of this first timepoint effect implies $\beta_b^* = 0$, or in the parameterization (11), $\beta_b = \beta_w$. Alternatively, if the study entry represented the occurrence of a particular event, e.g., diagnosis, then it seems plausible that $\beta_b^* \neq 0$.

If there are potential nonlinearities then each effect may be estimated with multiple degrees of freedom, e.g., by using splines. The interpretation of parameter estimates from model (11) will not be comparable to the estimate from a cross-sectional model because the cross-sectional model cannot distinguish the two effects and is estimating the combination. If the goal is to obtain an estimate that is comparable to the one obtained in a cross-sectional study, then see Section 3.3.

#### 3.3 Modeling the averaged parameter

If the true model has different between- and within-subject parameters, despite that it is incorrect, model (12) can still be used. This might be of interest when it is hard to interpret the within-subject effect. In this case, the weighted average from the cross-sectional model depends on whether (11) or (16) is the true model. We recommend targeting the parameter obtained from a cross-sectional model with one measurement per participant (the weighted average of the between- and within-subject parameters),

$$Y_{i1} = \beta_0 + \beta x_{i1} + \epsilon_{i1}. \tag{20}$$

so that longitudinal studies target the same parameter(s) as cross-sectional studies. If (16) is the true model, then that is a special case where the expected value of the cross-sectional model parameter estimator (using model (20)) is equal to the between-subject parameter in Equation (16). When model (20) is fit, but (11) is true, we can use formula (13) to assess the bias of the estimators under different models (Table SI6). As can be seen from Table SI6, only the GEE with independence working covariance structure targets and unbiasedly estimates the same parameter as the cross-sectional model, but it is still less efficient than a cross-sectional model. The commonly used LMM or GEE with another covariance structure is biased for this parameter, where the bias depends on the structure of the working covariance matrix. In the case of LMM, the working covariance matrix is assumed to be the true covariance. Thus, in this setting, only GEE with independence working covariance can be used to estimate $\beta$ in (20) in a longitudinal model. In general, when using other correlation structures in GEEs, to increase ES without biasing the parameter estimator, the between- and within-subject variance must be increased proportionally. This is because the weights for the between- and within-subject effects in $\mathbb{E}\hat{\beta}$ are determined by $\rho_x^2 = \frac{\sigma_b^2}{\sigma_b^2+\sigma_w^2}$ (as defined in Section 3.4); increasing $\sigma_b^2$ or $\sigma_w^2$ independently will change the value of the parameter. Thus, a sampling scheme that increases both proportionally must be adopted.

The longitudinal model is less efficient per scan than the cross-sectional model under mild assumptions that are likely to hold in BWAS. To see this, we compare the ratio of the variances of the two estimators. Assuming that in each study design we collect $Nm$ total brain measurements, then the ratio of the variance of the estimator in the longitudinal GEE with independence working covariance to the one of the cross-sectional model (20), is

$$\begin{aligned}\frac{\mathrm{var}(\hat{\beta}_{\mathrm{GEE}})}{\mathrm{var}(\hat{\beta}_{\mathrm{CS}})} &= \frac{\sum_{i=1}^{N}\left(\sum_{j=1}^{m}\sigma_\epsilon^2 X_{ij}^2 + \sum_{j\neq k}^{m}\rho\sigma_\epsilon^2 X_{ij}X_{ik}\right)}{\left(\sum_{i,j}X_{ij}^2\right)^2} \times \frac{\sum_{i}^{Nm}X_{i1}^2}{\sigma_\epsilon^2} \\ &\geq \frac{\sum_{i,j}X_{ij}^2 + \sum_{i=1}^{N}\sum_{j\neq k}^{m}\rho X_{ij}X_{ik}}{\sum_{i,j}X_{ij}^2} \geq 1,\end{aligned}$$

where the first inequality assumes that the behavioral measurements between subjects are on average larger than measurements within subjects, $\sum_i^{Nm} X_{i1}^2 \geq \sum_{i,j} X_{ij}^2$ and the second inequality assumes that brain and behavioral repeated measurements are each positively correlated within-subject. Practically, this analysis indicates that when targeting an averaged parameter (i.e., row 1 of Table SI6), it is better to perform a cross-sectional study rather than a longitudinal study. Note, however, that with a fixed number of subjects, collecting more longitudinal measurements will still generally increase efficiency relative to a cross-sectional study. Thus, longitudinal data should be used in the GEE framework if it is available.

| Model | Parameters | $\mathbb{E}\hat{\beta}$ |
|---|---|---|
| Cross-sectional study | $\rho = 0, m = 1$ | $\mathbb{E}\hat{\beta} = \beta_b \rho_x + \beta_w(1 - \rho_x)$ |
| GEE independence work covariance | $\rho = 0, m > 1$ | $\mathbb{E}\hat{\beta} = \beta_b \rho_x + \beta_w(1 - \rho_x)$ |
| LMM/GEE exchangeable covariance | $\rho \neq 0, m > 1$ | Equation (13) |

Table SI6: Expected values of the parameter estimates from (20), or (12) under different working covariances. If the goal is to estimate the same parameter as the cross-sectional model (20), then LMM/GEE with non-independence covariance structure are biased, because the weights for $\beta_b$ and $\beta_w$ are as given in (13). The only longitudinal model that is unbiased for the cross-sectional effect is the GEE with independence working covariance (second row). $\rho_x^2 = \frac{\sigma_b^2}{\sigma^2}$ as defined in Section 3.4.

#### 3.4 Derivation of ES under exchangeable design

After residualizing $x$ and $Y$ to the intercept design matrix, the expected value of $\hat{\beta}$ is

$$\mathbb{E}(\hat{\beta} \mid \mathbf{x}) = (\mathbf{x}^T \Sigma^{-1} \mathbf{x})^{-1} \mathbf{x}^T \Sigma^{-1} Y = (\mathbf{x}^T \Sigma^{-1} \mathbf{x})^{-1} \mathbf{x}^T \Sigma^{-1} [\bar{\mathbf{x}}, \mathbf{w}] \begin{bmatrix} \beta_b \\ \beta_w \end{bmatrix},$$

where $Y, \mathbf{x}, \bar{\mathbf{x}}, \mathbf{w} \in \mathbb{R}^{nm}$ with

$$\begin{aligned} \mathbf{x}_{(i-1)\times m+j} &= x_{ij} - \bar{x} \\ \bar{\mathbf{x}}_{(i-1)\times m+j} &= \bar{x}_i - \bar{x} \\ \mathbf{w}_{(i-1)\times m+j} &= \mathbf{x}_{(i-1)\times m+j} - \bar{\mathbf{x}}_{(i-1)\times m+j} \\ \bar{x} &= \frac{1}{nm} \sum_{i,j} x_{ij} \\ \bar{x}_i &= \frac{1}{m} \sum_{j=1} x_{ij} \end{aligned}$$

When $\Sigma_\epsilon$ is exchangeable,

$$\Sigma_\epsilon^{-1} = aI + b(I_n \otimes \mathbf{1}\mathbf{1}^T),$$

where $a = (\sigma_\epsilon^2(1-\rho))^{-1}$ and $b = a \left[ \frac{-\rho}{(1+(m-1)\rho)} \right]$, $\mathbf{1}$ denotes an $m$ dimensional vector of all ones, $I_n$ is the $n$ dimensional identity matrix, and $\otimes$ denotes the Kronecker product. The inverse variance of $\sqrt{n}\hat{\beta}$ is

$$\begin{aligned} \mathrm{Var}(\sqrt{n}\hat{\beta})^{-1} &= n^{-1}(\mathbf{x}^T \Sigma_\epsilon^{-1} \mathbf{x}) \\ &= \frac{m}{\sigma_\epsilon^2(1-\rho)} \left[ \sigma^2 - \sigma_b^2 \frac{m\rho}{(1+(m-1)\rho)} \right] \\ &= m \left[ \frac{(1+(m-1)\rho)\sigma_w^2 + (1-\rho)\sigma_b^2}{\sigma_\epsilon^2 (1+(m-1)\rho)(1-\rho)} \right] \\ &= m \frac{\sigma^2}{\sigma_\epsilon^2} \left[ \frac{(1+(m-1)\rho)(1-\rho_x^2) + (1-\rho)\rho_x^2}{(1+(m-1)\rho)(1-\rho)} \right] \end{aligned} \tag{21}$$

where

$$\sigma^2 = \frac{1}{nm}\sum_{i,j}(x_{ij} - \bar{x})^2$$
$$\sigma_b^2 = \frac{1}{n}\sum_{i=1}^{n}(\bar{x}_i - \bar{x})^2$$
$$\sigma_w^2 = \frac{1}{nm}\sum_{i,j}(x_{ij} - \bar{x}_i)^2$$
$$\rho_x^2 = \frac{\sigma_b^2}{\sigma^2} = \frac{\sigma_b^2}{\sigma_w^2 + \sigma_b^2}.$$

Then $\mathbb{E}\hat{\beta}$ simplifies to

$$\begin{aligned}
\mathbb{E}\hat{\beta} &= \beta_w + (\beta_b - \beta_w) \times \frac{a\mathbf{x}^T\bar{\mathbf{x}} + b\mathbf{x}^T(I_n \otimes \mathbf{1}\mathbf{1}^T)\bar{\mathbf{x}}}{a\mathbf{x}^T\mathbf{x} + b\mathbf{x}^T(I_n \otimes \mathbf{1}\mathbf{1}^T)\mathbf{x}} \\
&= \beta_w + (\beta_b - \beta_w)m\sigma_b^2(a + bm)\mathrm{Var}(\sqrt{n}\hat{\beta}) \\
&= \beta_w + (\beta_b - \beta_w)m\frac{\sigma_b^2}{\sigma_\epsilon^2(1 + (m-1)\rho)}\mathrm{Var}(\sqrt{n}\hat{\beta}) \\
&= \beta_w + (\beta_b - \beta_w)\left[\frac{(1-\rho)\sigma_b^2}{(1 + (m-1)\rho)\sigma_w^2 + (1-\rho)\sigma_b^2}\right] \\
&= \beta_b\left[\frac{(1-\rho)\sigma_b^2}{(1 + (m-1)\rho)\sigma_w^2 + (1-\rho)\sigma_b^2}\right] \\
&\quad + \beta_w\left[\frac{(1 + (m-1)\rho)\sigma_w^2}{(1 + (m-1)\rho)\sigma_w^2 + (1-\rho)\sigma_b^2}\right] \\
&= \beta_b\left[\frac{(1-\rho)\rho_x^2}{(1 + (m-1)\rho)(1-\rho_x^2) + (1-\rho)\rho_x^2}\right] \\
&\quad + \beta_w\left[\frac{(1 + (m-1)\rho)(1-\rho_x^2)}{(1 + (m-1)\rho)(1-\rho_x^2) + (1-\rho)\rho_x^2}\right]
\end{aligned}$$

Combining (21) and (13) gives (14).

### 4 Sample size code for modified sampling

```
library(pmultinom)
totalN = function(n, nbin=10, alpha = 0.05, eps=0.001,
                 g='unif', f='norm'){
  # target number in each bin
  if(tolower(g)=='unif'){
    gbin = rep(1/nbin, nbin) * n
  }
  # true distribution of the data
  fp = get(paste0('p', f))
  fq = get(paste0('q', f))
  fdomain = c(fq(alpha/2), fq(alpha/2, lower.tail=FALSE))
  fbin = c(-Inf, seq(fdomain[1], fdomain[2], length.out=nbin-1), Inf)
  fbin = diff(fp(fbin))
```

```
  # finds max sample size to get the desired precision
  nmax = 500
  tailProb = 1
  while(tailProb>eps){
      tailProb = 1-pmultinom(lower=gbin, size=nmax,
        probs = fbin, method='exact')
      nmax = nmax + 200
    }
# computes multinomial probabilities
  probs = sapply(0:nmax, function(N){pmultinom(lower=gbin, size=N,
    probs = fbin, method='exact')})
  # Using expectation of CDF to compute this expectation
  sum(1-probs)
}
```

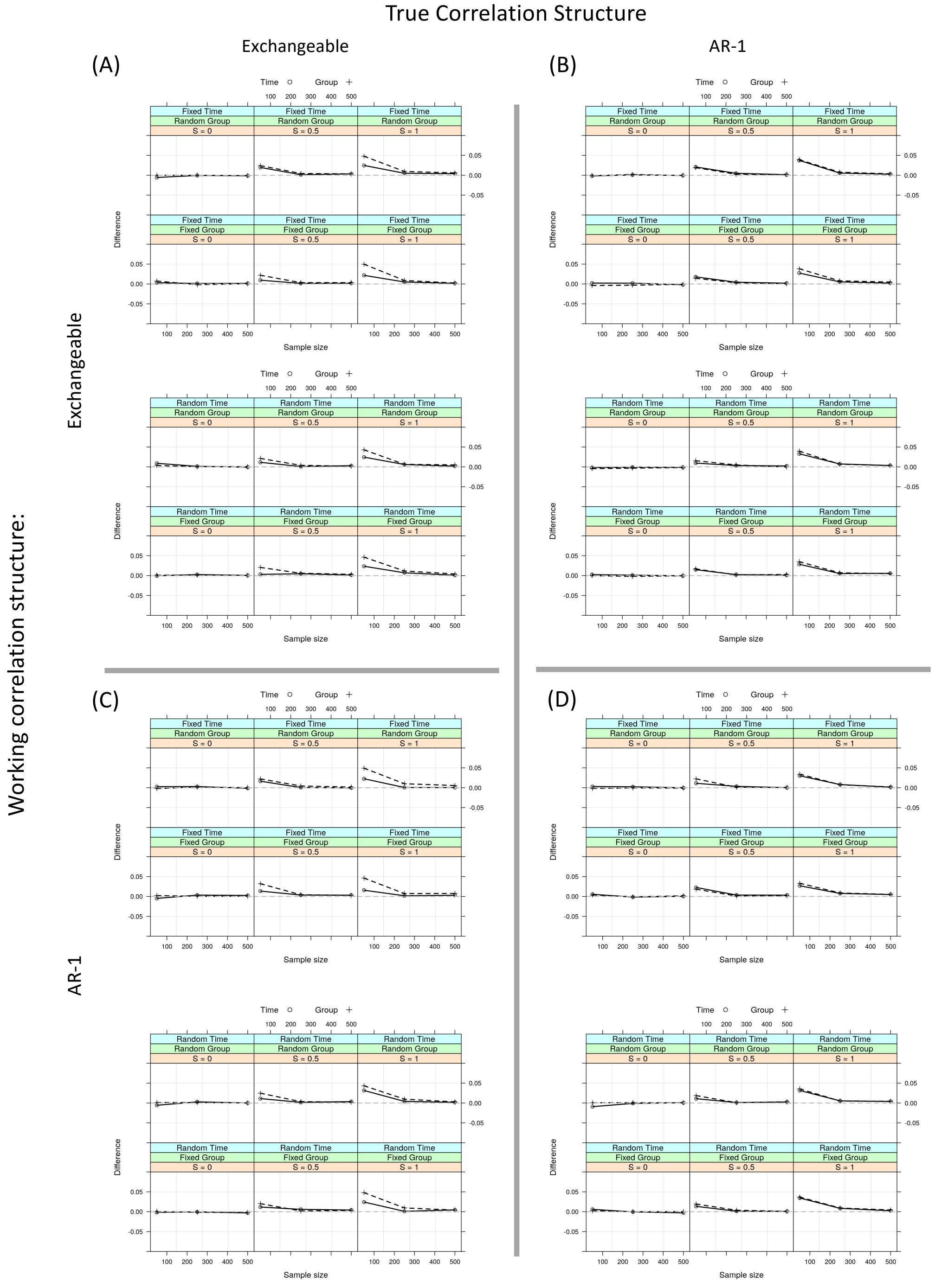


Figure SI5: The mean difference between the CS-RESI in longitudinal studies and the regular RESI in the equivalent cross-sectional studies. The mean difference approaches zero in all scenarios including when the models are misspecified, indicating the consistency/comparability between the CS-RESI in longitudinal studies and the regular RESI in cross-sectional studies. $S$ denotes the longitudinal RESI, which is specified the same for both the group and time variables.

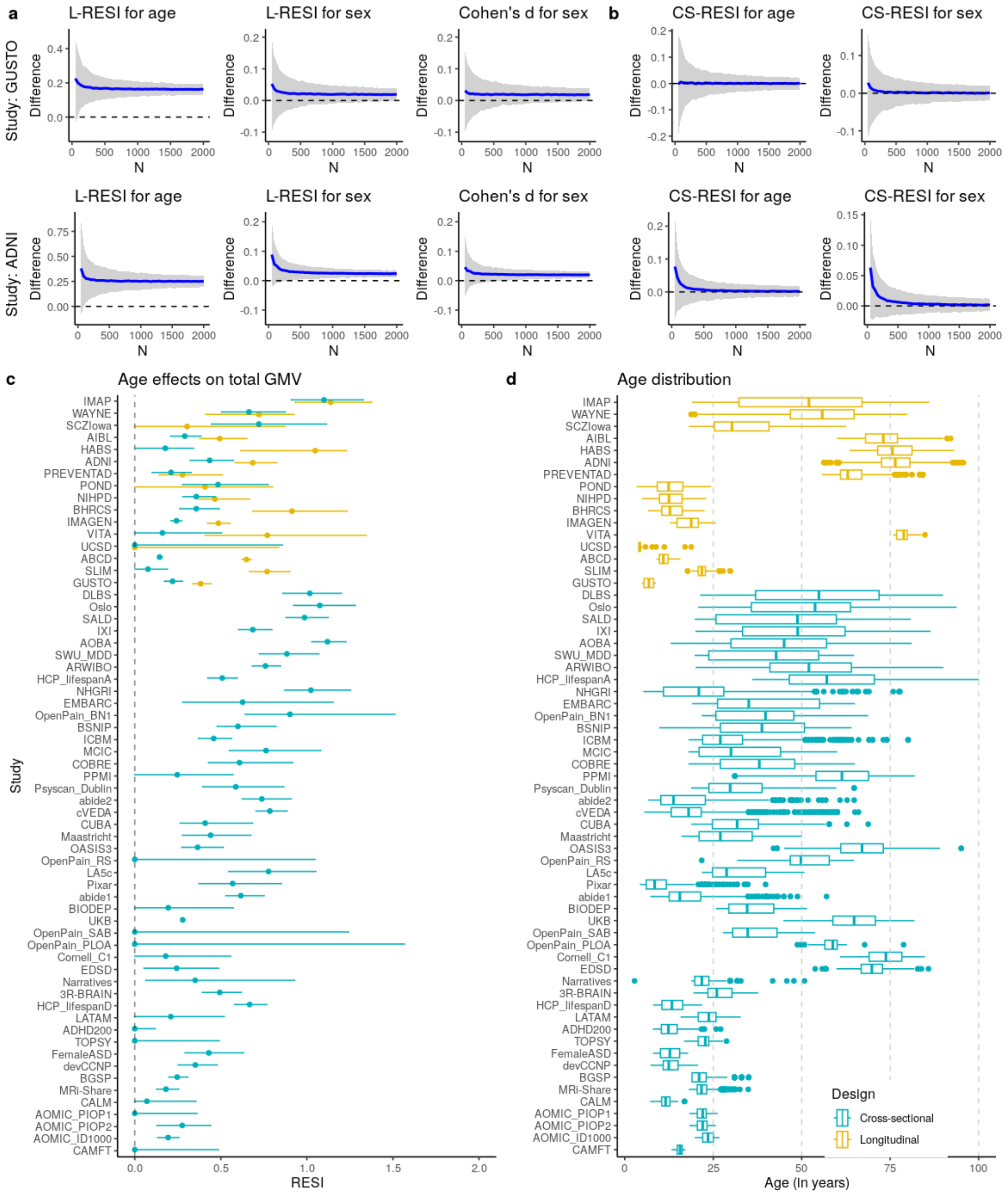


Figure SI6: (a) Bootstrap analysis shows that standardized effect sizes (ESs) of age and sex on total gray matter volume (GMV) in longitudinal studies are systematically larger than the standardized ESseffect sizes in cross-sectional studies (Difference = L-RESI - RESI) and (b) the CS-RESI for longitudinal analysis estimates and removes the difference (Difference = CS-RESI - RESI). We conduct 1,000 bootstraps per sample size (indicated by N on the x-axes). The blue curves represent the mean difference curves and the gray areas are the 95% confidence bands from the bootstraps. (c) Forest plot of the standardized ESs for total GMV-age association shows that the CS-RESIs in longitudinal studies are more comparable to the regular RESIs in cross-sectional studies. Regular RESIs (blue) are estimated for cross-sectional studies, CS-RESIs (blue) and L-RESIs (yellow) are both estimated for longitudinal studies. The dots and lines represent RESI estimates and their 95% confidence intervals (CIs). The studies are sorted by study design and decreasing standard deviation (SD) of age in each study. (d) The age distribution in each study; studies with smaller SD of age tend to have smaller age effects.

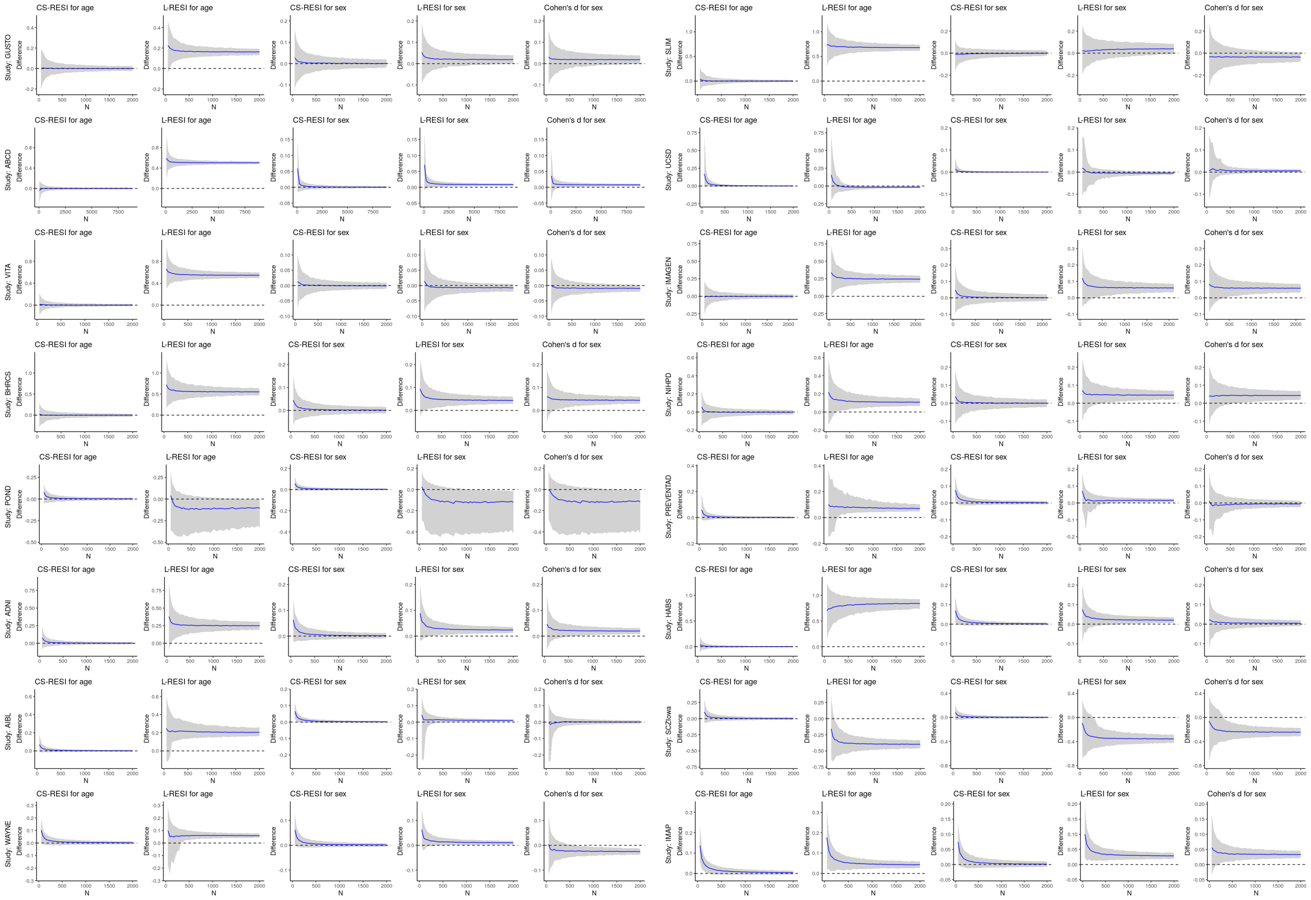


Figure SI7: The systematic difference in standardized effect size (ES) by study designs and the comparability of CS-RESI (cross-sectional RESI) in longitudinal studies to RESI in cross-sectional studies. We conduct 1,000 bootstraps per sample size N. The blue curves represent the mean difference curves and the gray areas are the 95% confidence bands from the bootstraps. For most of the studies, the differences of L-RESI (longitudinal RESI) in longitudinal datasets to regular RESI in cross-sectional datasets approach non-zero constants as sample size increases. However, the differences of CS-RESI in longitudinal datasets to the regular RESI in cross-sectional datasets always approach zero at large sample sizes, indicating the systematic difference in standardized ESseffect sizes between longitudinal and cross-sectional datasets is removed by using CS-RESI.

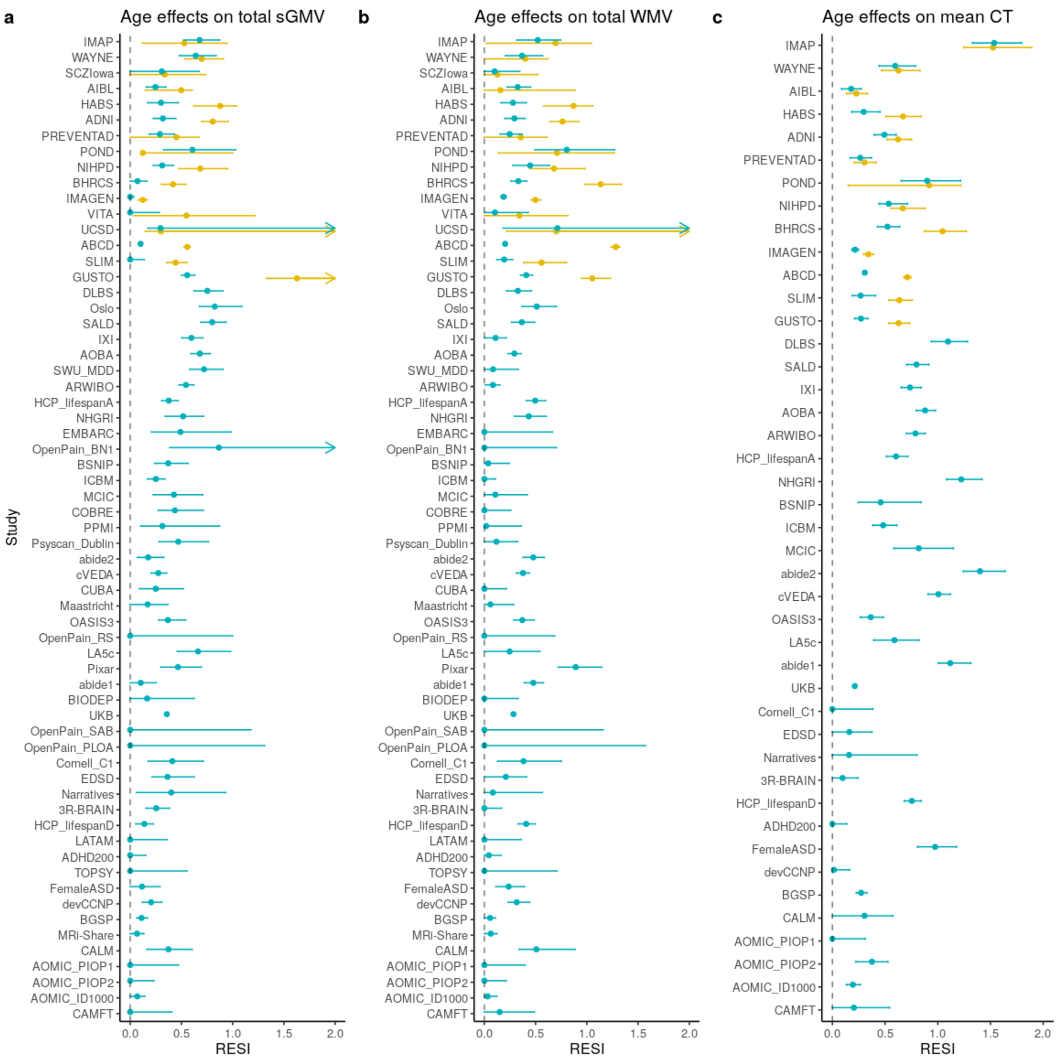


Figure SI8: Forest plots of age effects on total subcortical gray matter volume (sGMV), total white matter volume (WMV) and mean cortical thickness (CT), respectively. The age effects on total sGMV and WMV are estimated in 63 studies and the age effects on mean CT are estimated in 43 studies where mean CT is available. RESIs (blue) are estimated for cross-sectional studies, both CS-RESIs (blue) and L-RESIs (yellow) are estimated for longitudinal studies. The dots and lines represent the point estimates of the RESIs and their 95% confidence intervals (CIs). The blue dots and lines represent the CS-RESIs in longitudinal and RESIs in cross-sectional studies. The yellow dots and lines are for the L-RESIs in longitudinal studies. Arrows mean the upper limits of the CIs are greater than the axis boundary.

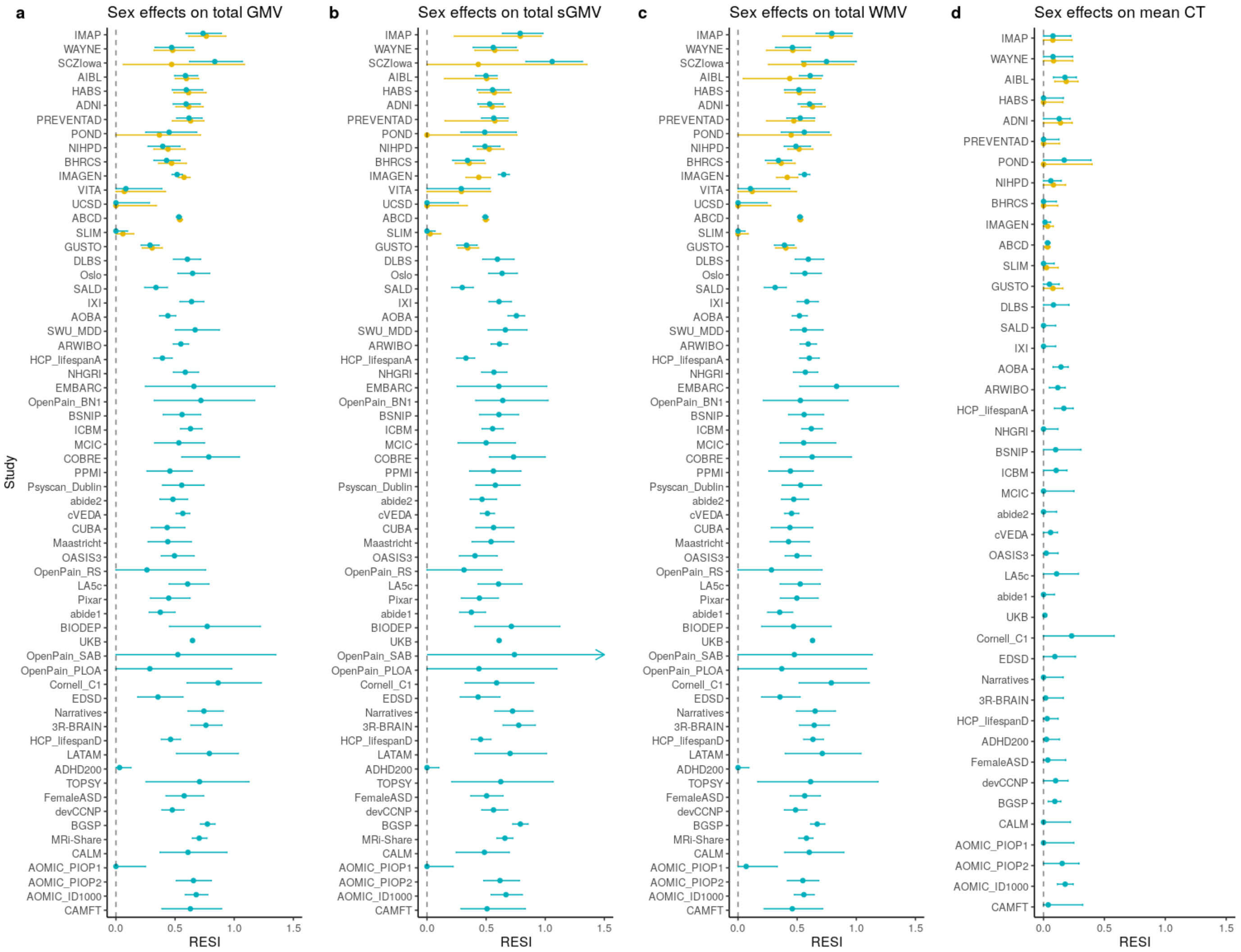


Figure SI9: Forest plots of sex effects on total gray matter volume (GMV), total subcortical gray matter volume (sGMV), total white matter volume (WMV) and mean cortical thickness (CT), respectively. The sex effects on total GMV, sGMV and WMV are estimated in 63 studies and the sex effects on mean CT are estimated in 43 studies where mean CT is available. Both CS-RESI (blue) and L-RESI (yellow) are estimated for longitudinal studies. The dots and lines represent the point estimates of the RESIs and their 95% confidence intervals (CIs). The blue dots and lines represent the CS-RESIs in longitudinal and RESIs in cross-sectional studies. The yellow dots and lines are for the L-RESIs in longitudinal studies. Arrow means the upper limit of the CI is greater than the axis boundary.

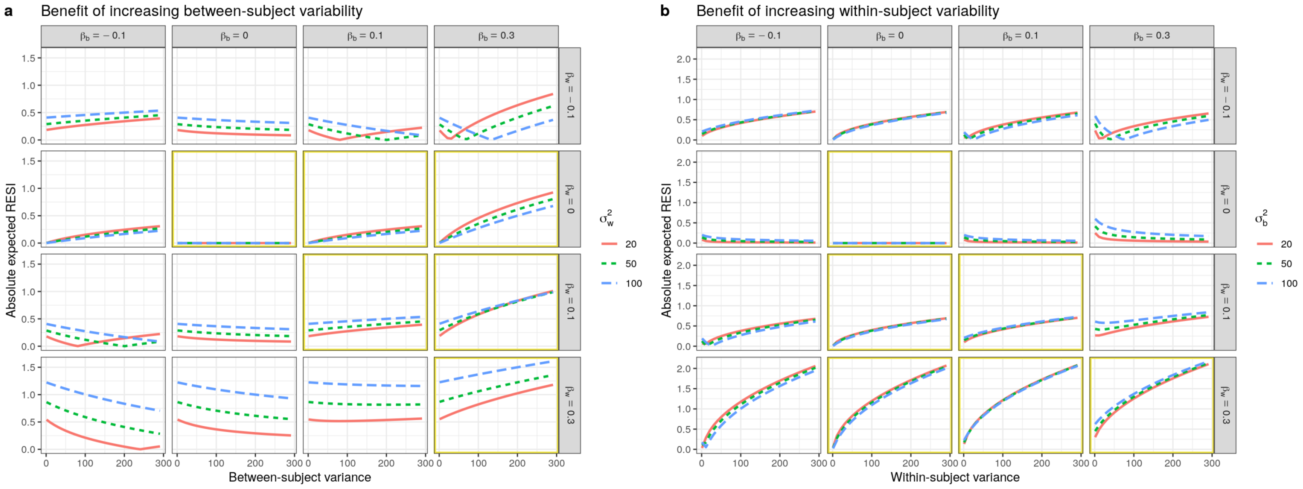


Figure SI10: The expected ESs for $\beta$ in model (12) assuming an exchangeable correlation structure. (a) The benefit of increasing between-subject variance is dependent on the between-subject parameter being larger and the same sign as the within-subject parameter (i.e., $\beta_w$; highlighted in yellow). (b) The benefit of increasing within-subject variance is dependent on the within-subject parameter being larger and the same sign as the between-subject parameter (i.e., $\beta_b$; highlighted in yellow). When equal ($\beta_b = \beta_w$; diagonal), there is a benefit to increasing either. Note, these changes to the ES are related to bias in the estimation of the parameter due to differing between- and within-subject effects. In all other cases, the curves are negative or non-monotonic. The formula for the curves is given in (14) and shown for $y^2 = 30$ and $\rho = 0.6$. Absolute expected ESs are shown here as their directions do not affect power.
