## Extended Display Item for "Study design features increase replicability in cross-sectional and longitudinal brain-wide association studies"

### Partial regression plots for the meta-analysis on the standardized effect sizes of age on regional measures

Kaidi Kang

7/30/2023

#### Cortical Thickness (CT)

##### Frontal Lobe (Left hemisphere)

**Caudal Anterior**

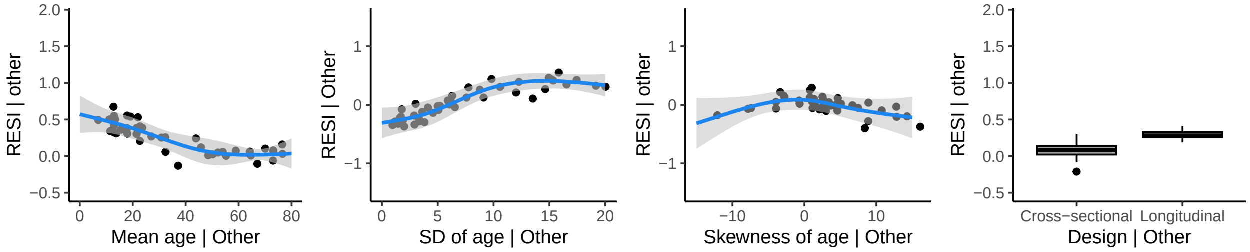


**Caudal Middle Frontal**

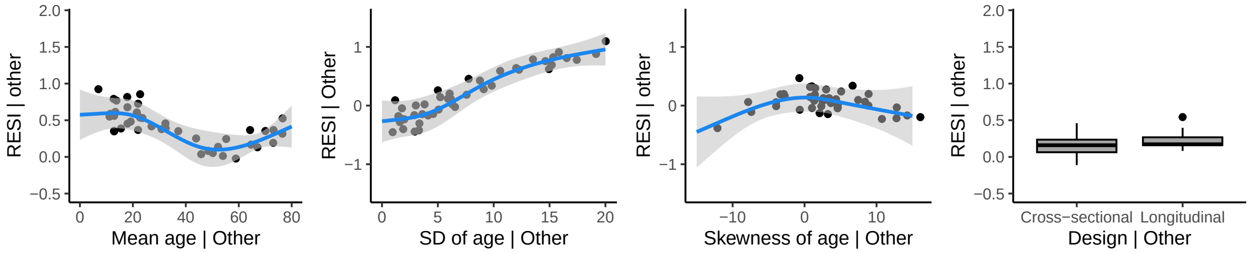


**Frontal Pole**

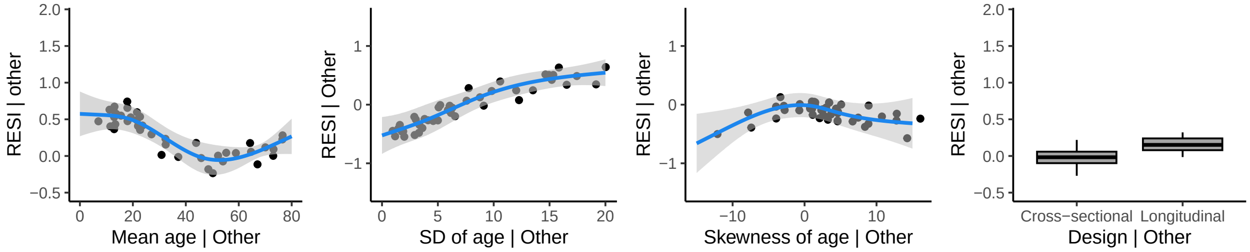

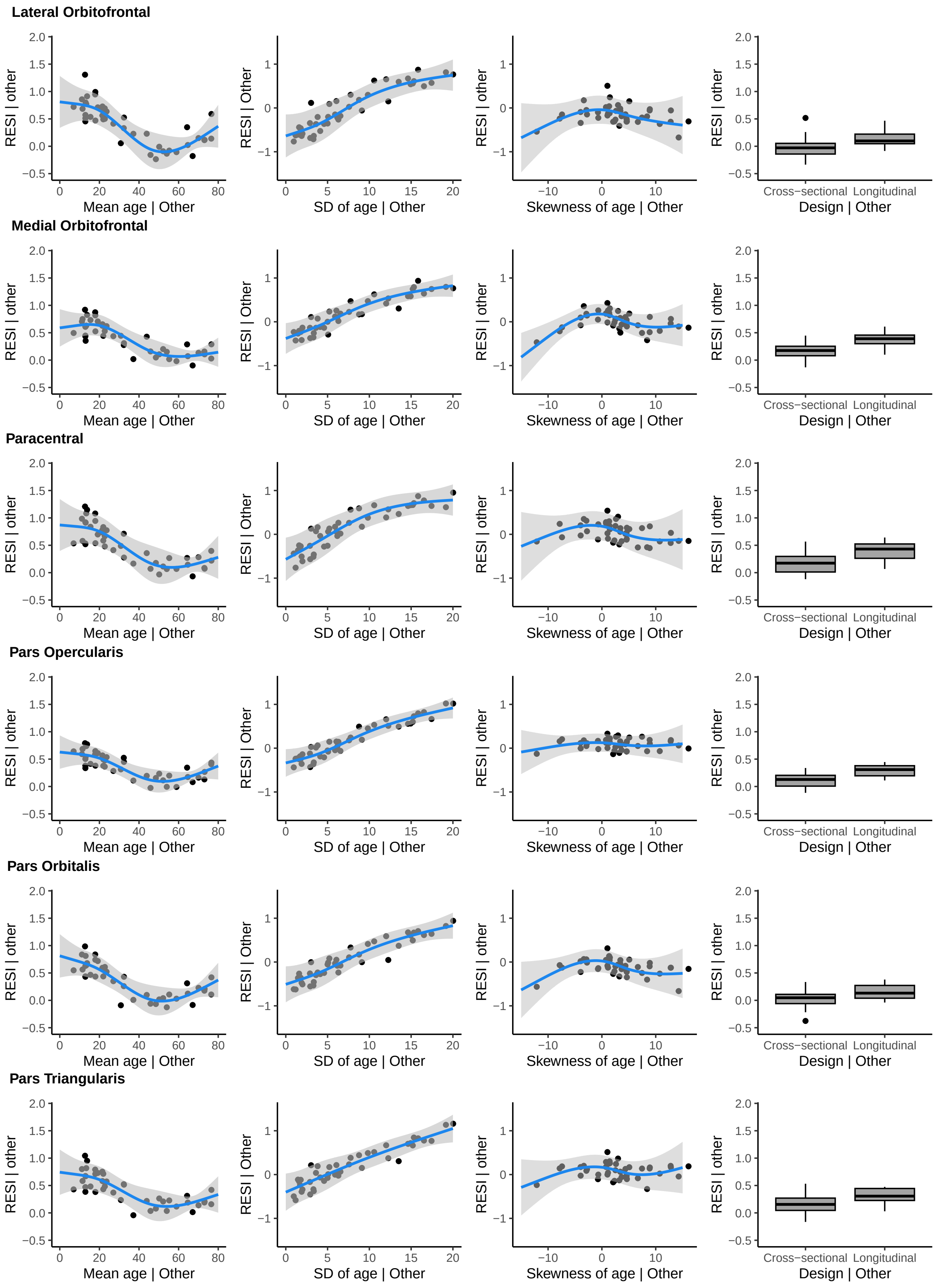

Lateral Orbitofrontal
Medial Orbitofrontal
Paracentral
Pars Opercularis
Pars Orbitalis
Pars Triangularis
RESI | other
RESI | Other
Mean age | Other
SD of age | Other
Skewness of age | Other
Design | Other
Cross−sectional
Longitudinal

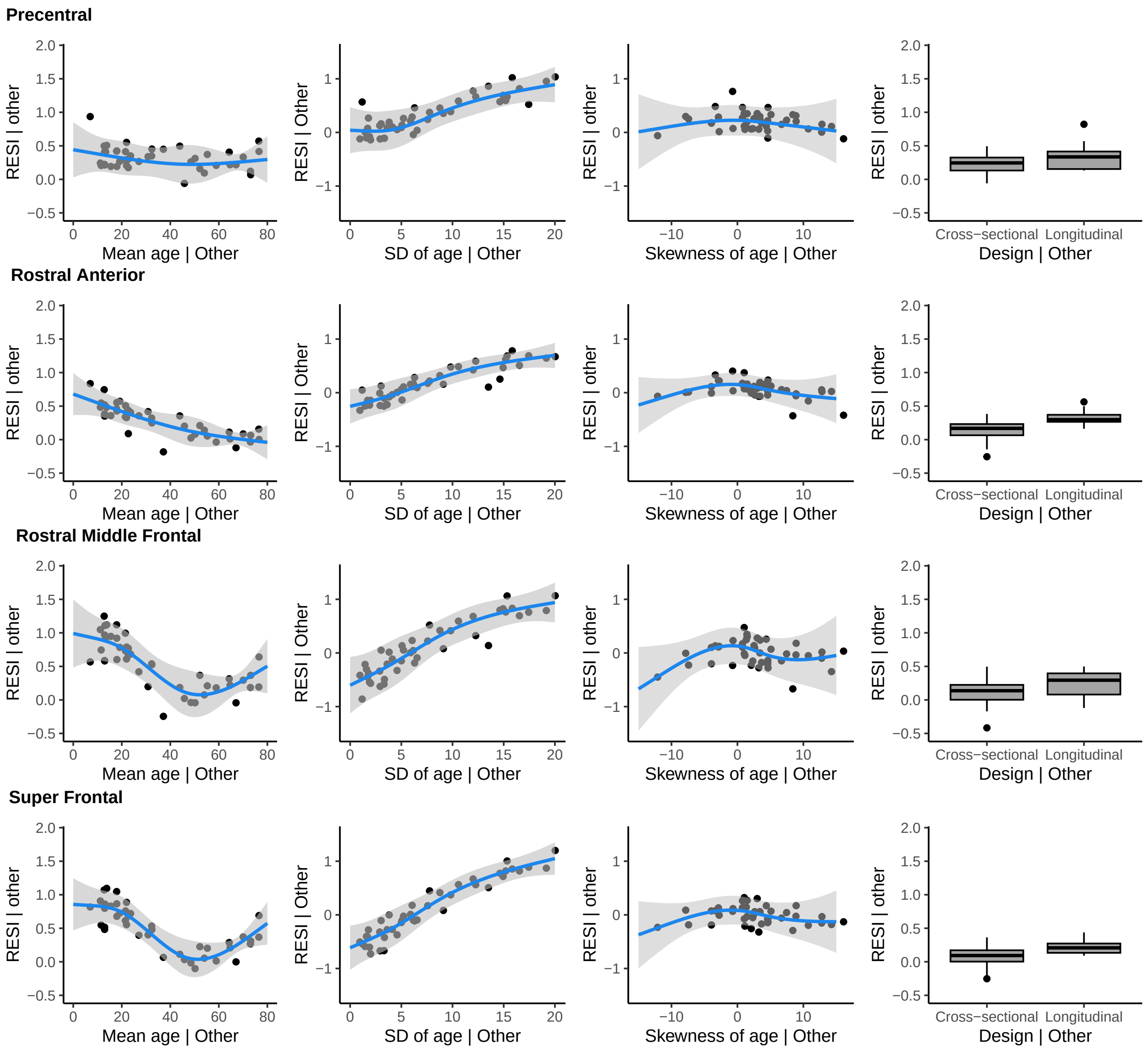
Precentral
Rostral Anterior
Rostral Middle Frontal
Super Frontal
RESI | other
RESI | Other
Mean age | Other
SD of age | Other
Skewness of age | Other
Design | Other
Cross−sectional
Longitudinal
2.0
1.5
1.0
0.5
0.0
−0.5
1
0
−1
0
20
40
60
80
5
10
15
−10

### Frontal Lobe (Right hemisphere)

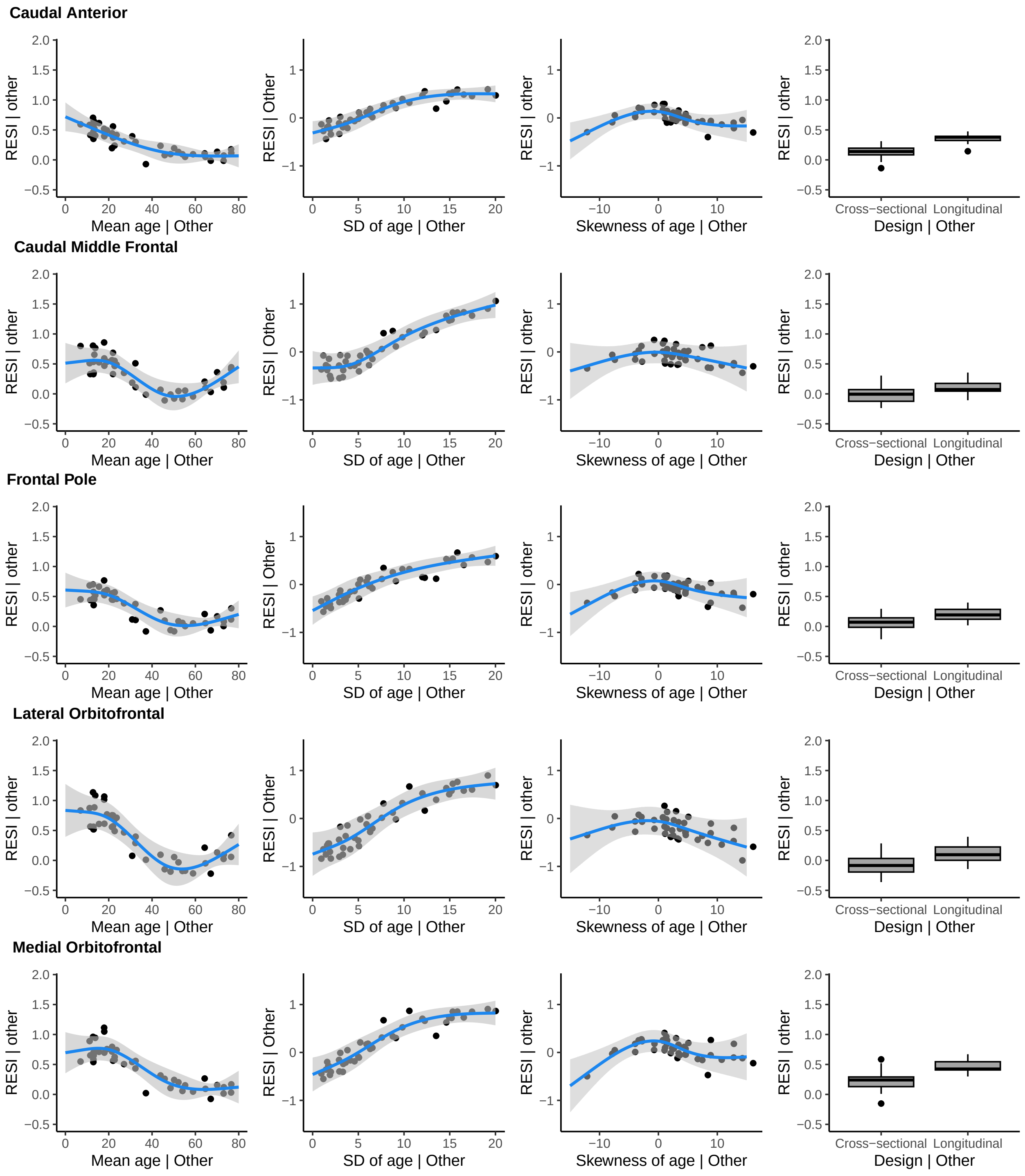

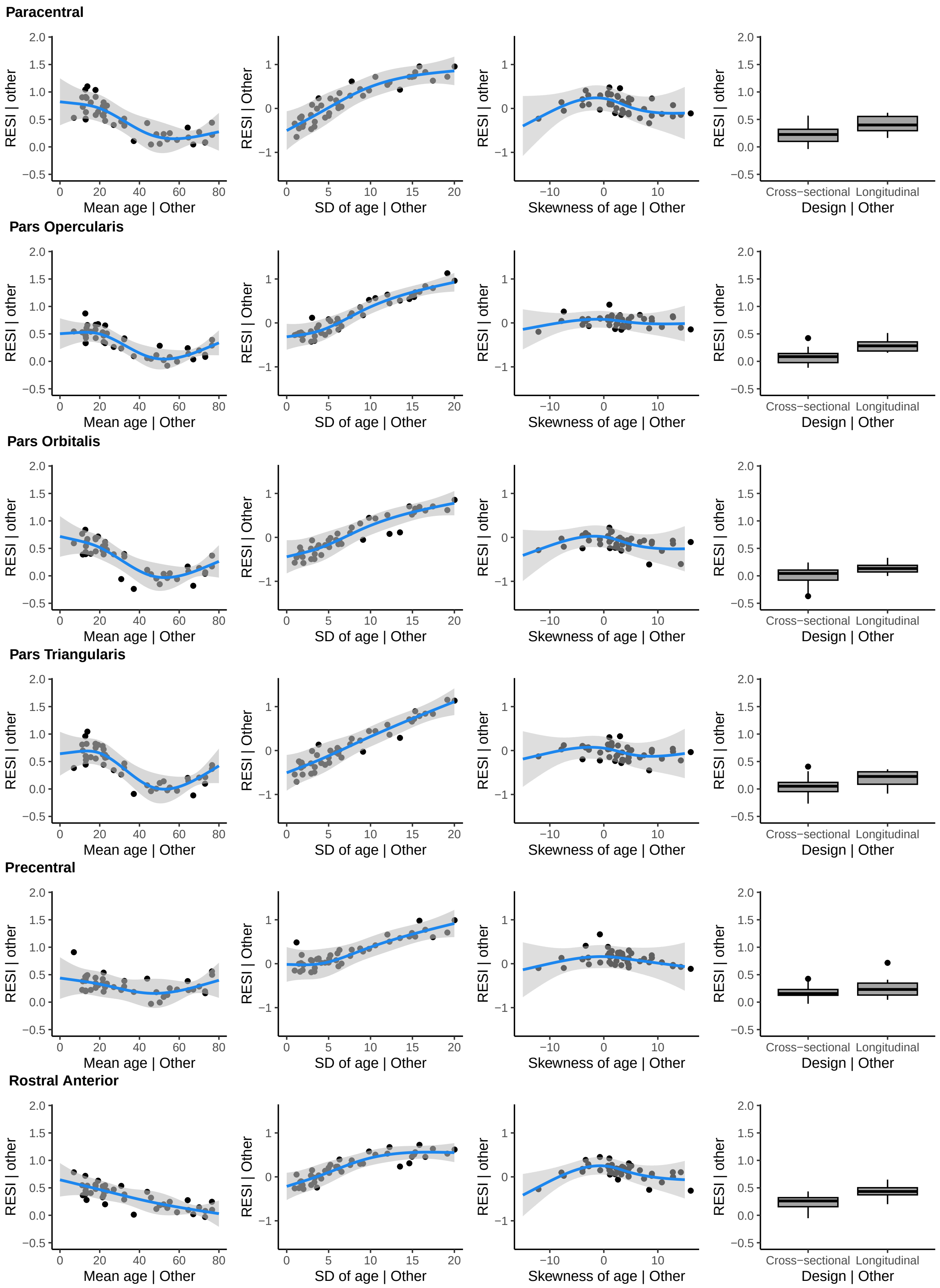

Paracentral
Pars Opercularis
Pars Orbitalis
Pars Triangularis
Precentral
Rostral Anterior
RESI | other
RESI | Other
Mean age | Other
SD of age | Other
Skewness of age | Other
Design | Other
Cross−sectional
Longitudinal

**Rostral Middle Frontal**

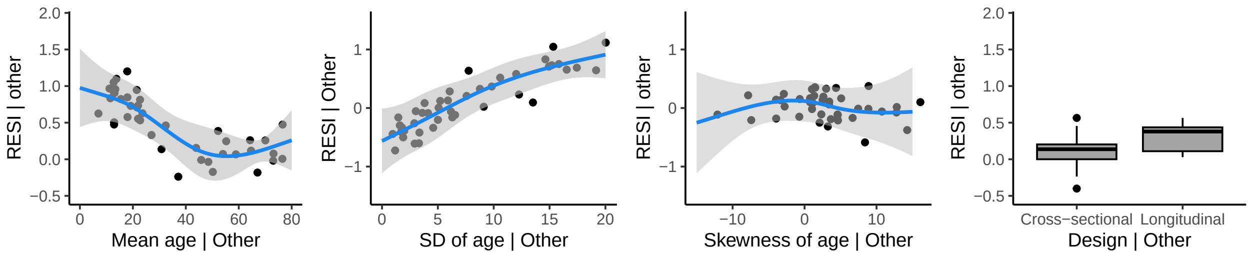


**Super Frontal**

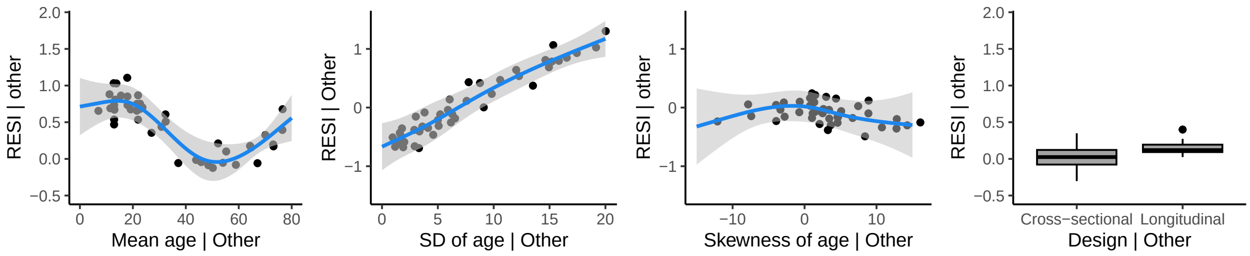


### Occipital Lobe (Left hemisphere)

**Cuneus**

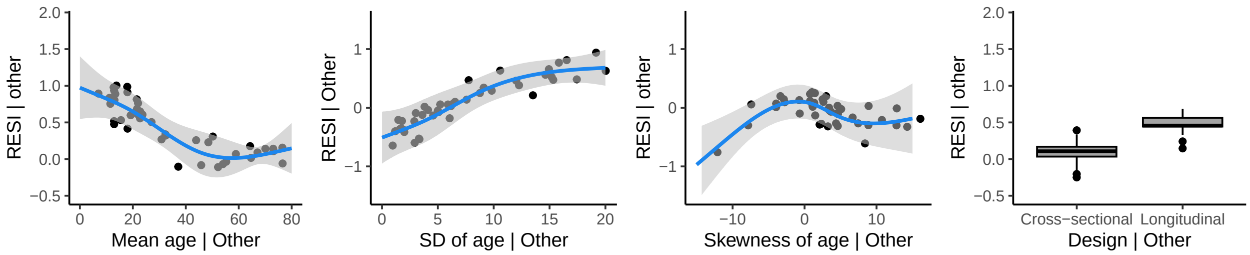


**Lateral Occipital**

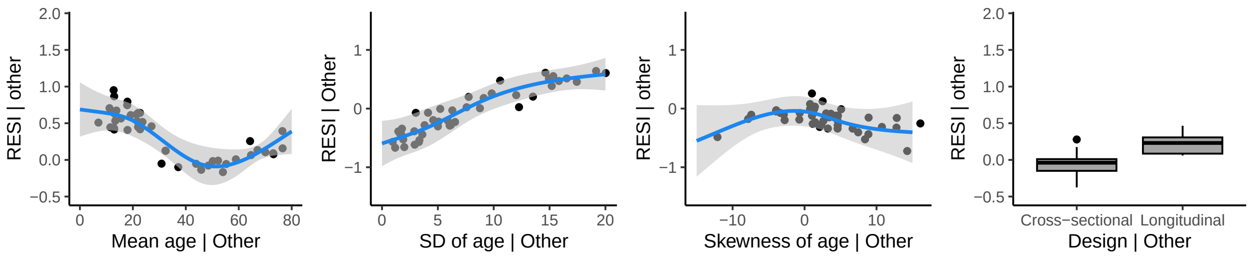


**Lingual**

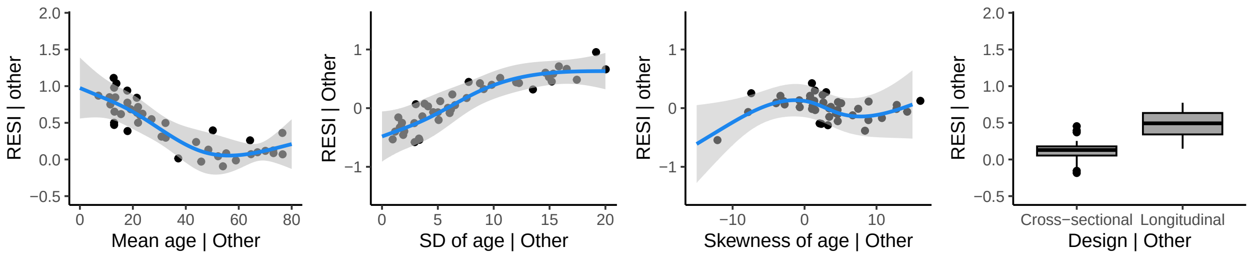

**Pericalcarine**

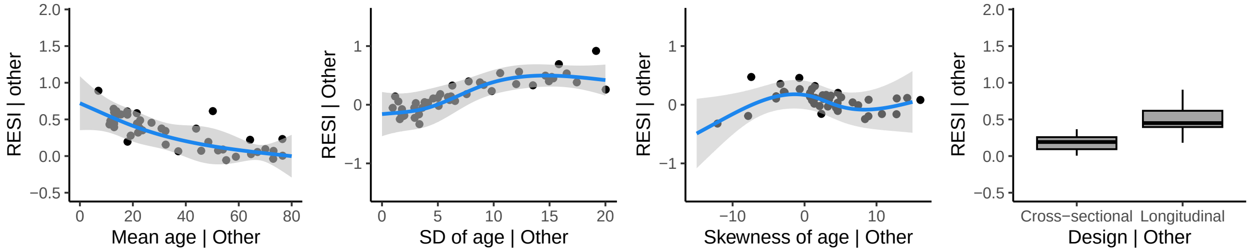


#### Occipital Lobe (Right hemisphere)

**Cuneus**

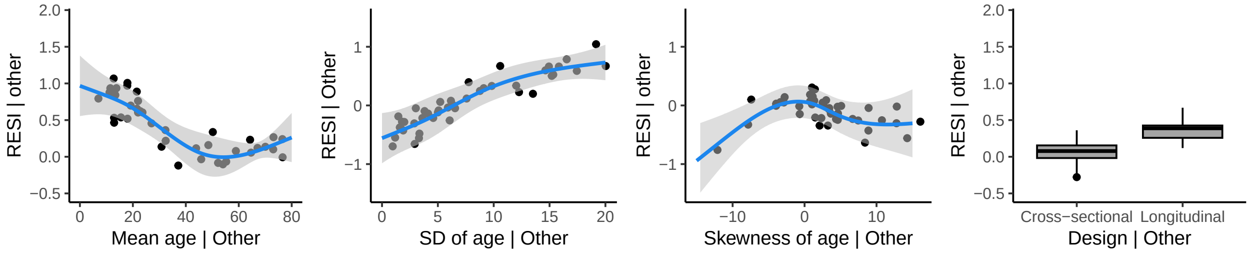


**Lateral Occipital**

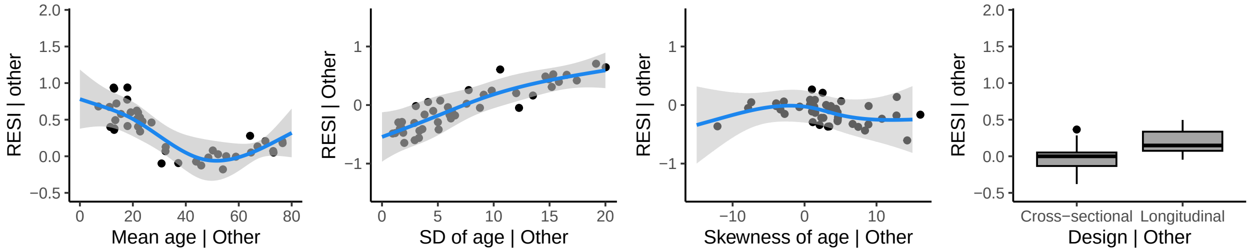


**Lingual**

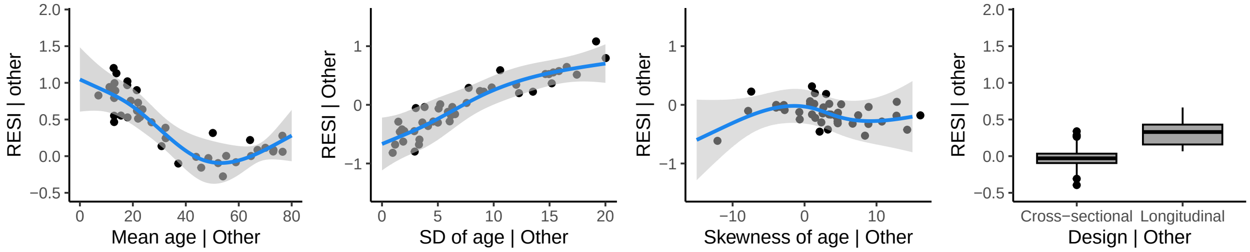


**Pericalcarine**

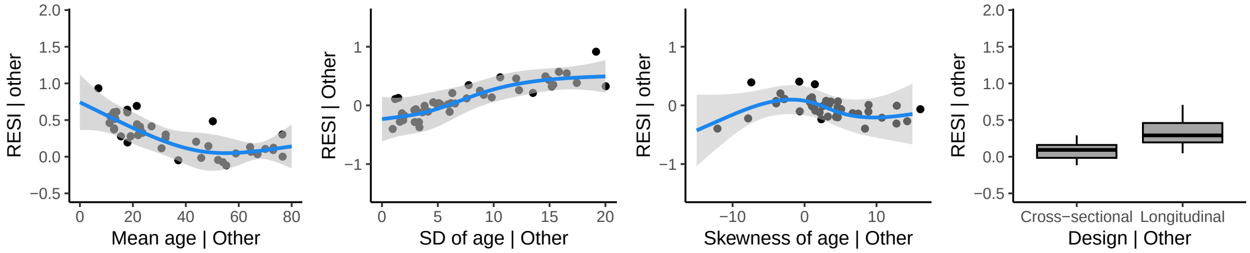

### Parietal Lobe (Left hemisphere)

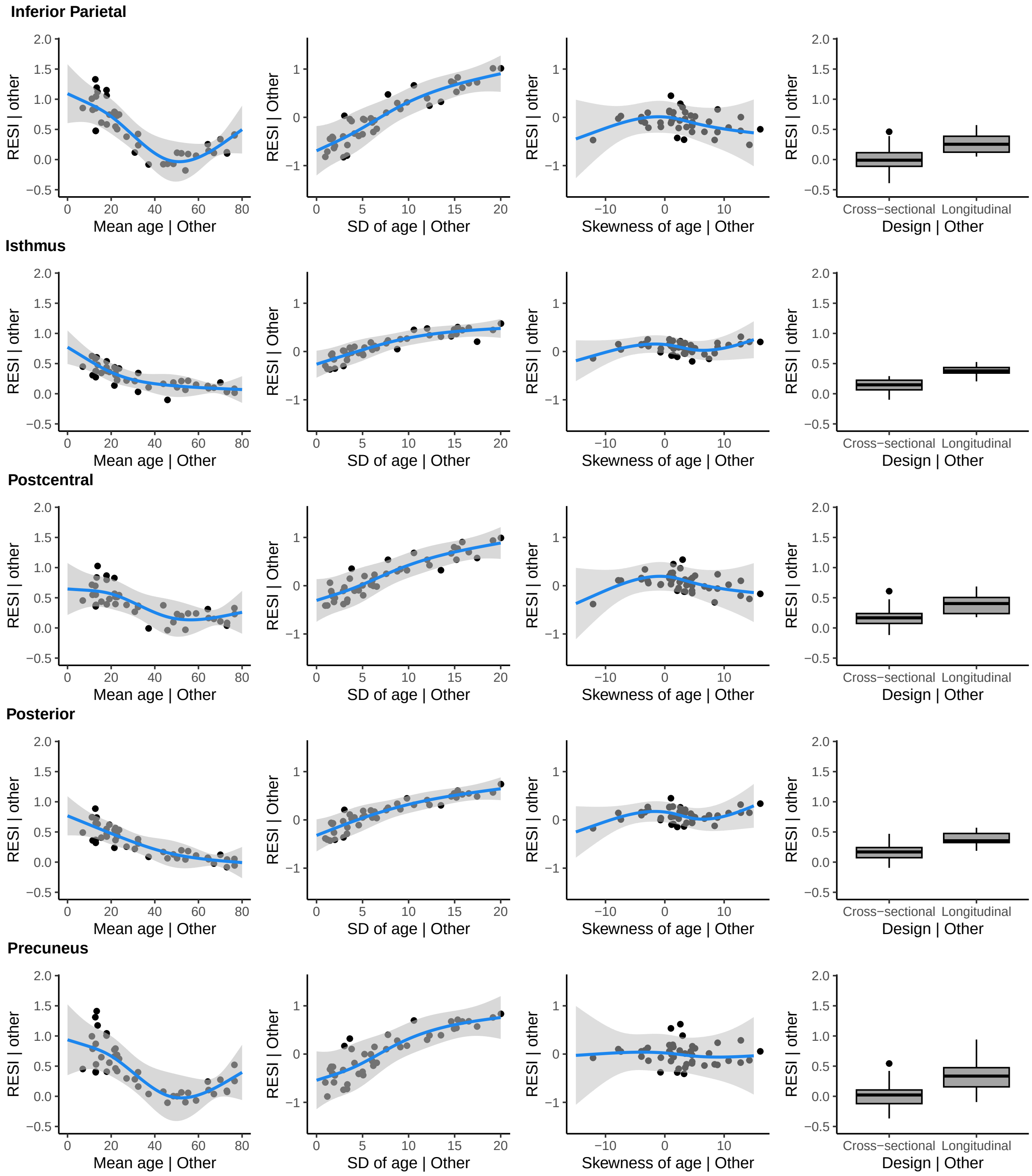

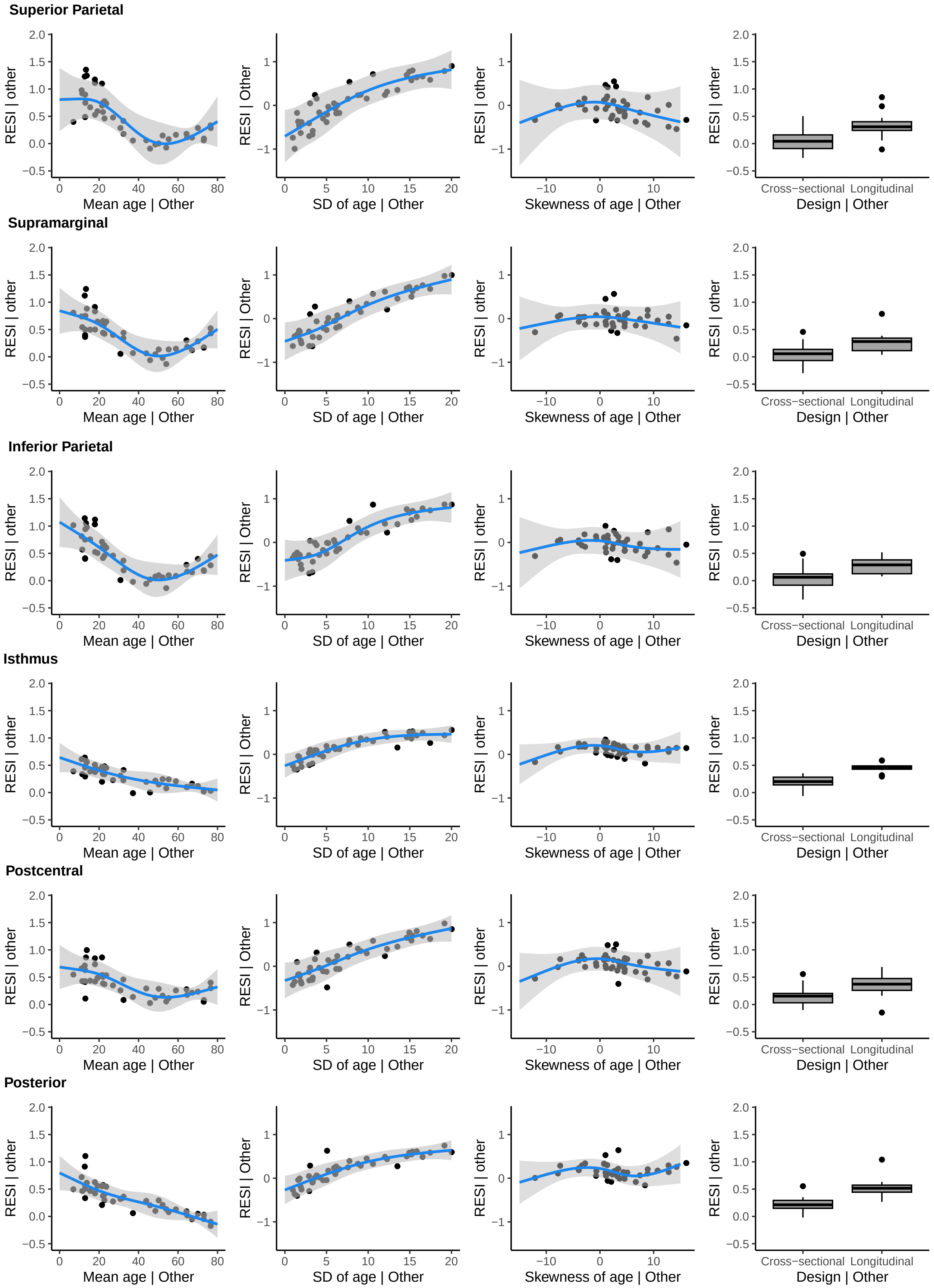
Superior Parietal
Supramarginal
Inferior Parietal
Isthmus
Postcentral
Posterior
RESI | other
RESI | Other
Mean age | Other
SD of age | Other
Skewness of age | Other
Design | Other
Cross−sectional
Longitudinal

**Precuneus**

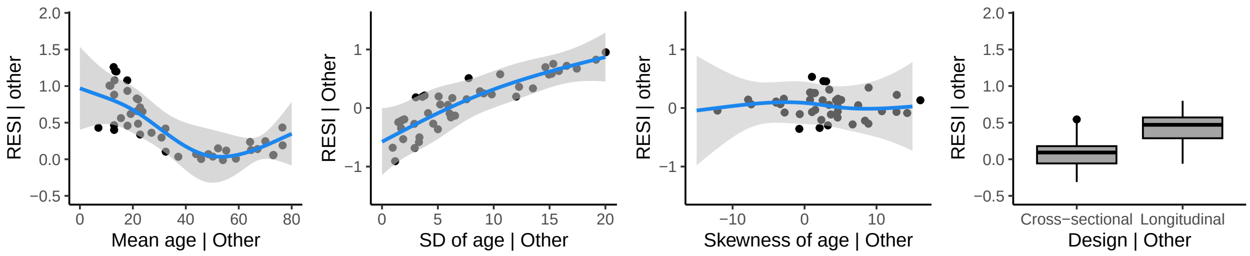


**Superior Parietal**

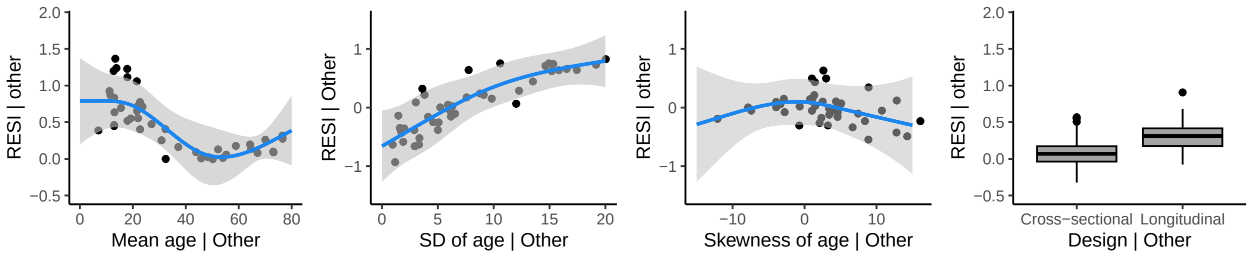


**Supramarginal**

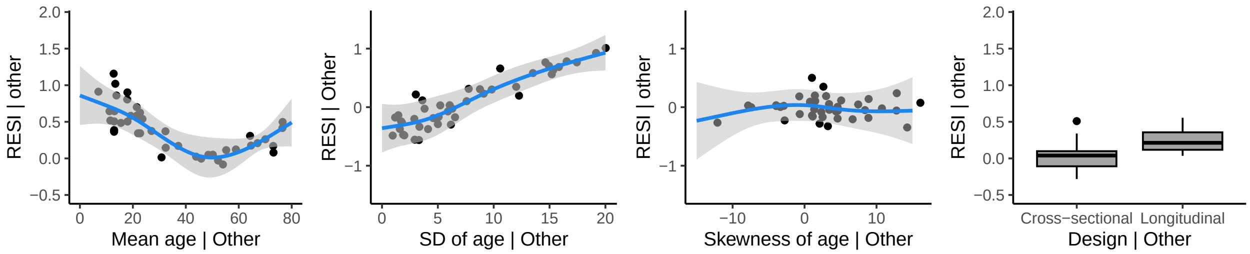


### Temporal Lobe (Left hemisphere)

**Banks of the Superior Temporal Sulcus**

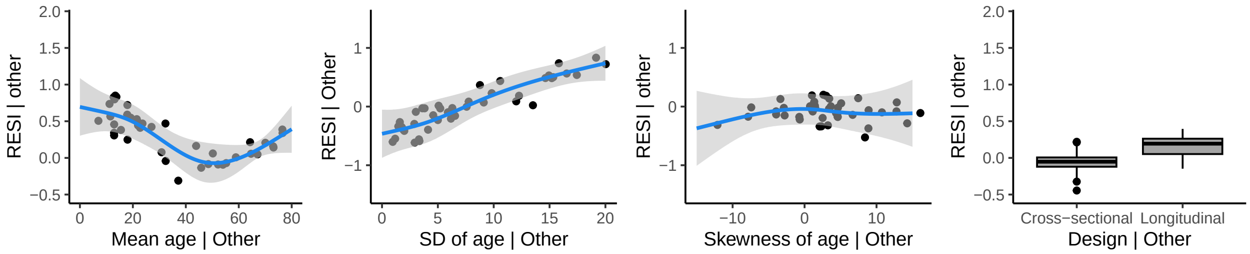


**Entorhinal**

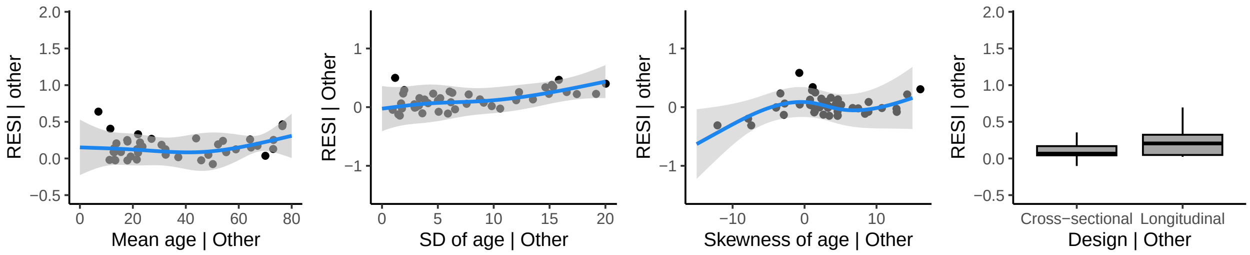

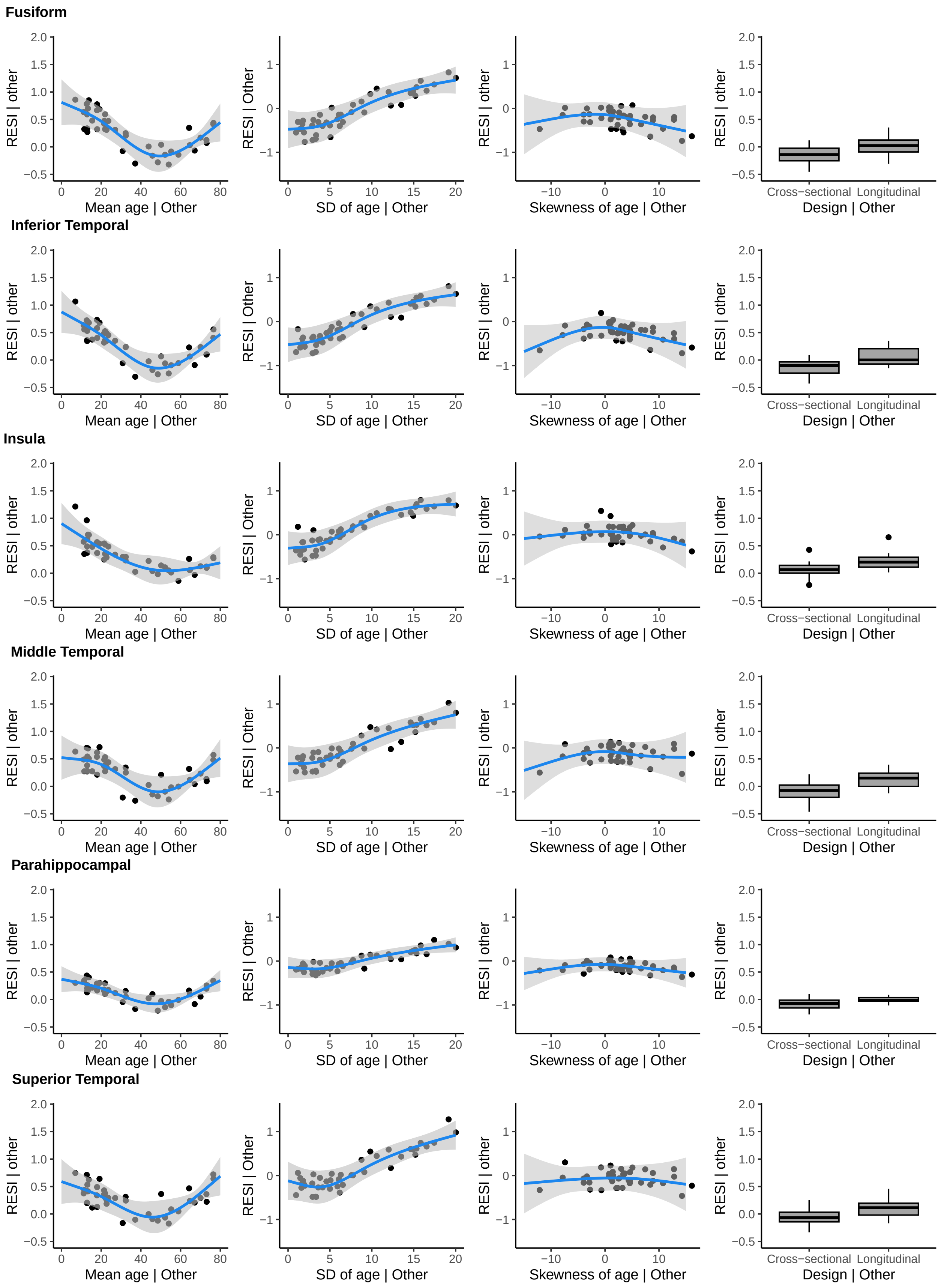
Fusiform
Inferior Temporal
Insula
Middle Temporal
Parahippocampal
Superior Temporal
RESI | other
RESI | Other
Mean age | Other
SD of age | Other
Skewness of age | Other
Design | Other
Cross−sectional
Longitudinal

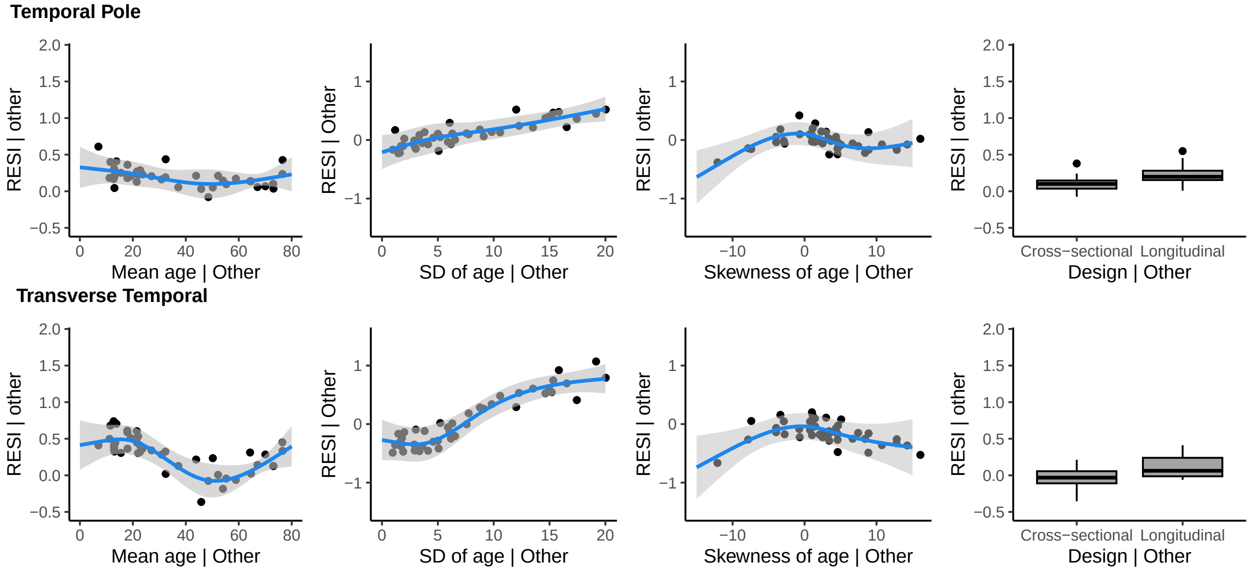


### Temporal Lobe (Right hemisphere)

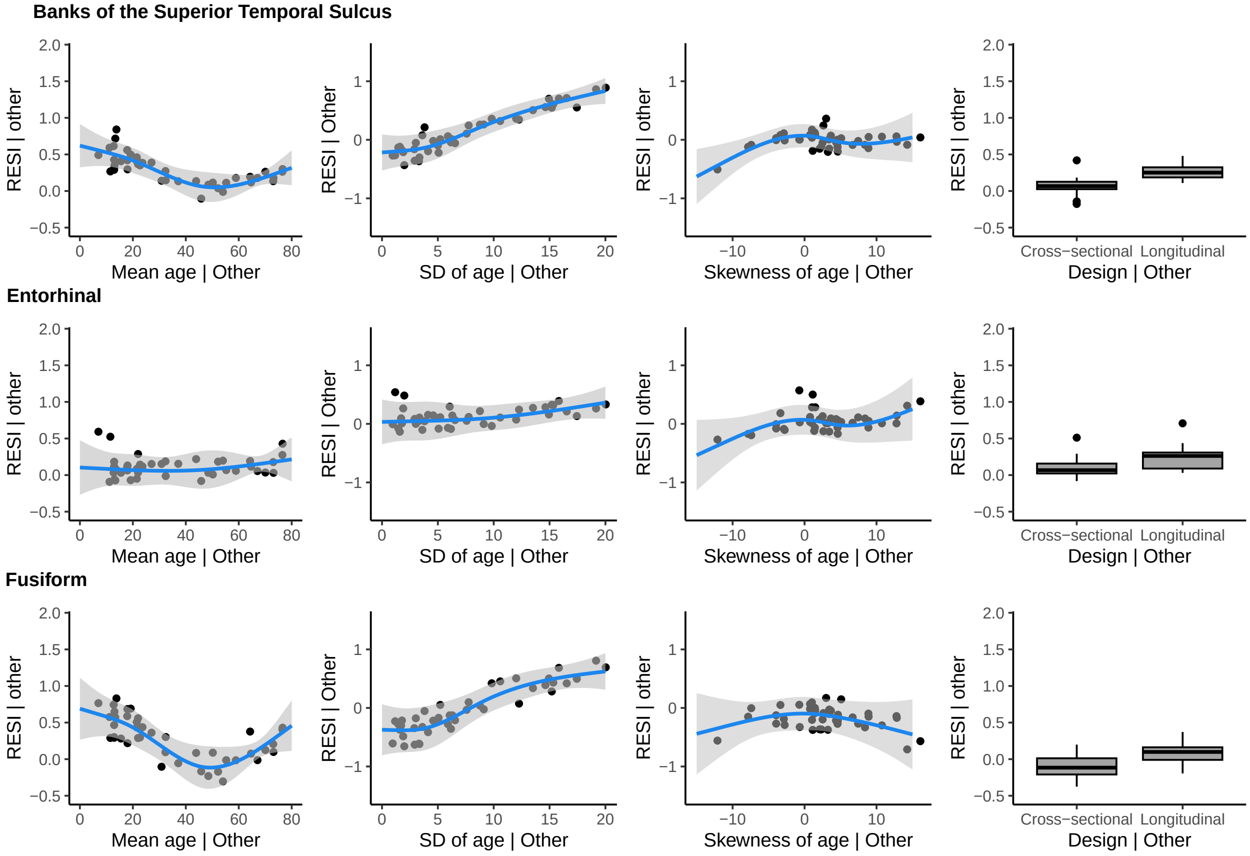

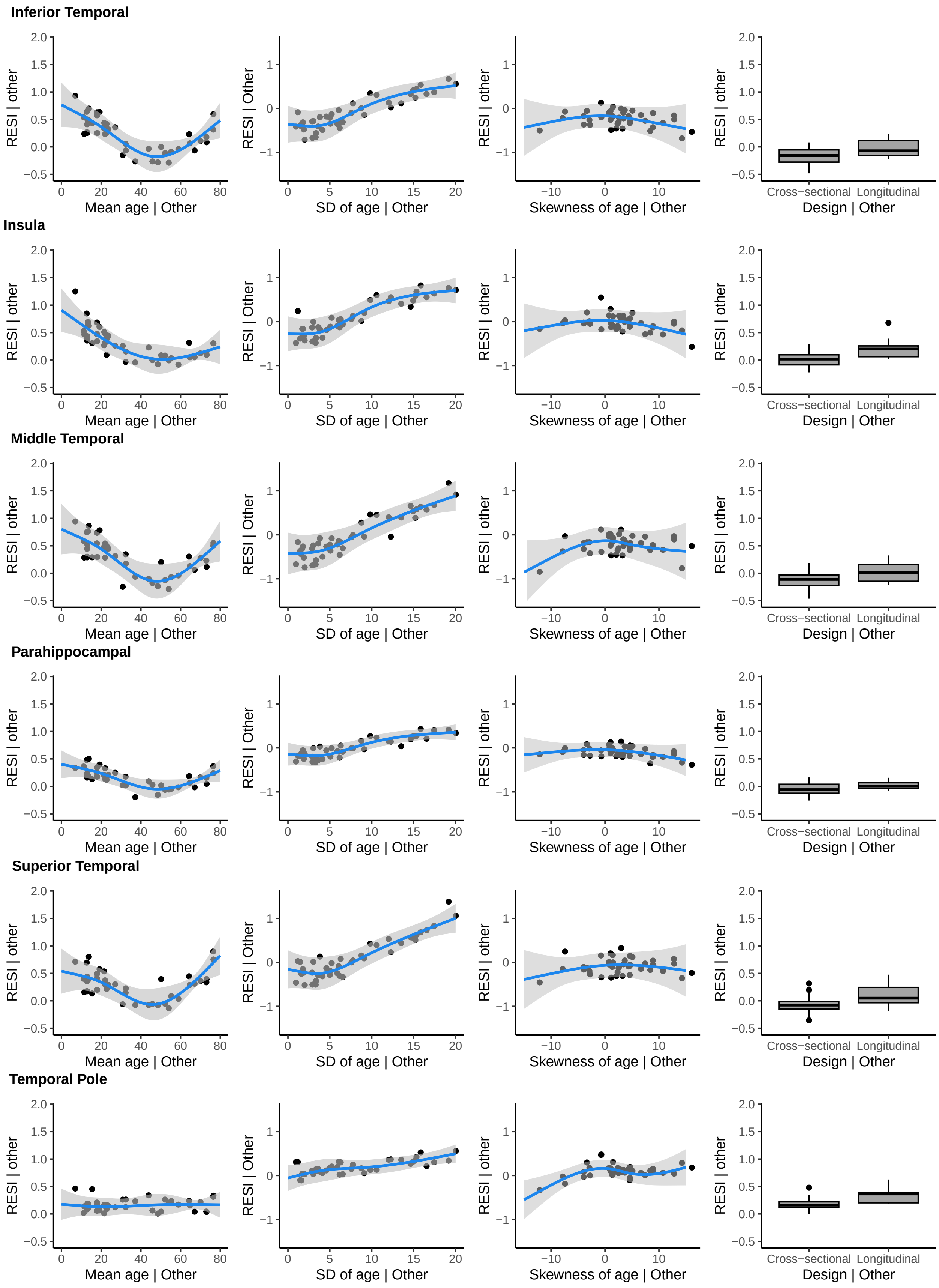
Inferior Temporal
Insula
Middle Temporal
Parahippocampal
Superior Temporal
Temporal Pole
RESI | other
RESI | Other
Mean age | Other
SD of age | Other
Skewness of age | Other
Design | Other
Cross−sectional
Longitudinal

**Transverse Temporal**

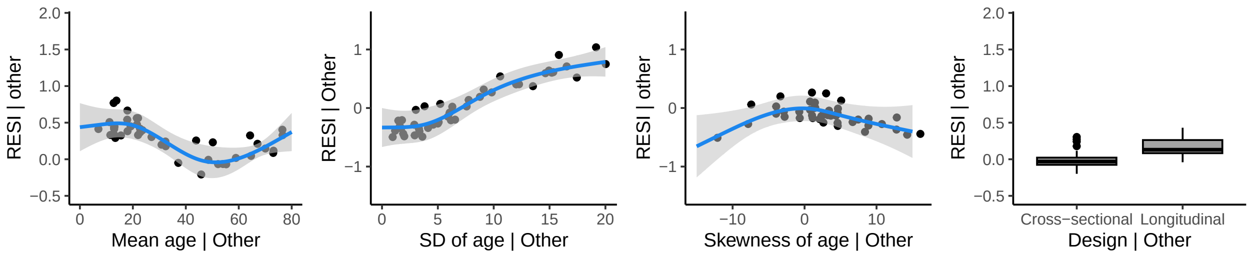


### Gray Matter Volume (GMV)

#### Frontal Lobe (Left hemisphere)

**Caudal Anterior**

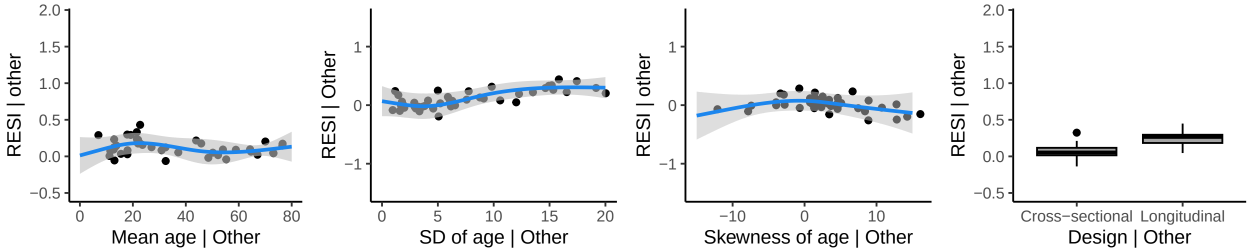


**Caudal Middle Frontal**

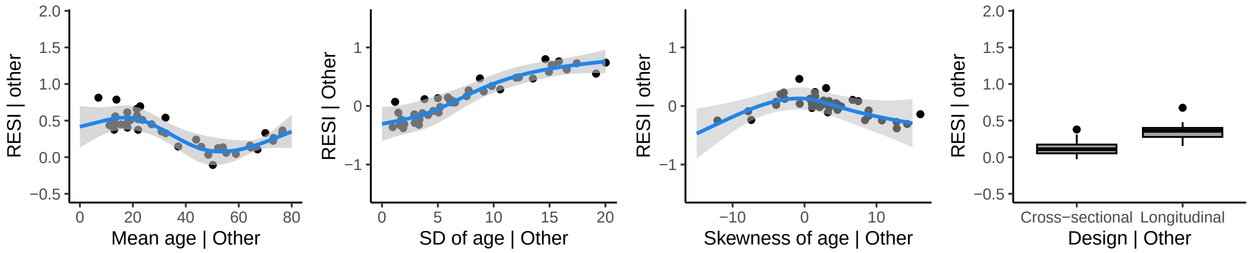


**Frontal Pole**

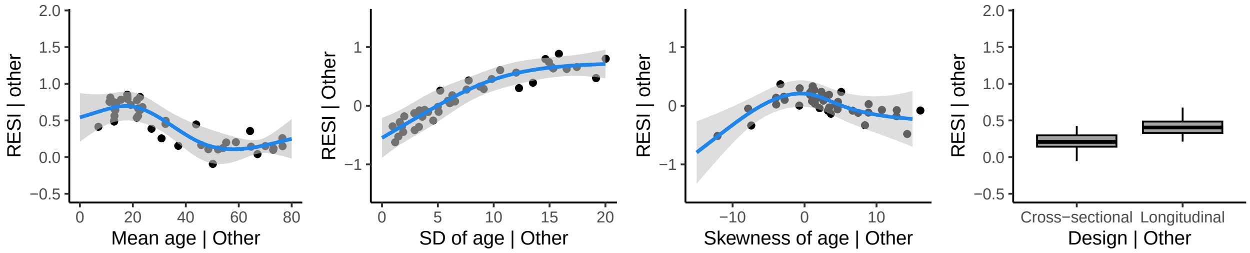


**Lateral Orbitofrontal**

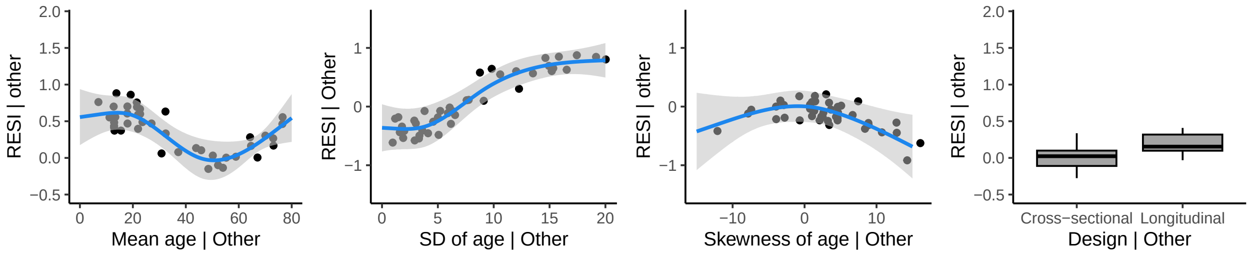

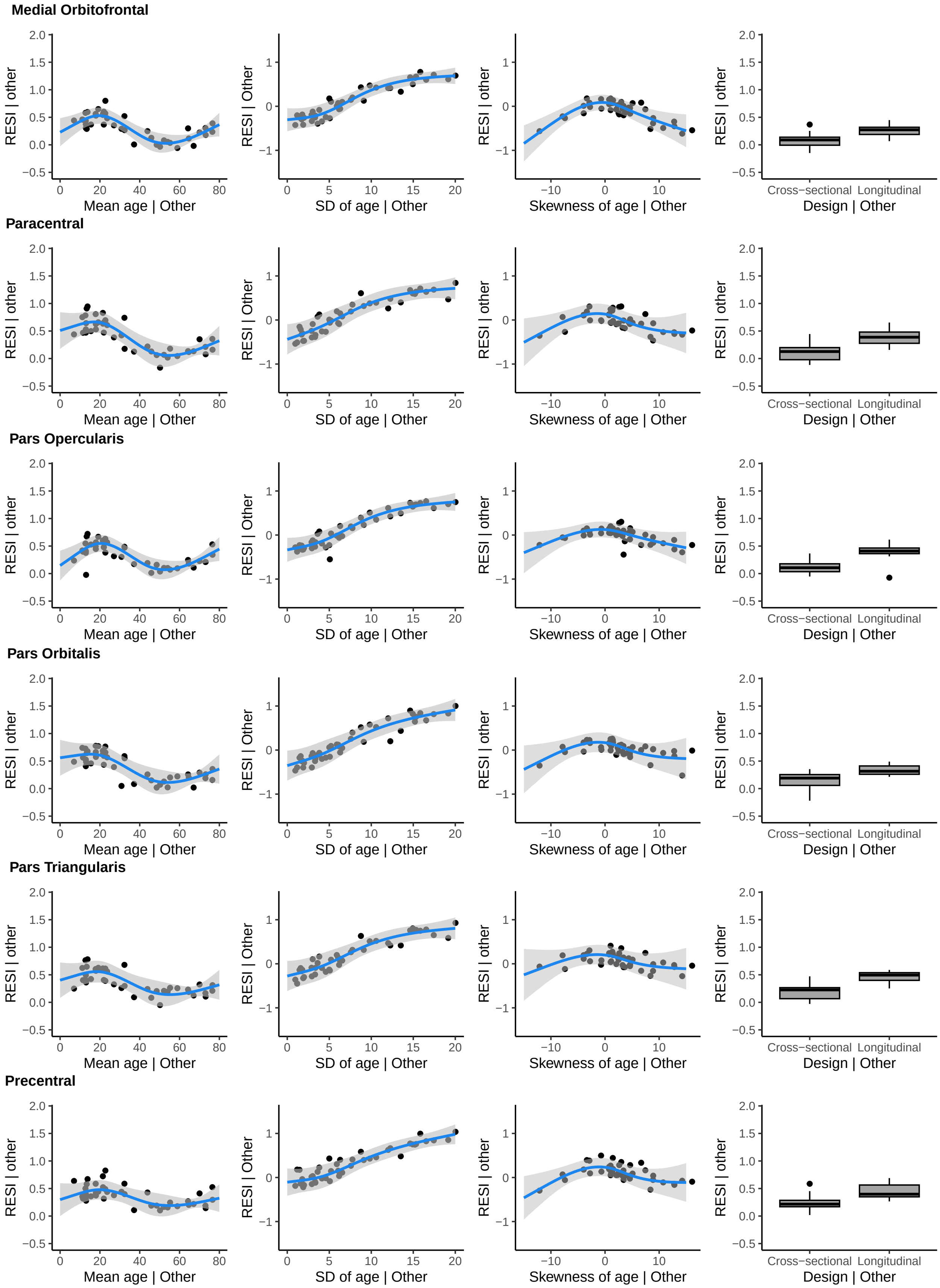
Medial Orbitofrontal
Paracentral
Pars Opercularis
Pars Orbitalis
Pars Triangularis
Precentral
RESI | other
RESI | Other
Mean age | Other
SD of age | Other
Skewness of age | Other
Design | Other
Cross−sectional
Longitudinal

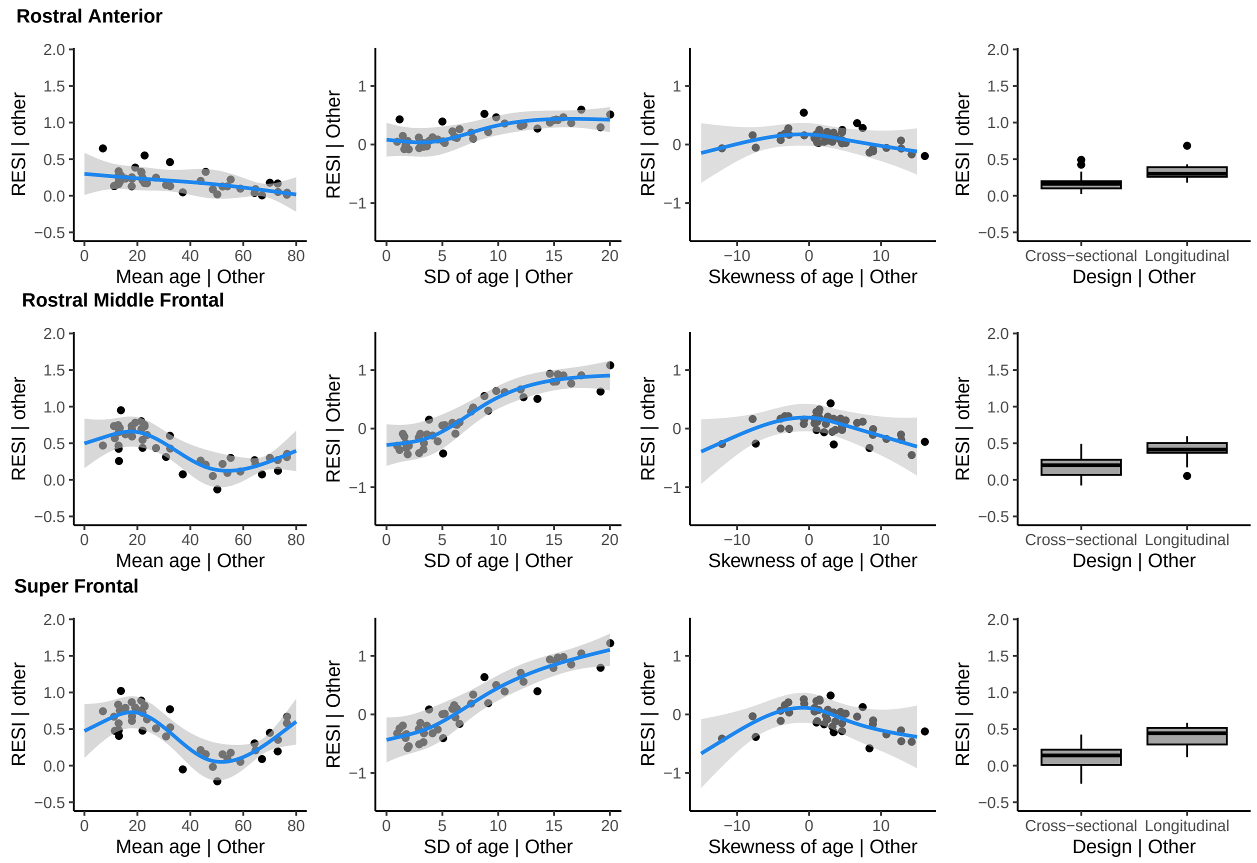


### Frontal Lobe (Right hemisphere)

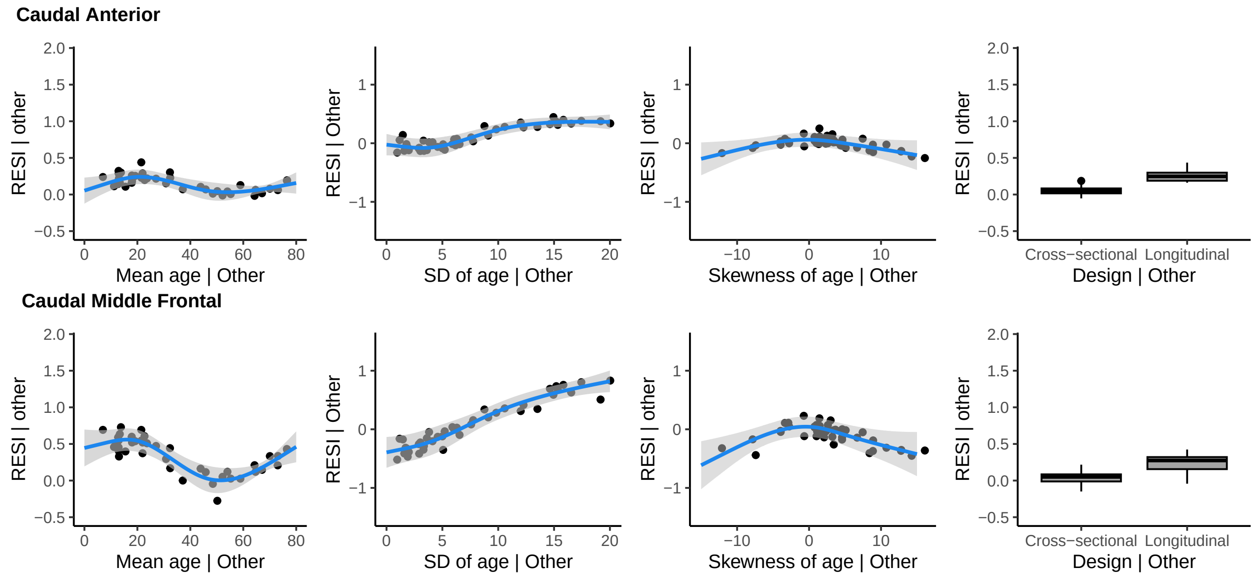

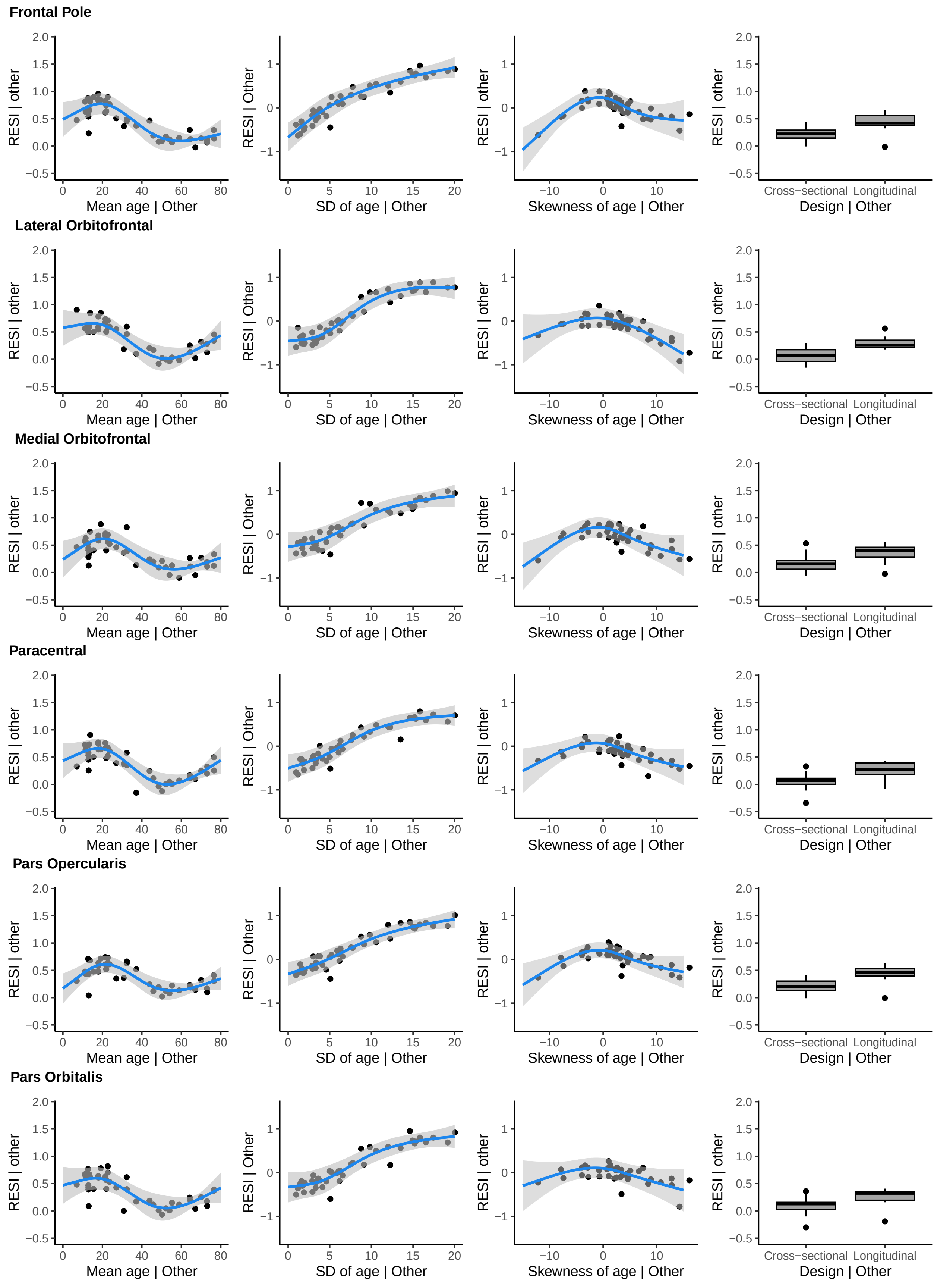
Frontal Pole
Lateral Orbitofrontal
Medial Orbitofrontal
Paracentral
Pars Opercularis
Pars Orbitalis
RESI | other
RESI | Other
Mean age | Other
SD of age | Other
Skewness of age | Other
Cross−sectional
Longitudinal
Design | Other

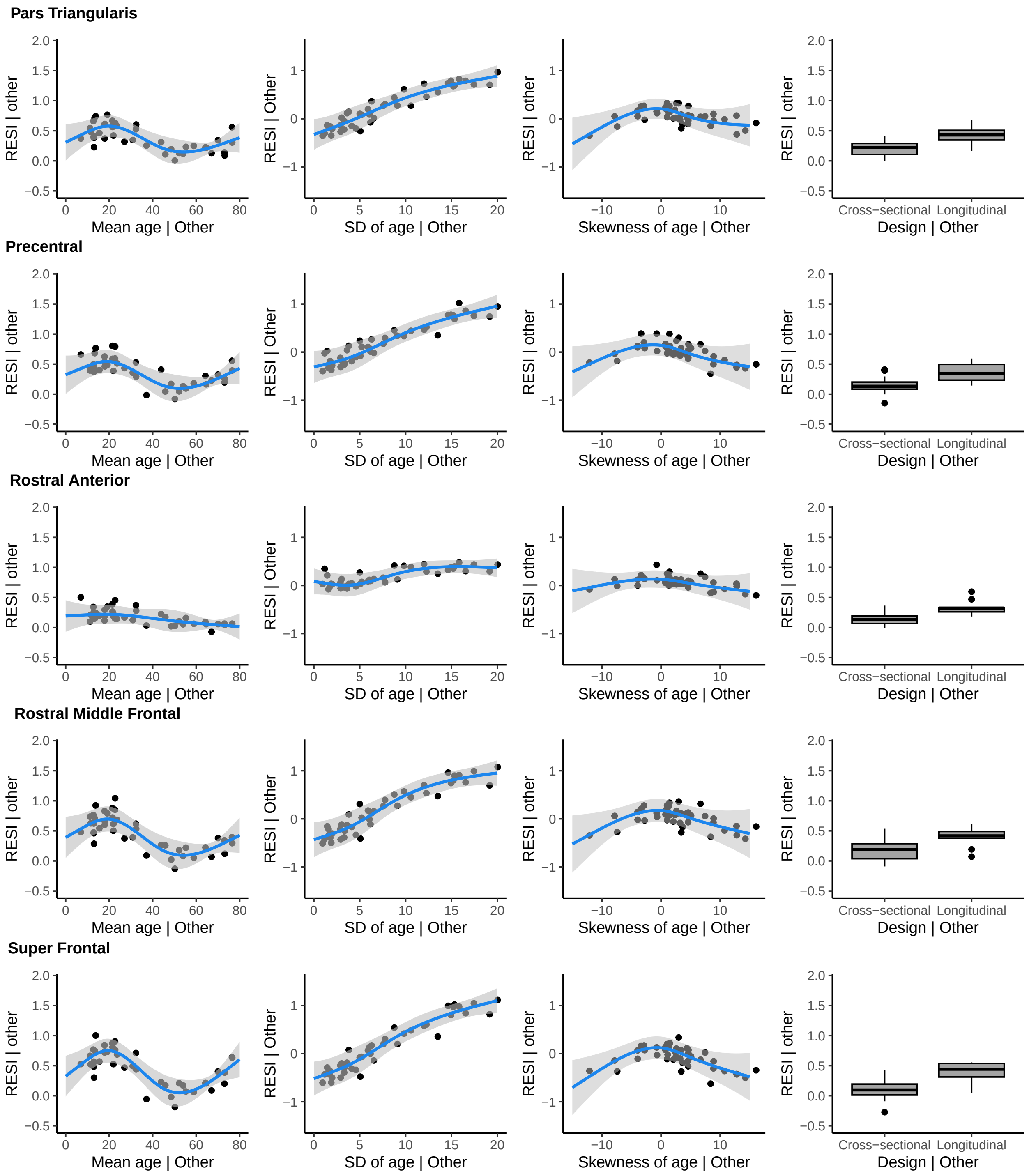
Pars Triangularis
Precentral
Rostral Anterior
Rostral Middle Frontal
Super Frontal
RESI | other
RESI | Other
Mean age | Other
SD of age | Other
Skewness of age | Other
Design | Other
Cross−sectional
Longitudinal

### Occipital Lobe (Left hemisphere)

**Cuneus**

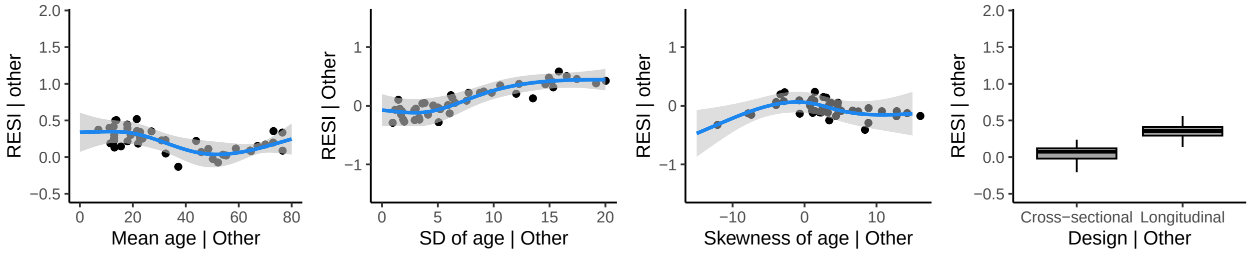


**Lateral Occipital**

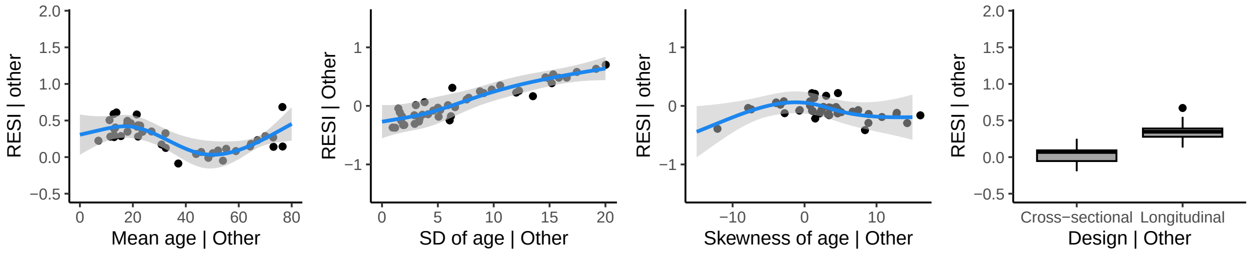


**Lingual**

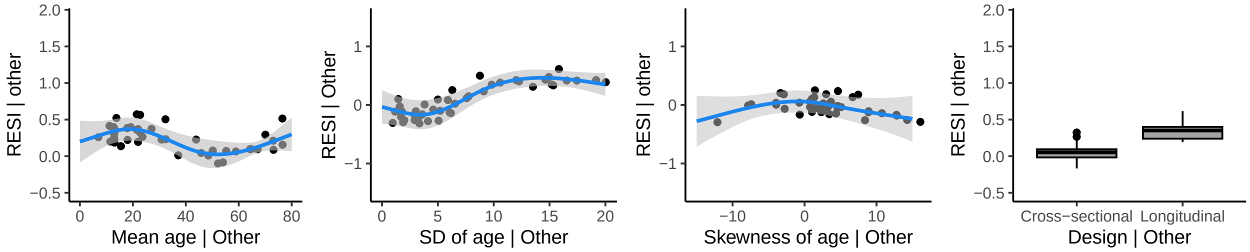


**Pericalcarine**

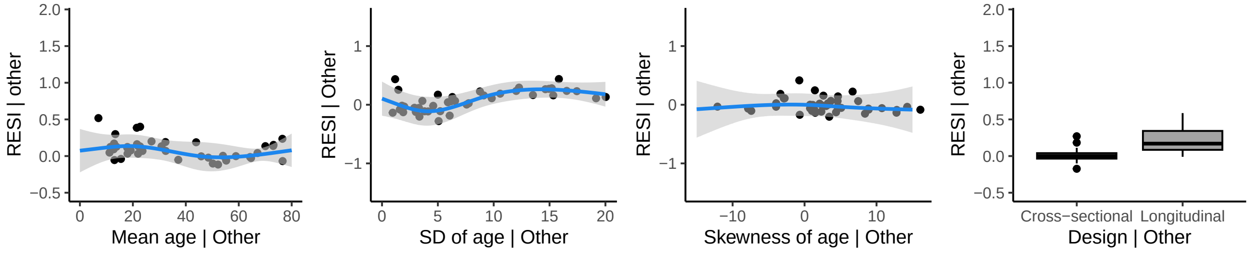

### Occipital Lobe (Right hemisphere)

**Cuneus**

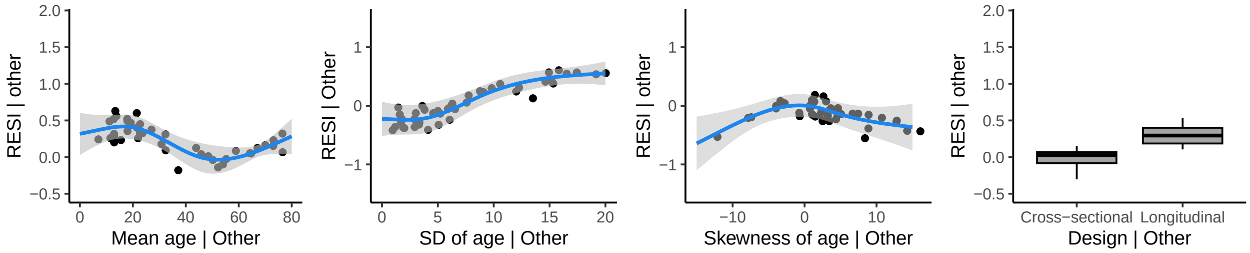


**Lateral Occipital**

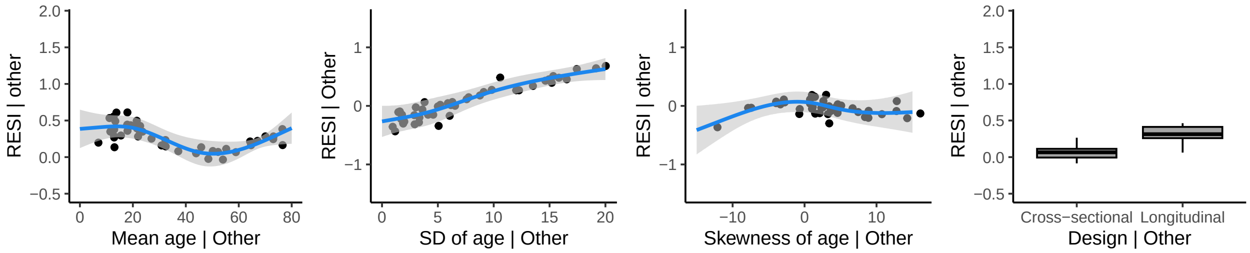


**Lingual**

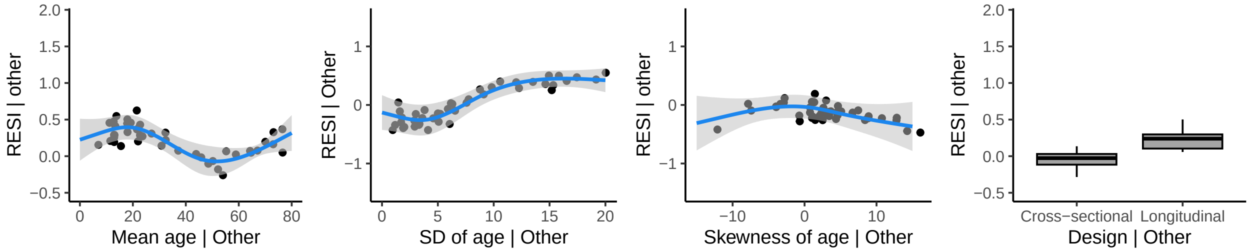


**Pericalcarine**

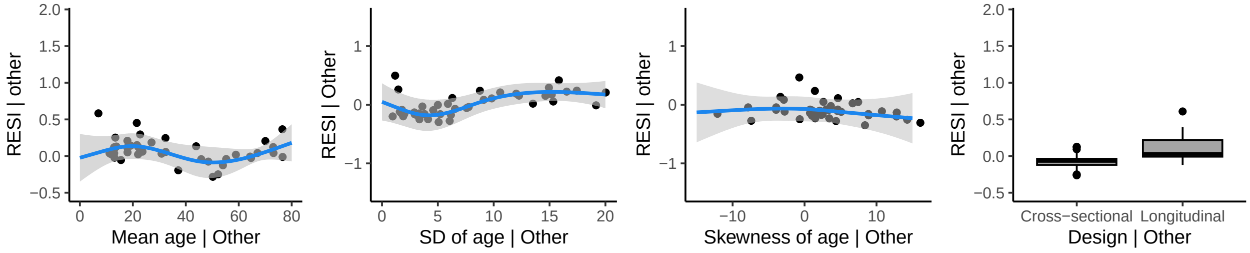

### Parietal Lobe (Left hemisphere)

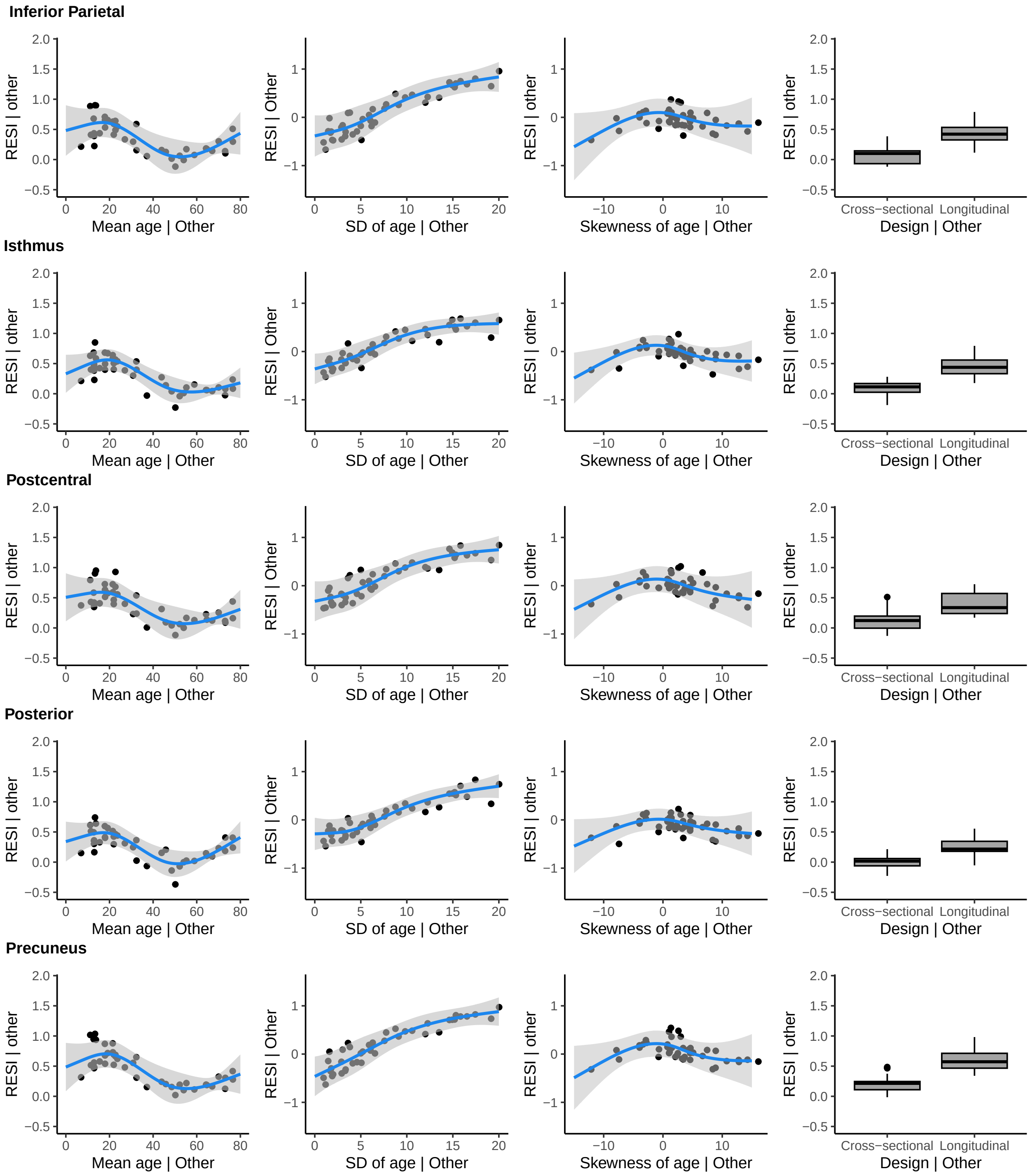

**Superior Parietal**

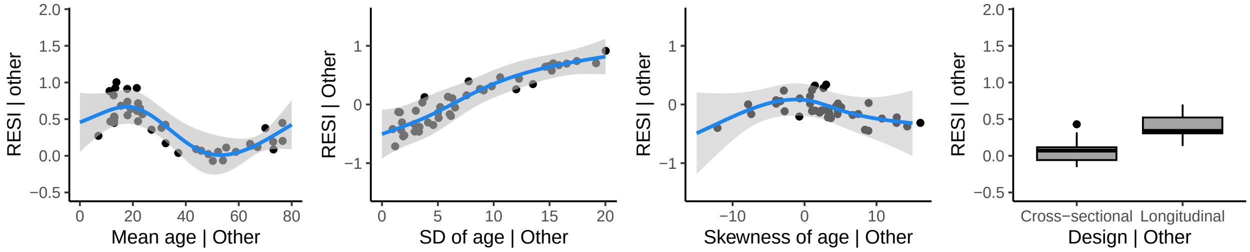


**Supramarginal**

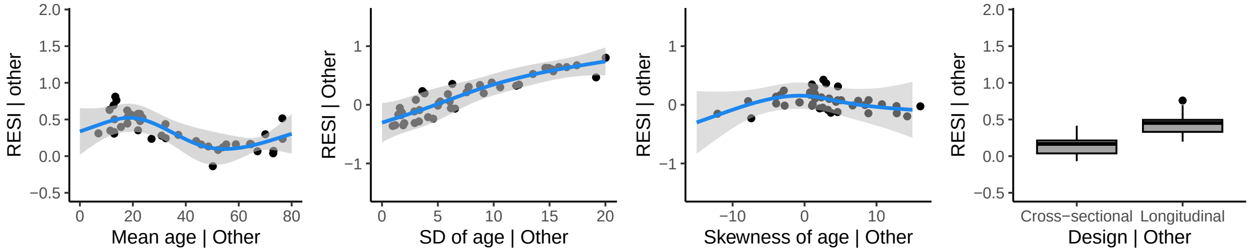


### Parietal Lobe (Right hemisphere)

**Inferior Parietal**

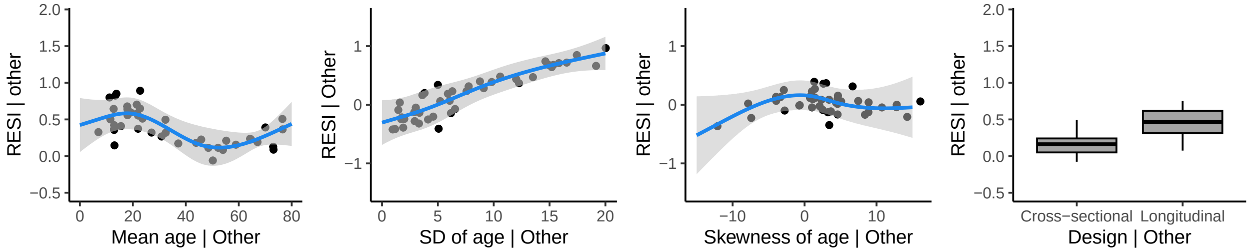


**Isthmus**

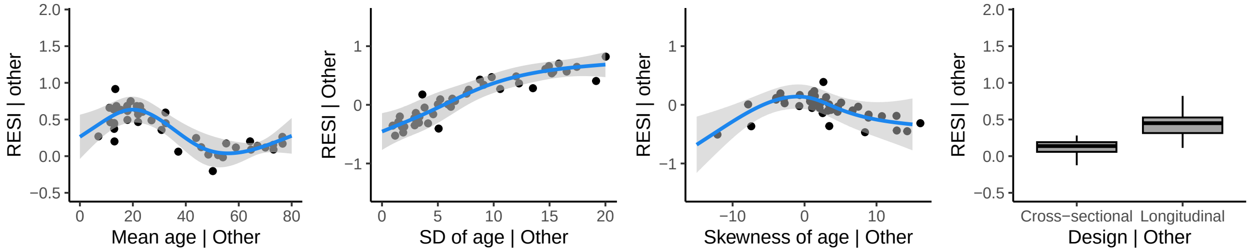


**Postcentral**

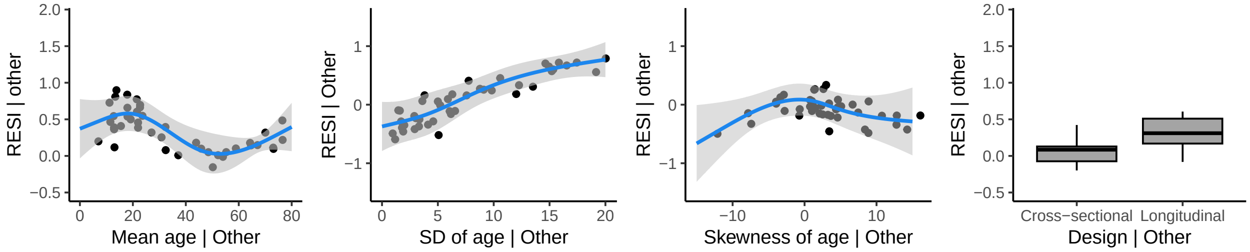

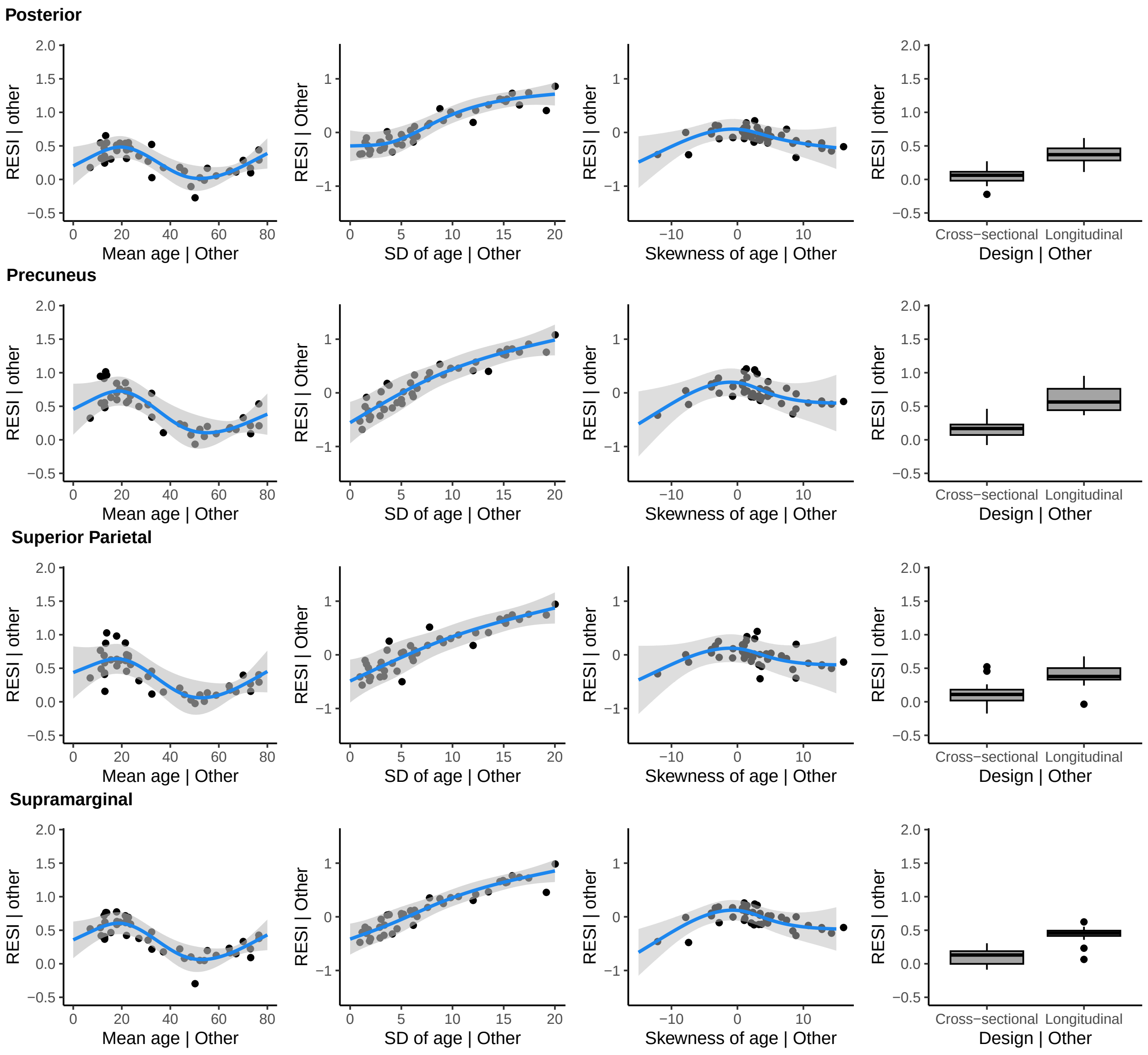

Posterior
Precuneus
Superior Parietal
Supramarginal
RESI | other
RESI | Other
Mean age | Other
SD of age | Other
Skewness of age | Other
Design | Other
Cross−sectional
Longitudinal
2.0
1.5
1.0
0.5
0.0
−0.5
1
0
−1
0
20
40
60
80
5
10
15
−10

### Temporal Lobe (Left hemisphere)

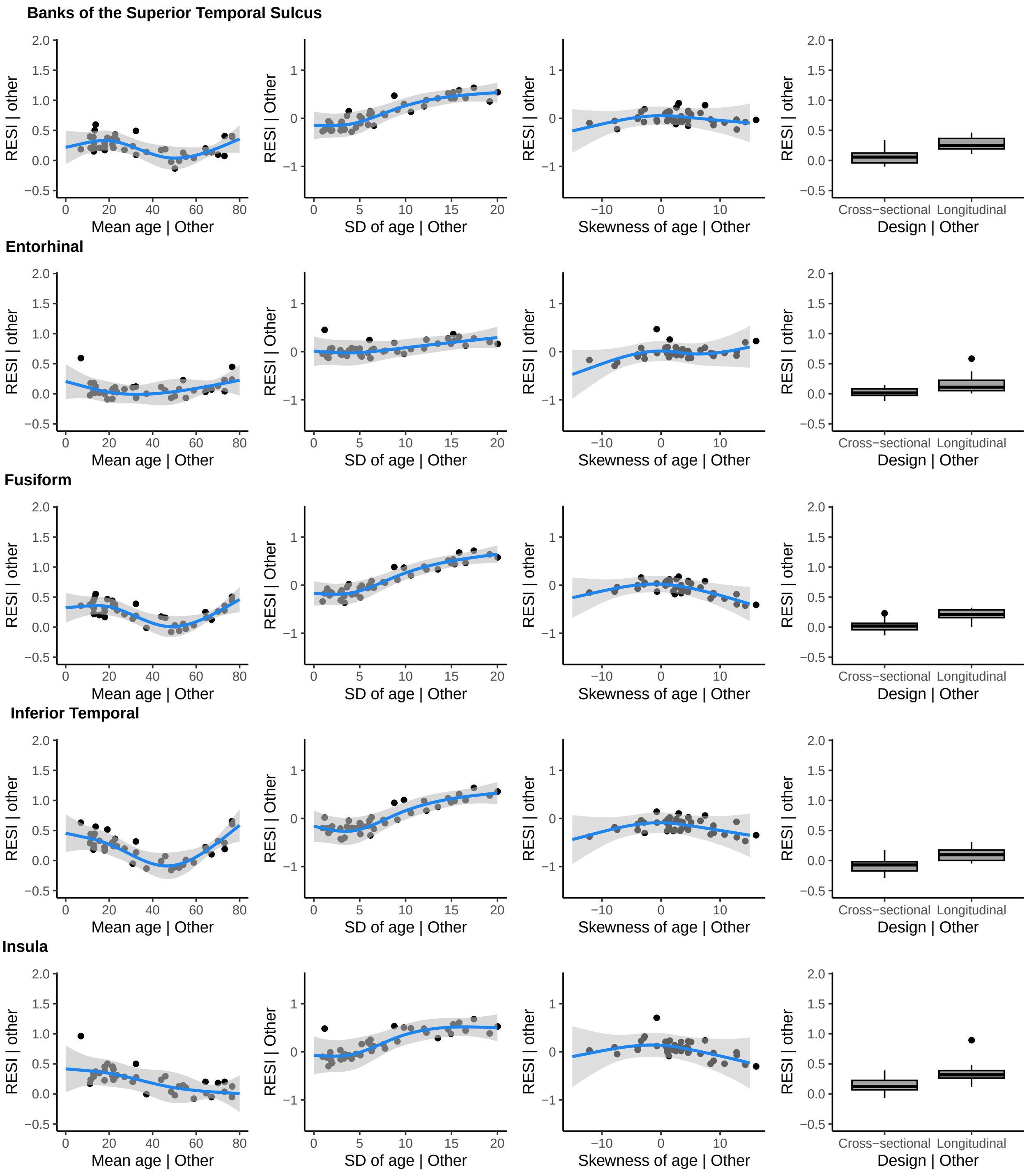

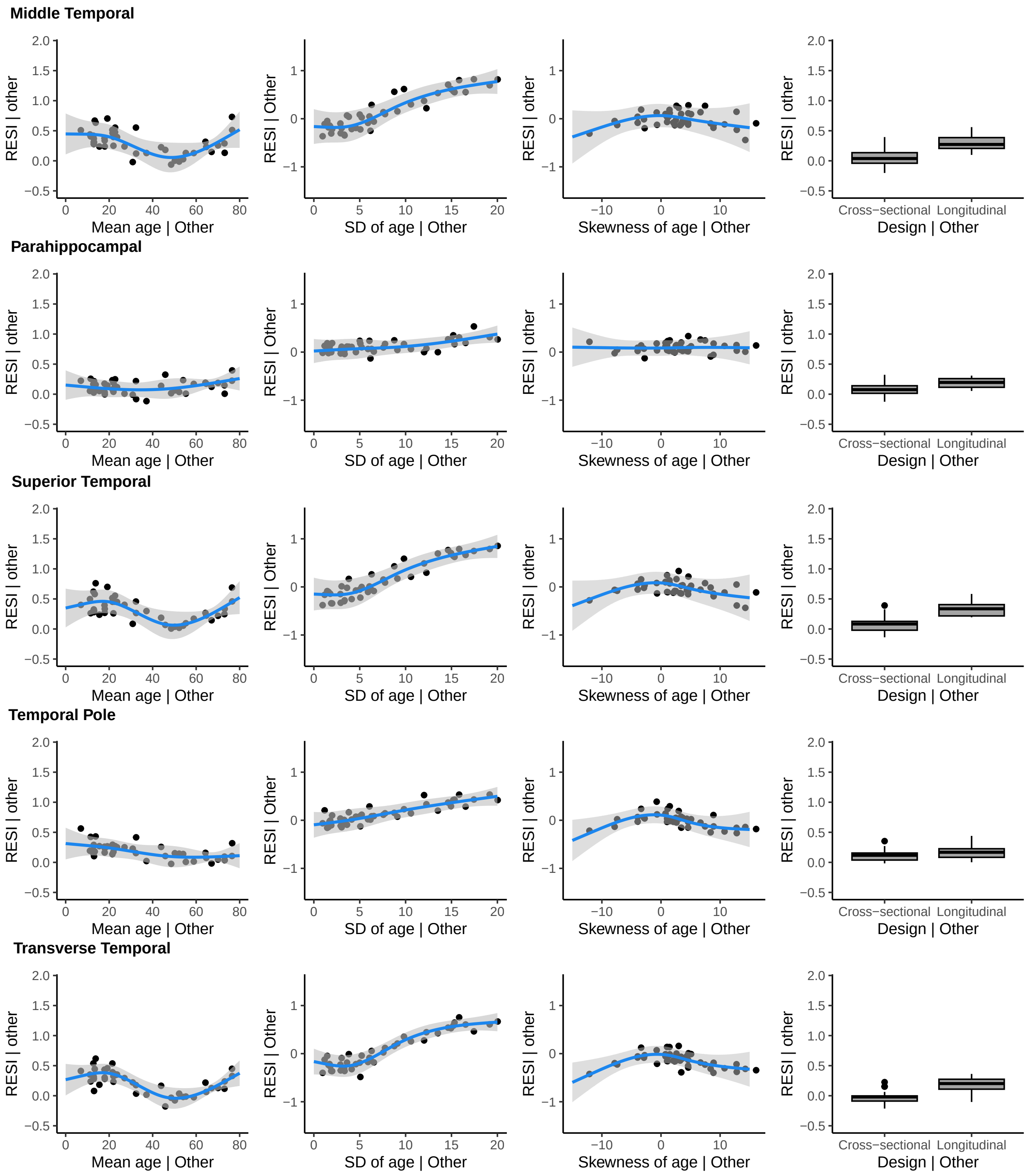
Middle Temporal
Parahippocampal
Superior Temporal
Temporal Pole
Transverse Temporal
RESI | other
RESI | Other
Mean age | Other
SD of age | Other
Skewness of age | Other
Design | Other
Cross−sectional
Longitudinal

### Temporal Lobe (Right hemisphere)

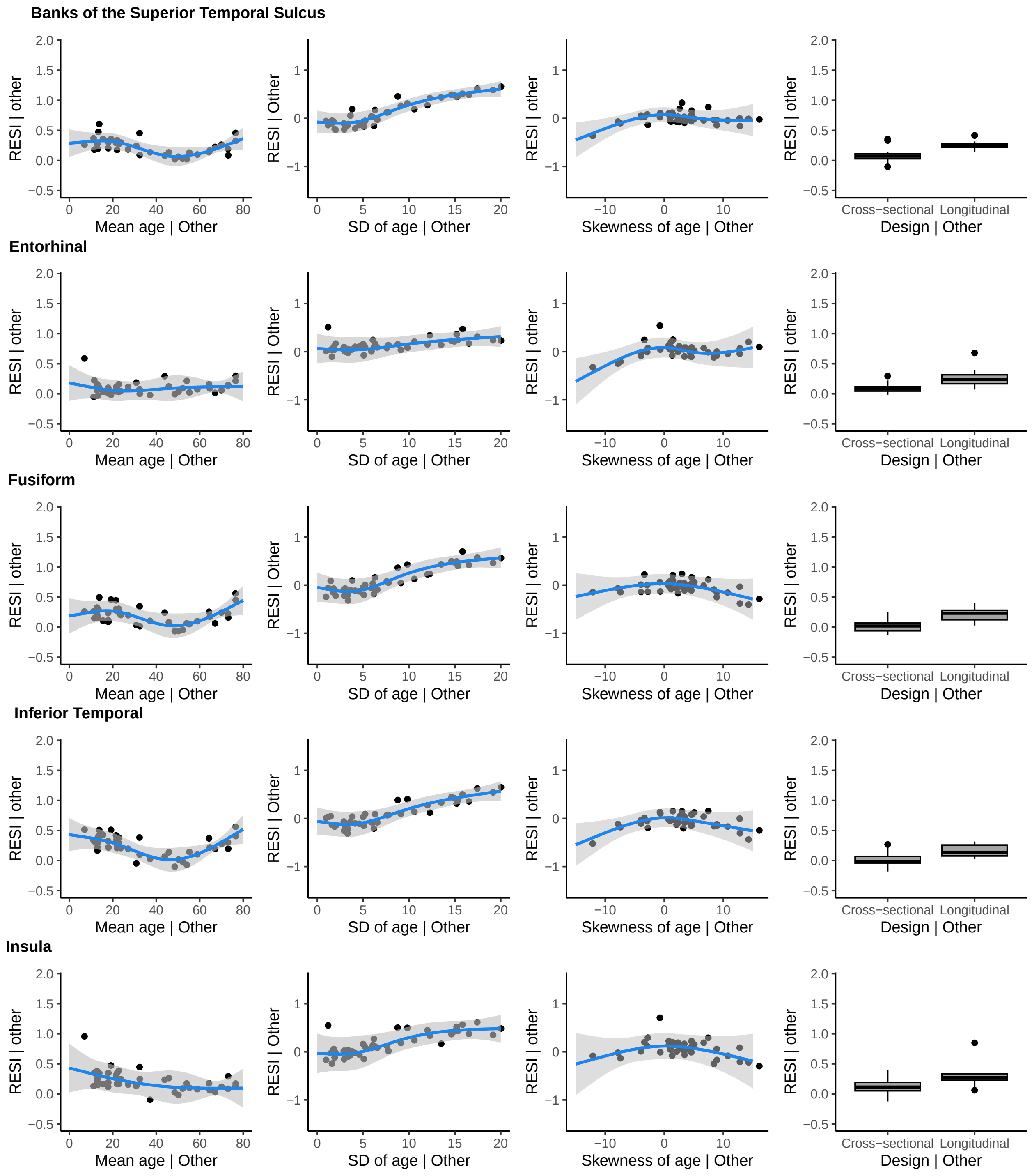

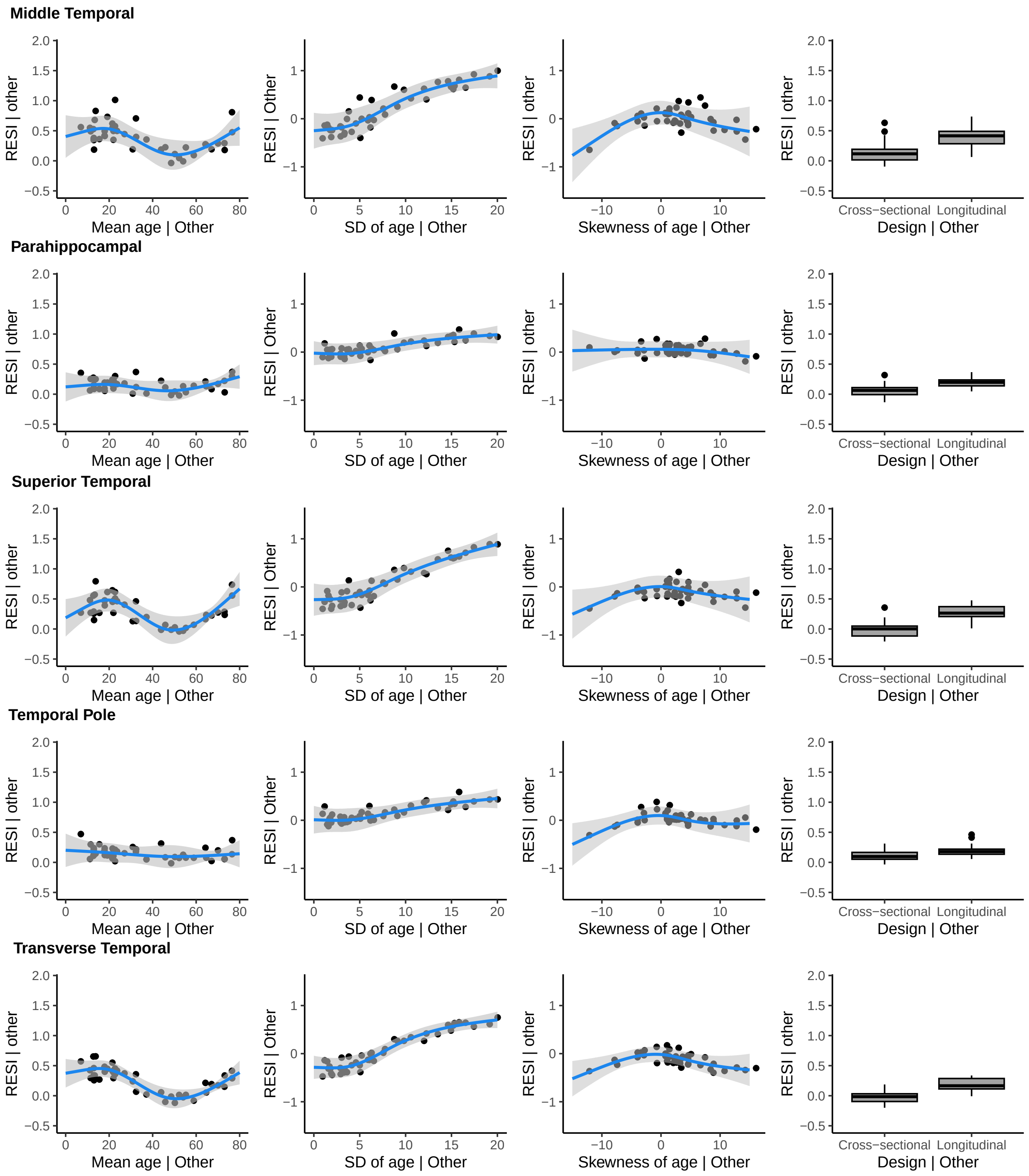
Middle Temporal
Parahippocampal
Superior Temporal
Temporal Pole
Transverse Temporal
RESI | other
RESI | Other
Mean age | Other
SD of age | Other
Skewness of age | Other
Design | Other
Cross−sectional
Longitudinal
